# Forward Entrainment: Evidence, Controversies, Constraints, and Mechanisms

**DOI:** 10.1101/2021.07.06.451373

**Authors:** Kourosh Saberi, Gregory Hickok

## Abstract

We define forward entrainment as that part of the entrainment process that outlasts the entraining stimulus. In this study, we examine conditions under which one may or may not observe forward entrainment. In part 1, we review and evaluate studies that have observed forward entrainment using a variety of psychophysical methods (detection, discrimination and reaction times), different target stimuli (tones, noise, gaps), different entraining sequences (sinusoidal, rectangular or sawtooth waveforms), a variety of physiological measures (MEG, EEG, ECoG, CSD), in different modalities (auditory and visual), across modalities (audiovisual and auditory-motor), and in different species. In part 2, we review those studies that have failed to observe forward entrainment, with emphasis on evaluating the methodological and stimulus design differences that may clarify the contrasting findings across these two classes of studies. In part 3, we describe those experimental conditions under which we ourselves have failed to observe forward entrainment, and provide new data on use of complex envelope patterns as entraining stimuli, show data on intersubject variability, and provide new findings on psychometric functions that characterize the strength of forward entrainment at different SNRs. In part 4 we theorize on potential mechanisms, describe how neurophysiological and psychophysical studies approach the study of entrainment, and caution against drawing direct causal inferences between the two without compelling evidence beyond correlative measures.

## Introduction

An extensive body of literature has investigated neural and psychophysical entrainment to periodic stimuli in different sensory modalities using a variety of experimental methods. These studies have shown that neural activity patterns at several levels of the cortex phase lock to periodic stimuli and that cortical entrainment to the stimulus modulation envelope is both correlated with and predictive of behavioral measures. For reviews see Sameiro-Barbosa and Geiser (2016), VanRullen (2016, 2018), Zoefel and VanRullen (2017), Haegens and Zion Golumbic (2018), Obleser and Kayser (2019), and Bauer et al. (2020). The current study focuses exclusively on a subset of these studies that have shown sustained entrainment (cortical or psychophysical) *after* termination of the driving stimulus. We use the term “forward entrainment” to refer to that part of the entrainment process that outlasts the entraining stimulus, analogous to the concept of forward masking in psychoacoustics where masking effects are observed in signal detection after the masking sound has terminated. We contrast this to “simultaneous entrainment” that describes phenomena which are observed while the entraining stimulus is ongoing. We begin with an overview of studies that have shown forward entrainment (including our own work), we then discuss those studies that have failed to show such entrainment with emphasis on the methodological differences between these two classes of studies, we then describe constraints on experimental conditions that optimize detection of forward entrainment, and conclude with a discussion of how entrainment is evaluated across disciplines (physics, neurophysiology, cognitive science) and speculate on potential mechanisms that underlie forward entrainment.

## 1. Evidence for Entrainment

Forward entrainment typically lasts a fraction of a second and dissipates rapidly by the third or fourth cycle of the expected modulation envelope. In this section, we review those studies that have definitively shown existence of such brief entrainment in different sensory modalities and using a variety of methodological approaches and measurement techniques. We intentionally exclude studies of informational or symbolic cuing (Posner, 1980; Treisman, 1963; Xu et al., 2021; Correa et al., 2004; Coull and Nobre, 1998; Stefanics et al., 2010) and focus instead on those that use implicit cues to capture attention or other involuntary rhythmic-coding (automatic) processes. These studies are summarized in Table 1.

**Table 1.** Selected studies showing (green) or failing to show (red) forward entrainment. The list only includes widely cited studies that have been reviewed in the current paper.

| Study | Paradigm | Entrained Signal | Entraining Sequence | Entrain. Freq. (Hz) |
| --- | --- | --- | --- | --- |
| Lawrance et al. (2014) | Detection | Noise pulses | Noise pulses | 4 |
| Hickok et al. (2015) | Detection | Tone | AM noise | 3 |
| Farahbod et al. (2020) | Detection | Tone | AM noise | 2, 3, 5 |
| Barnes and Jones (2000) | Discrimination | Silent gaps | Temporal intervals | 1.7 |
| Jones et al. (2002) | Discrimination | Tone | Tone sequence | 1.7 |
| de Graaf et al. (2013) | Discrimination (visual) | x or + | Flickering annuli | 5.3, 10.6 |
| Spaak et al. (2014) | Discrimination (visual) | Sinewave grating | Square flashes | 10 |
| Lange (2009) | RT | Gap | Tone sequence | 2 |
| Ellis and Jones (2010) | RT | Tone | Tone sequence | 1, 2, 4 |
| Rimmele et al. (2011) | RT | Tone | Tone sequence | 5 |
| Sanabria and Correa (2013) | RT | Tone | Tone sequence | 1.1, 2.5 |
| Lakatos et al. (2013) | Neural (CSD, MUA) | Deviant tones | Tone sequence | 0.8, 1.6, 3.2, 6.2 |
| Spaak et al. (2014)* | Neural (MEG) | Sinewave grating | Square flashes | 10 |
| Simon and Wallace (2017) | Neural (EEG) | Tone | AM noise | 3 |
| Forseth et al. (2020) | Neural (ECoG) | Tone | AM noise | 3 |
| Bauer et al. (2015) | Pitch Discrimination | Tone | Tone sequence | 1.7 |
| Lin et al. (2021) | FM Discrimination | FM | Tone sequence | 0.8 to 10 |
| Sun et al. (2021) | Tone-in-Noise detection | Tone | AM noise | 3 |

\*Behavioral and Neural

### 1.1. Psychophysical detection and discrimination in forward entrainment

Figure 1 shows data from four different psychophysical studies which have demonstrated forward entrainment.^1^ The top two panels show results from auditory experiments and the bottom two from vision experiments. In each panel, the green shaded region represents the period during which an entraining sequence was active and ongoing. Time zero represent the point at which the forward entrainment period begins. For clarity, we only show the last few cycles of the driving sequence. The red sinusoidal functions are simply a schematic representation of the frequency and phase of the driving modulator (entraining stimulus) and not the actual shape of the sequence envelopes used which ranged from rectangular and sinusoidal acoustic envelopes, to flickering annuli or square patches in visual tasks as described more fully below.

**Figure 1.**
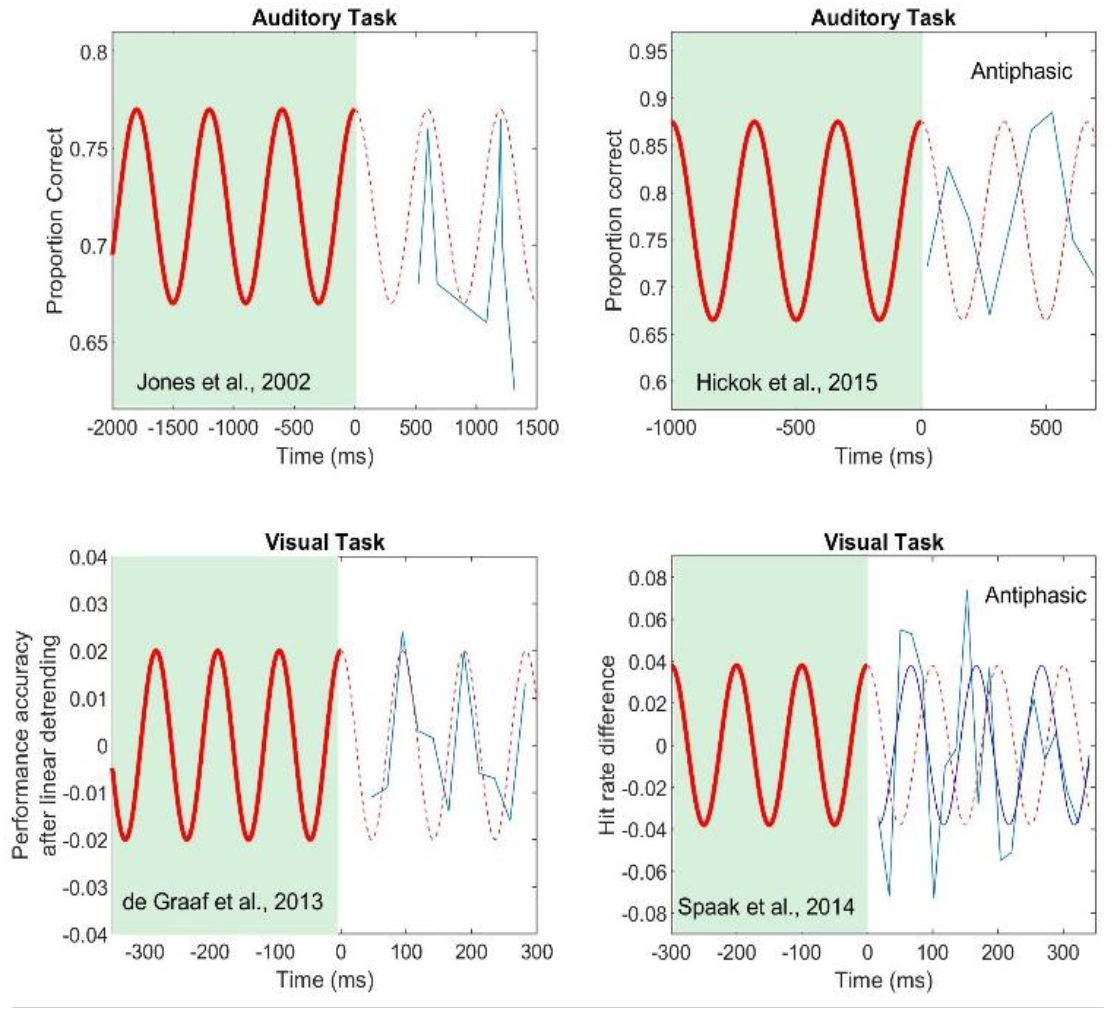
Results from four psychophysical studies that have shown multicycle forward entrainment. Top two panels show results from auditory tasks, and bottom two from vision tasks. Left panels show an in-phase pattern of forward entrainment and right panels show an antiphasic pattern.

Figure 1A displays results from the seminal work of Jones et al. (2002). The results shown are concatenated from two experiments that separately investigated pitch discrimination at different temporal positions after termination of the entraining stimulus (Figs. 3 and 4 of Jones et al., 2002 from 23 subjects). The driving sequence was a binaural (diotic) set of 9 tones with a fixed intertone interval of 600 ms. The first tone in the driving sequence was 150 ms in duration (called the *standard* tone) whose frequency was randomly selected from a closed set of 5 values (415 to 698 Hz corresponding to musical notes A-flat_4_ to F_5_). This was followed by 8 tones that were each 60 ms in duration (random frequency from a closed set) for a total of 9 tones in the driving sequence.^2^ After the final tone in the driving sequence, the *comparison* tone was presented. This comparison tone was also 150 ms in duration and either had the same frequency (pitch) as the standard, or was higher or lower by one semitone. The subject’s task was to indicate whether the pitch of the comparison tone was higher, lower, or the same as that of the standard. The critical variable of interest was the onset time of the comparison tone (called the critical IOI) which occurred either at the expected temporal interval (600 ms) or slightly off (one of 4 shifted onset times 524, 579, 621, and 676 ms). The *a priori* probabilities of the 5 critical IOIs were the same. In a second experiment, the comparison tone was presented at 1200 ms (twice the intertone interval of the driving sequence) to investigate the persistence of oscillatory effects in pitch discrimination. Their results clearly showed a cyclic pattern in pitch discrimination driven by the temporal expectancy set by the driving sequence. They speculated that this effect is based on attentional capture and a purely reflexive adaptive shift of attention in time toward the temporal locus of the target sound. Jones and colleagues have confirmed these general findings in several related or follow-up studies (Jones et al., 2006; Ellis and Jones, 2010; Barnes and Jones, 2000; Barnes and Johnston, 2010).

**Figure 2.**
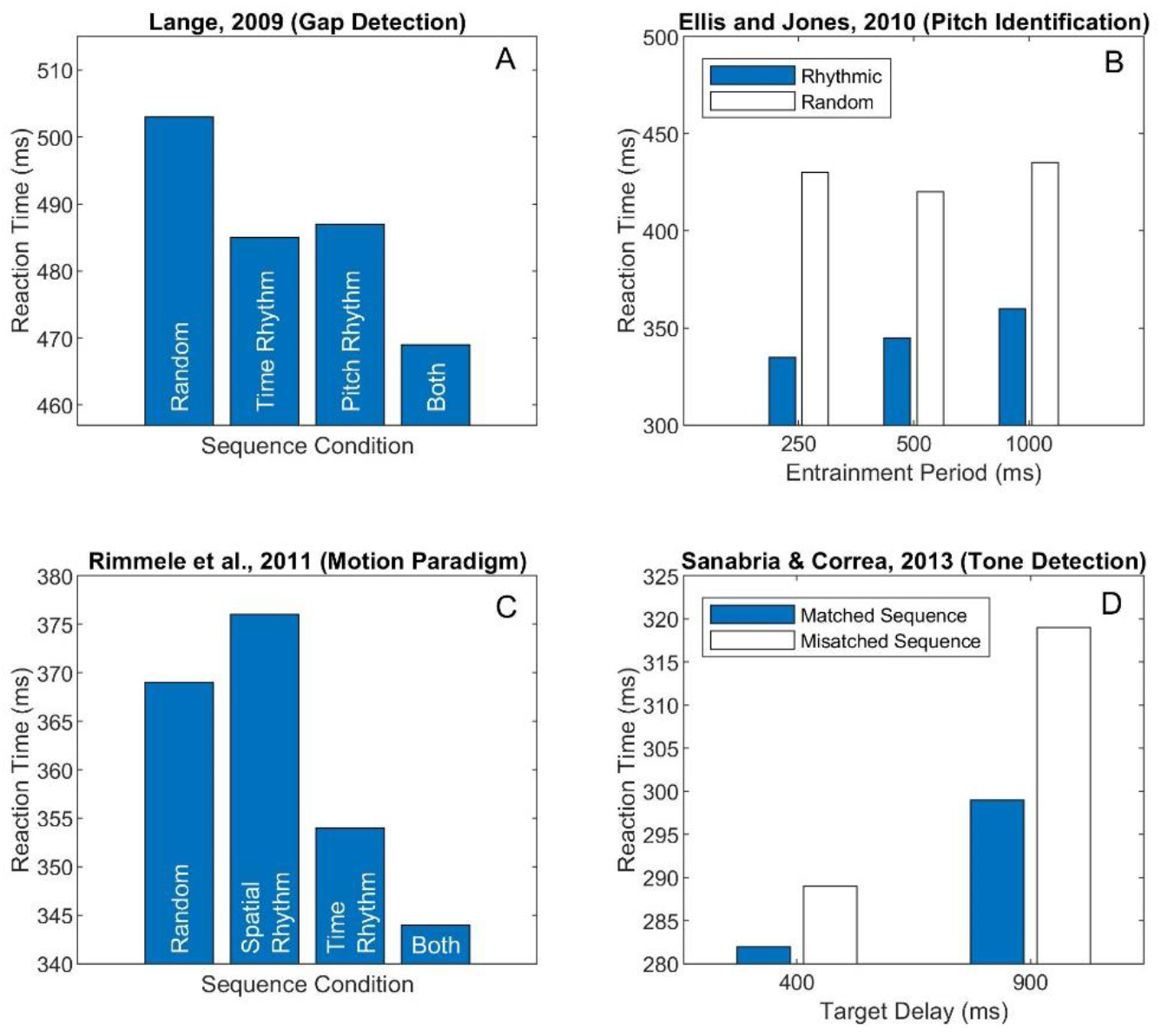
Results from four studies that have shown forward entrainment using reaction-time (RT) measures in four different auditory tasks (gap detection, pitch identification, motion paradigm, and tone detection). See text for detailed explanation of each study.

**Figure 3.**
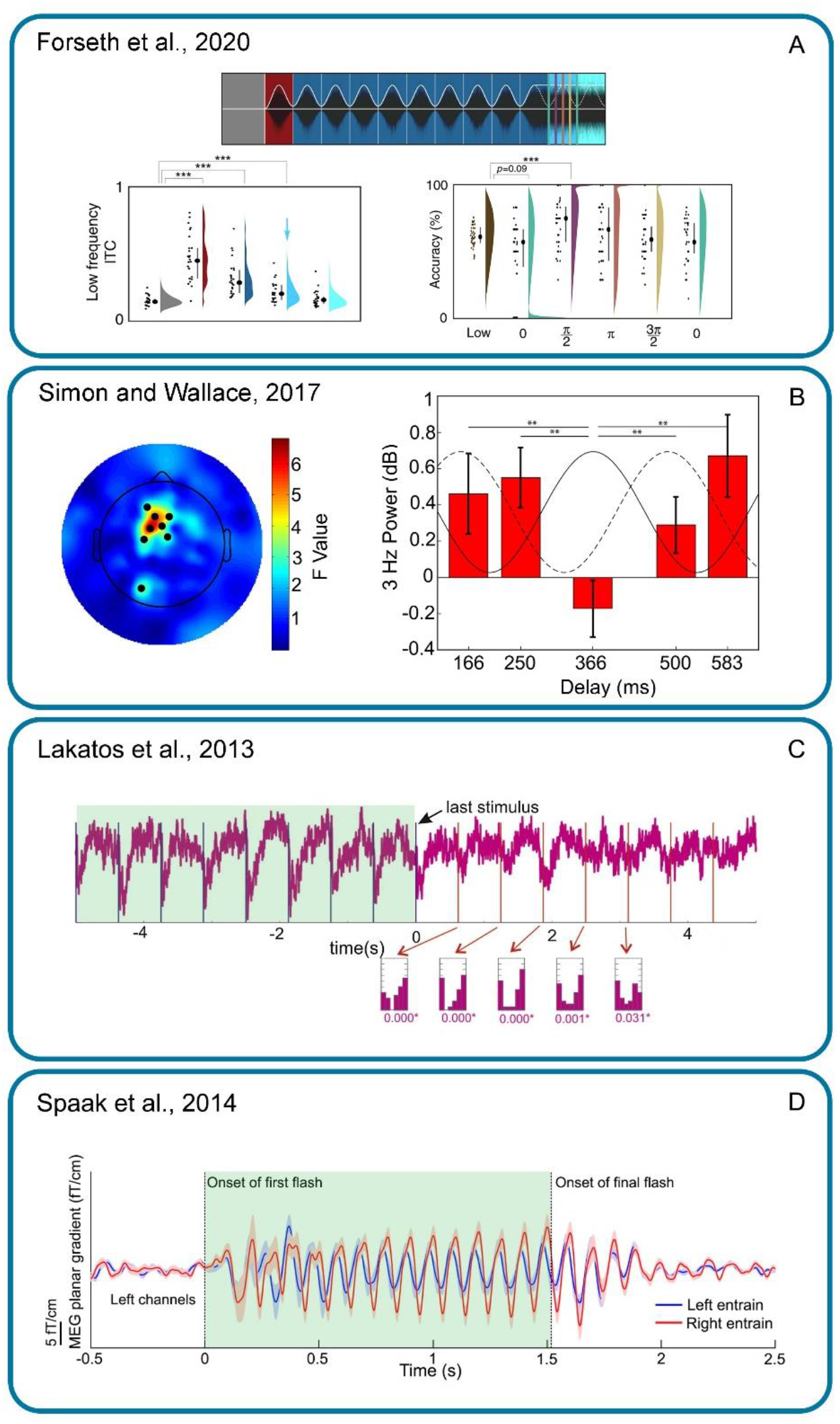
Four neurophysiological studies of forward entrainment using 4 different measurement methods (ECoG, EEG, CSD, and MEG). See text for details. Permission to use granted by Elsevier under STM (The International Association of Scientific, Technical and Medical Publishers) permission guidelines, and by The Journal of Neuroscience and Nature Communications under the terms of the [Creative Commons Attribution 4.0 International License](https://creativecommons.org/licenses/by/4.0/).

**Figure 4.**
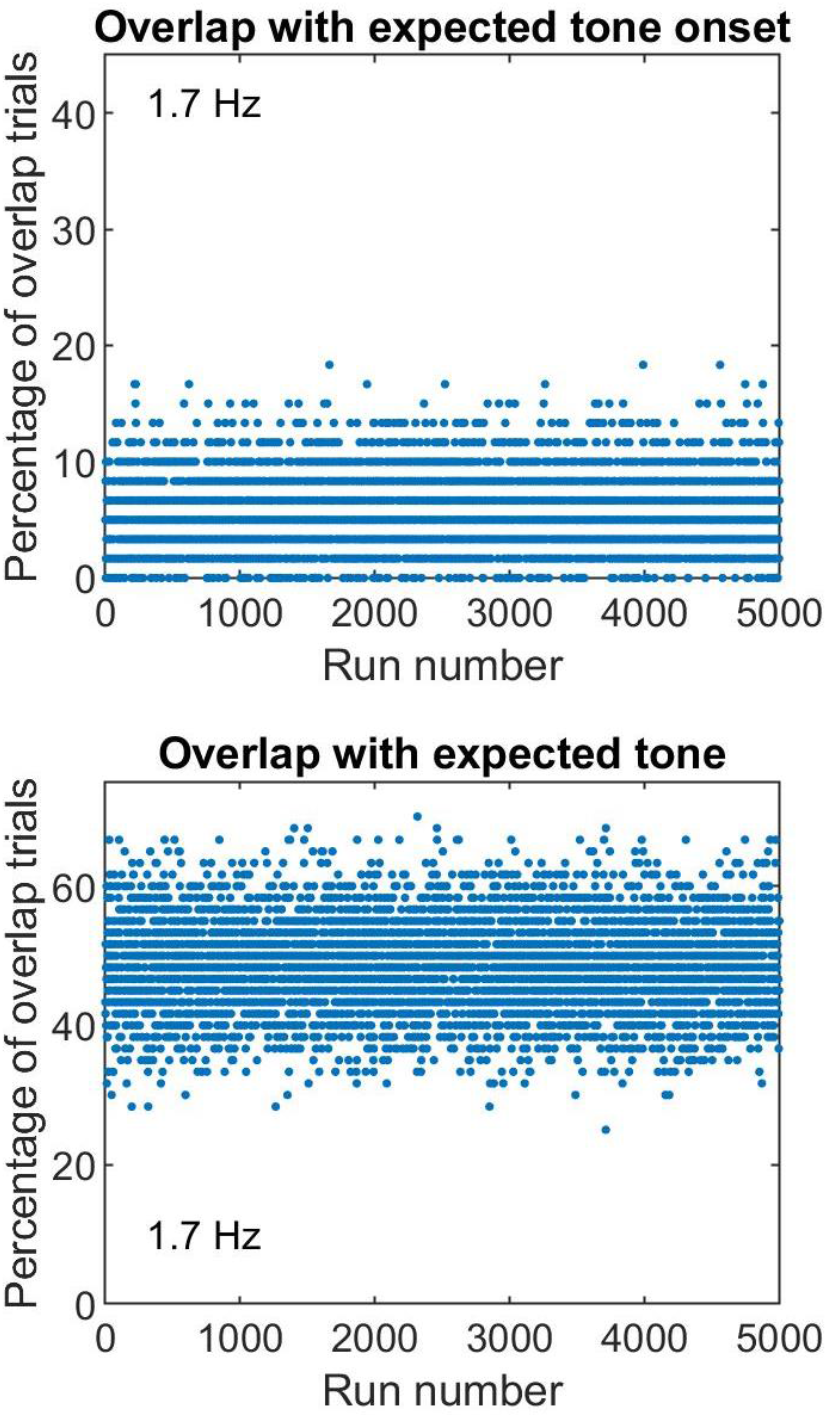
Monte Carlo simulations showing the percentage of trials on which the “random” target occurred during the expected (not random) target window in Lin et al. (2021). This confound diluted the difference in forward entrainment between rhythmic and “random” targets. Top panel shows results for overlap with the *onset* of the temporal expectancy window, and bottom for its full duration.

Figure 1B shows our own findings in an auditory signal-detection study (Hickok et al. 2015). In this experiment, the entraining stimulus was a 3-Hz sinusoidal amplitude modulated (SAM) noise (80% modulation depth) that terminated on the cosine phase of the modulating envelope. The entraining stimulus was then followed immediately by steady state (flat envelope) noise whose amplitude matched the peak of the modulating noise that preceded it. This allowed for a seamless transition between the modulating and steady state noise segments without introduction of acoustic artifacts (see Fig. 1 of Hickok et al., 2015). The signal to be detected was a 50-ms 1-kHz pure tone with a 5-ms rise-decay ramp. On each trial, the tone was randomly presented during the steady state noise at one of 9 temporal positions, corresponding to two full cycles of the driving modulator had it continued (spaced evenly at 0.5π-radian or quarter-cycle intervals). On each trial, the tone’s intensity was randomly selected from one of 5 levels spanning a 12-dB range, sufficient to generate performance levels from near chance to near perfect detection. This level uncertainty appears to be important in observing forward entrainment in *near-threshold* signal detection (Farahbod et al., 2020).

Two findings from this study are immediately apparent. First, there is a cyclic pattern in signal detection that lasts for two cycles after termination of the driving modulator, consistent with findings on pitch discrimination by Jones et al. (2002). Second, and contrary to Jones, we observed an antiphasic pattern in signal detection with best performance near the *expected* troughs of the driving modulator (had it continued), and worst performance at phases corresponding to the expected peaks. In most entrainment studies, the driving stimulus is of the same class as the signal to be detected, e.g., tone sequence and tone signal. In our case, the driving stimulus is noise, i.e., what is to be avoided. To optimize performance, listeners may implicitly adopt a “listening-in-the-dip” strategy where SNR would be most favorable, a phenomenon well established in auditory psychophysics (Hopkins and Moore, 2009; Peters et al., 1998; Festen and Plomp, 1990). Listening in the dip allows subjects to take advantage of “glimpses” in the troughs of the expected modulating masker. In Jones et al. (2002) subjects heard the standard-comparison tones in quiet at suprathreshold levels where attending to the exact in-phase temporal positions would be beneficial (instead of focusing on the gaps between tones). In other words, in our case, the noise is what the subjects are trying to avoid (to extract the tonal signal) hence an antiphasic pattern that entrains against the noise modulation, whereas in the Jones study, there is no noise to avoid, and hence no need to listen at the dips (gaps). This antiphasic pattern, coupled with findings on signal uncertainty (Farahbod et al., 2020) suggests to us that forward entrainment may be largely attention driven (even if implicitly so). Findings from Hickok et al. (2015) have been supported by studies using nearly identical stimuli both psychophysically (Forseth et al., 2020; Farahbod et al., 2020) and neurophysiologically (Simon and Wallace, 2017; Forseth et al., 2020).

The bottom panels of Fig. 1 show findings from two vision studies that have demonstrated forward entrainment in psychophysical signal detection. In the study by de Graaf et al. (2013), subjects were required to detect which of two visual targets (+ or x) briefly (11.8 ms) flashed on the screen. The entrainment sequence preceding the target comprised flashing annuli that flickered at either 5.3 or 10.6 Hz (fixed within a run). The fractional frequency was due to the limitations of their stimulus-presentation system. They found a rhythmic pattern of behavioral performance that lasted for 3 cycles after termination of the entraining stimulus (Fig. 1C, reproduced from Fig. 3A of de Graaf et al., 2013). The oscillatory pattern of performance was restricted to 10 Hz, regardless of whether the entraining frequency was ∼5 or ∼10 Hz, consistent with MEG measures in the same 15 subjects (see section 1.3 below). Figure 1D shows findings from Spaak et al. (2014) reproduced from their Fig. 1C. In the Spaak study, the entraining stimulus comprised brief (17 ms) white square flashes presented rhythmically at a rate of 10 Hz in one visual hemifield, and simultaneously, arrhythmically (jittered) at an *average* rate of 10 Hz to the contralateral hemifield. On each trial, the hemifield that received the rhythmic sequence was randomly selected. The target was a *near-threshold* circular sine-wave grating 1° in diameter and was presented either in the hemifield that carried the rhythmic sequence or the hemifield that carried the arrhythmic (jittered) sequence. The subject’s task was to identify the hemifield within which the target appeared. On each trial the target was presented randomly at one of 20 discrete temporal positions *after* termination of the driving sequence (17 to 340 ms, in 17 ms steps). They found behavioral forward entrainment for 3 cycles after termination of the entraining stimulus.

The blue sinusoid in Fig. 1D represents Spaak et al.’s best-fitting 10-Hz sinusoid to their data. Note that unlike de Graaf, they found that forward entrainment is antiphasic, similar to that reported by Hickok et al. (2015). The key similarity is that both studies have employed *near-threshold* signal intensities: in the Hickok et al. study signal detection was limited by external noise, whereas in the Spaak study it was limited by internal noise (barely visible sine wave grating). Spaak et al. further analyzed the antiphasic nature of their data by directly contrasting performance at target delays that were precisely antiphasic to the driving modulator (100 and 200 ms after end of the entraining stimulus) to only those delays that were in phase (150 and 250 ms). They found a statistically significant difference favoring the antiphasic delays (higher accuracy^3^; see Fig. 1D of Spaak et al., 2014). Similar to de Graaf et al. (2013), the behavioral findings of Spaak et al. were consistent with MEG measurements of alpha cortical activity patterns in the same subjects, demonstrating forward neural entrainment that outlasted the entraining sequence for several cycles. de Graaf et al. (2013) note that they adjusted their target’s salience to approximately 80% performance level, though it is not clear what is meant by salience (intensity, contrast, duration, etc.). Interestingly, although not explicitly noted in their paper, a careful examination of the behavioral performance in the de Graaf study shows an approximately antiphasic pattern when the driving stimulus was 5.3 Hz, i.e., the subharmonic of the observed 10-Hz behavioral oscillation in performance (see their Fig. 3B). The 10-Hz fit to the behavioral data shows lowest performance near where the 10-Hz oscillation peaks would have occurred (i.e., 1^st^ harmonic of the 5 Hz stimulus). The psychophysical results of de Graaf and Spaak are consistent with several other vision studies (Mathewson, 2012; Doherty et al., 2005).

Two other auditory psychophysical studies of forward entrainment are noteworthy. The first is a noise-in-noise detection study by Lawrance et al. (2014) that investigated how a rhythmic noise sequence can preferentially affect the detection of a subsequent near-threshold noise signal.

There are important parallels between this study and Hickok et al. (2015). First, the Lawrance study used signals whose *intensity* was near threshold. Second, the signal (noise) was to be detected in a continuous background noise after termination of the driving sequence. Third, the driving sequence itself was made up of amplitude modulated noise. The entraining sound was a sequence of 7 brief (25ms) rectangular noise bursts superimposed on top of a continuous background noise and presented at a rate of 4 Hz. The intensity of each pulse in the entraining sequence was progressively decreased to generate the percept of a periodic sound that faded into the background noise. This section of the stimulus was followed immediately by a steady state section that either contained or did not contain a signal to be detected. The signal comprised a set of 5 equal-amplitude noise bursts (25ms each). On each trial of a 2IFC task, both a signal and a no-signal stimulus were presented in random order. The subject’s task was to determine which of the two intervals contained the target signal (i.e., the 5 noise bursts). There were two experimental conditions: 1) the *target* was rhythmic (4 Hz), and 2) the target was arrhythmic with interburst intervals randomly selected on each trial from a uniform distribution (2.9 to 6.7 Hz). The two conditions (rhythmic and arrhythmic) were interleaved within the same run so that on a given trial, there was an equal likelihood that a rhythmic or arrhythmic target would occur. Thresholds were measured using an adaptive psychophysical procedure that generated psychometric functions across a range of target intensities (note that here multiple levels are presented within the same run, adding to trial-by-trial uncertainty in level similar to Hickok et al., 2015). Lawrance et al. found that 21 of 26 subjects showed an improvement in detection of rhythmic over arrhythmic targets following the termination of the entrainment sequence. The improvement averaged to approximately 1.5 dB SNR, though across individuals, this advantage could be as high as 3 to 5 dB. Five participants either showed the reverse (higher SNR for arrhythmic targets) or no effect. One difference between results of Lawrance et al. (2014) and Hickok et al. (2015) is that the former found better performance for in-phase targets whereas the latter showed best performance for signals that were antiphasic to the entraining stimulus. Both studies used amplitude modulating noise as the entraining stimulus, and both required subjects to detect a signal in a background of steady state noise after termination of the entrainment segment. There is, however, a fundamental difference between the two studies in the nature of the target signal to be detected. In the Hickok study, the signal was a pure tone. As discussed above, an ideal observer would adopt a strategy to optimize performance by listening at the expected dips of the masker (had the masker modulation continued). This would generate an antiphasic pattern of performance, as observed. In the Lawrance study, however, the signal was a noise burst with identical spectrotemporal and statistical properties as the masker. An implicit strategy to listen in the expected dips of the modulating masker would simply result in “filling in” the gap with a statistically identical noise burst (signal), resulting in a flat noise envelope (and no signal to be detected). It would therefore be advantageous to listen for the noise signal at a point in time where it may be expected. Note that Lawrance et al. did not measure performance for an antiphasic condition, but rather contrasted performance for an in-phase signal condition to a random-phase condition.

The final study we’d like to highlight in this section is by Barnes and Jones (2000). It is an important study in that it observes forward entrainment using an interesting and categorically different type of discrimination task. Barnes and Jones measured the ability of listeners to determine if two temporal intervals (silent gaps) whose edges were marked by brief (60ms) tones were the same or different. Each temporal interval of the entraining sequence was 600ms, i.e., a silent temporal interval bounded by short tone pulses. The last temporal interval of the entraining sequence, however, was selected from one of 5 equally likely values centered on 600ms (ranging from 524 to 676ms). This final “silent” interval was called the *standard* to which a *comparison* “silent” interval was to be compared. Subjects were instructed to ignore the entraining sequence and focus exclusively on judging the difference between the standard and comparison intervals. Two interesting findings emerged. First, they found that performance was better by approximately 20% when the entraining intervals matched the standard, compared to when no entraining sequence was present (see their Fig. 8B). This advantage declined from 20% to 10% when the standard was slightly off (by about 20 ms) relative to the entraining sequence intervals, and to near 0% advantage when the standard and entraining intervals were significantly different (∼75ms). In a very interesting follow-up experiment, they investigated the nature of the internal temporal referent generated by the entraining sequence. The goal was to determine if this referent was based on stored central memories of temporal intervals (i.e., a cognitive effect) or a consequence of an implicit oscillatory resonance. They found that a harmonically related entraining interval (300 ms) more effectively entrained the 600-ms standard, than an entraining interval (500ms) that was closer in duration (but inharmonically related) to the 600-ms standard (Experiment 3, Fig 6 from Barnes and Jones, 2000). This finding supported an oscillatory model of forward entrainment as contrasted to a central memory-storage model.

### 1.2. Reaction time paradigms in forward entrainment

In the previous section we examined forward entrainment in psychophysical detection and discrimination paradigms in which performance was evaluated using an accuracy measure (e.g., proportion correct). In this section, we review studies of forward entrainment that have employed reaction time (RT) as a dependent measure. While RT and accuracy are often correlated (e.g., speed-accuracy trade-off) they are not two sides of the same coin (Kahana and Loftus, 1999; Prinzmetal et al., 2005). One measures the amount of information that a sensory or cognitive process contains (accuracy), and the other, the time it takes to complete the process (RT). Some studies have shown that a variable can have a significant impact on performance accuracy without affecting RTs (MacLeod and Nelson, 1984; Sternberg, 1969) and others have shown significant effects on RT without a correlated change in accuracy that is far from ceiling (Sanders et al., 1974; Santee and Egeth, 1982; Kahana and Loftus, 1999). It is therefore important to explore both accuracy and RT measures in developing theories of psychophysical entrainment. We describe here four auditory psychophysical studies (using four different tasks) that have conclusively demonstrated auditory forward entrainment in a reaction-time paradigm.^4^ Lange (2009) has shown that RTs for detection of a brief gap (10ms) in the middle of a 100ms target sound can significantly improve when the target is preceded by an entraining sequence presented at a rate of ∼2 Hz (this is within the range of rates for which we and others have observed optimum forward entrainment in auditory signal detection and discrimination). The task was to respond as rapidly as possible in a 2IFC task (is there a gap present or not?). The entrainment sequence implicitly cued one of 3 things about the target: 1) when the target may occur, 2) its pitch (via an ascending pitch sequence), or 3) both time *and* pitch. Accuracy (detection performance) was intentionally set to a high level in all cases (average 94%) so that the effects of forward entrainment on RT could be isolated. Results are shown in Fig. 2A. Compared to a control sequence (randomized time/pitch), RTs were significantly lower when either the target’s time or pitch were independently cued. RTs were fastest when the entraining sequence cued *both* the time of occurrence and pitch of the target sound within which the gap occurred. Lange (2009) also reported a neural correlate of these psychophysical findings in both the N1 and P300 components of the ERP signal generated by the rhythmic sequence compared to the arrhythmic (random) sequence.

Ellis and Jones (2010) designed an interesting entrainment sequence that statistically and simultaneously cued for 3 harmonically related entrainment periods: 250, 500, and 1000 ms (4, 2, and 1 Hz). These periods were interleaved within the same entraining sequence creating a nested hierarchical structure of multiple entrainment periods. Each sequence comprised brief tone pulses (10ms) followed by a target tone (25ms) at *one* of the 3 cued periods after termination of the entraining sequence. All tones within a sequence had a fixed frequency of 740 Hz (F#_5_). The target tone had a different frequency that was either higher or lower than that entraining tones by 6 semitones (either 523.3 Hz [C_5_] or 1046.4 Hz [C_6_]). The subject’s task was to determine if the pitch of the target was high or low (a pitch identification task). Subjects had no difficulty with this task as evidenced from high accuracy rates (∼98%). Fig. 2B shows results reproduced from Fig. 5A of Ellis and Jones (2010). RTs for rhythmic sequences were significantly lower than those for scrambled arrhythmic sequences at all 3 entrainment periods, even though these three entraining periods were interleaved within the same sequence.

**Figure 5.**
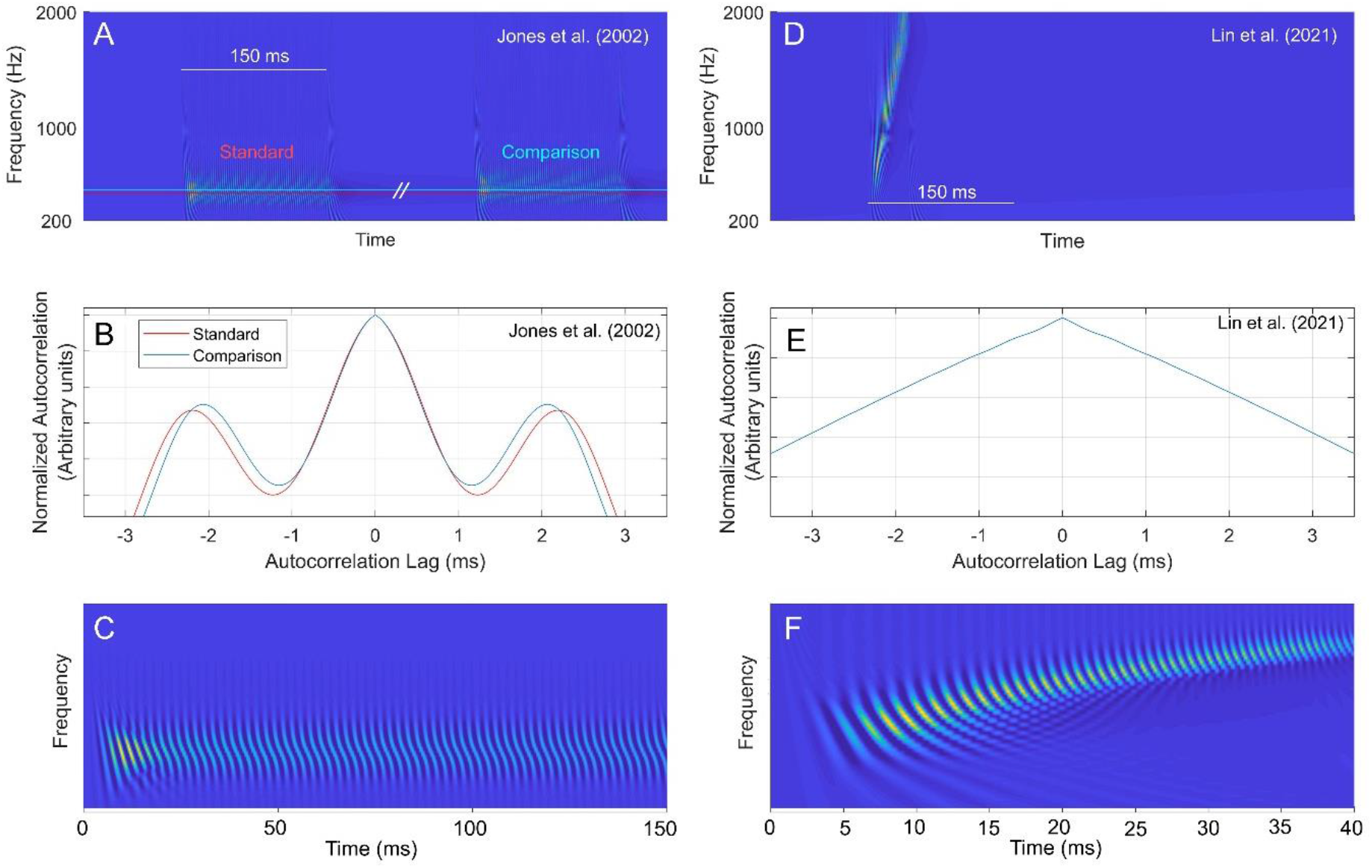
Output of an autocorrelation model of pitch extraction for the type of stimuli used by Jones et al., 2002 (left panels) and Lin et al., 2021 (right panels). No pitch estimate (secondary peak) is obtained from processing Lin et al.’s stimuli through the model. See text for details.

Rimmele et al. (2011) investigated both RTs and detection sensitivity (*dꞌ*) in a study that combined temporal and spatial regularity by use of auditory motion stimuli. Four possible entraining sequence permutations included motion stimuli that either had temporal regularity only (rhythmic sequence), spatial regularity only, temporal *and* spatial regularity, or no regularity (random phase). At the end of the entrainment sequence, a brief (occluding) noise was presented, followed by either a target signal (tone) or no signal. The subject’s task was to respond as quickly as possible if they detected a signal (go no-go task). Figure 2C shows their results. They found lower RTs (by about 25ms) and slightly higher *dꞌ*s for targets preceded by a temporal entrainment sequence, but not for targets preceded by spatial regularity. Their results were consistent with ERP measures where the P1, N1, and N2 components (reflecting pre-motor response, early perceptual processing, and task related responses) were modulated by the rhythmic entrainment cues, but not by spatial entrainment. Spatial regularity cues, however, did modulate later response-related processing stages of the ERP waveform (P3) but only when temporal regularity was concurrently present in the entrainment sequence.

Sanabria and Correa (2013) reported similar findings in a tone-detection task. They showed that when a 50-ms target tone is preceded by a rhythmic sequence of 50-ms tones, RTs are lower when the target ISI matched the rhythmic cadence of the entraining sequence. They used two entraining ISIs of 400 or 900ms (rates of 2.5 and 1.1 Hz), and two target ISIs (400 or 900ms) randomly mixed within the same block of trials (4 possible permutations). The subject’s task was to respond as quickly as possible when they heard the target tone which had a different frequency (400 Hz) than the sequence tones. On 17% of the trials, no target signal was presented (catch trials to estimate false alarm rates). Figure 2D shows results from Sanabria and Correa (2013). They found that RTs were lower when the sequence and target ISIs were matched, though only the slower entrainment rate (900 ms) resulted in a statistically significant effect.

This behavioral improvement in RT was accompanied by correlated modulations of the N1 and P2 potentials of ERP recordings in the same subjects. Interestingly, there was a significant difference in the N1 component in response to the 400ms target ISI, for which a statistically significant behavioral improvement had not been observed. This suggests that the underlying neural process may not have triggered a sufficiently strong behavioral response to be observed in psychophysical measurements.

One interesting observation about these four RT studies is the absence of an antiphasic pattern of performance. Improvements in RT occur when the signal is in-phase with the temporal expectancies set by the entraining sequence. RT measures, however, are made in quiet at suprathreshold signal intensities (near ceiling) with a sequence that is of the same class as the signal type (tones). No prior RT study has used near-threshold signals in noise where avoiding the temporal expectancies set by the peak of the noise modulator would be the optimum strategy in isolating the tonal signal.

What do RT studies of forward entrainment tell us beyond findings from detection and discrimination paradigms? There is clearly a correlation between the two classes of studies in that entraining sequences can improve accuracy (amount of information) and reduce RTs (time it takes to accumulate sufficient information for a decision). The magnitude of improvements in RT resulting from forward entrainment is generally small, under 100 ms (Ellis and Jones, 2010) and more often under 40 ms (Lange, 2009; Rimmele et al., 2011; Sanabria and Correa, 2013). The magnitude of improvements in performance accuracy for entrained auditory signals can be as large as 10 to 20% (Jones et al., 2002; Hickok et al., 2015, Farahbod et al., 2020). How do we compare these two measures? There have been a number of attempts to relate RT to accuracy using detection-theoretic modeling approaches (Ratcliff, 1978, 2018; Kahana and Loftus, 1999; Wagenmakers, 2007; Laming, 1968, 1986; Kornblum, 1973). These, however, require RT measurements at several different SNRs (or detection levels). Currently, no study has measured entrainment effects simultaneously on RT and accuracy at different performance levels. Typically, studies using RT as a dependent measure maintain accuracy performance at a single value near ceiling level. It would be useful to investigate forward entrainment using joint RT and accuracy measures, as it is likely that these two measures are evaluating the same underlying dimension of information from different perspectives.

### 1.3 Neurophysiological findings in forward entrainment

Several studies have observed neural forward entrainment using a variety of measurement techniques from ECoG, EEG, and MEG in humans to single or multiunit electrode recordings in animals. Some of these studies have concurrently measured psychophysical performance that appears correlated with the modulating neural activity patterns. In this section we focus on four of these studies, a recent ECoG study of awake human subjects performing an auditory signal-detection task while their brain activity was directly recorded using intracranial (depth and/or grid) electrodes (Forseth et al., 2020), an EEG study showing antiphasic auditory forward entrainment in fronto-central brain regions while attending to poststimulus auditory targets but not when attending to audiovisual targets (Simon and Wallace, 2017), an MEG study that reported forward entrainment of neural activity in occipitoposterior areas of the cortex in response to rhythmic visual stimuli (Spaak, 2014), and an animal neurophysiological study that used microelectrodes to directly record from the awake monkey primary auditory cortex and found forward entrainment that was critically dependent on attention to the sound sequence (Lakatos et al., 2013). Two of these studies (Forseth et al., 2020, and Simon and Wallace, 2017) employed stimuli that were identical to those used by Hickok et al. (2015). We describe each of these studies in more detail below.

Forseth et al. (2020) measured oscillatory entrainment of cortical activity using ECoG (electrocorticography) in 37 patients. The patients were surgically fitted with depth or surface-grid electrodes (or both) and cortical recordings were made while the patient was awake and performing the same signal-detection task used by Hickok et al. in which an entraining modulating noise (3 Hz, 80% depth) was followed by steady state (flat envelope) noise during which subjects were to detect a brief (50 ms) pure-tone signal (1kHz). Figure 3A shows their results reproduced from their Figs. 4A, C and D. Two of their findings are especially relevant here. First, they found that modulation of neural activity in the early auditory cortex (Heschl’s gyrus [HG] and the transverse temporal sulcus [TTS]) continues phase-locked to the driving modulator, as measured by intertrial phase coherence (ITC), for one cycle after termination of stimulus modulation (mid-blue color in left panel of Fig. 3A marked by arrow). Second, they found that modulation of behavioral performance (near threshold signal-detection) measured simultaneously during neural recordings outlasted the rhythmic stimulus for one cycle after termination of stimulus modulation (they only examined one cycle post modulation). Psychophysical measurements were made for signal (tone) temporal positions that started at the negative cosine phase of the first expected modulation cycle (the first expected dip), and spaced at quarter-cycle intervals thereafter for one full cycle. Psychophysical results are shown in the right panel of Fig. 3A where performance at π/2 is significantly higher than baseline. These findings are consistent with Hickok et al. (2015) and Farahbod et al. (2020) though the exact phase at which best performance is observed is slightly different (by a quarter of a cycle). Psychophysical performance in the Forseth et al. study shows a peak at the rising slope (acoustic edge) of the expected modulating envelope (had it continued), whereas Hickok et al. (2015) see best performance at delays that are antiphasic to the expected modulating envelope.

Simon and Wallace (2017) also used stimuli identical to those used by Hickok et al. (2015) with the exception that after the entraining 3-Hz modulated noise terminated, the tone pulse was presented in *quiet* to isolate EEG measurements that exclusively reflected forward entrainment. The target tone (50ms, 1kHz) was presented at one of five temporal positions (166, 250, 366, 500, or 583 ms after the end of the modulating noise). On a small proportion of trials, the target tone had a “deviant” frequency (333 Hz) and the subject’s task was to press a key when the deviant tone was detected (oddball task) and do nothing otherwise. The deviant tone always occurred at the 366-ms delay. This was an easy task as evidenced by hit rates of greater than 95%. Analysis of EEG recordings focused on the FCz (fronto-central) and neighboring electrodes as this ROI had been previously shown to be associated with typical scalp distributions of auditory generators (Lenz et al., 2007). The main findings from Simon and Wallace (2017) is shown in Fig. 3B (reproduced from their Fig. 6D). Time zero represents the end of the entraining stimulus, and the abscissa represents the delay between the end of the entrainment and the target tone. The solid black curve indicates the phase of the entraining stimulus (had it continued), and the dotted curve indicates what they refer to as brain phase (stimulus phase + a brain lag of 120 ms). They found that the magnitude of the 3-Hz power (the entraining frequency) showed significant phase dependency that was consistent with a 3-Hz oscillatory cycle. The power modulation at the fronto-central sites was antiphasic to the terminated driving modulator, similar to that reported for behavioral measurements by Hickok et al. (2015). This means that the forward oscillations induced by the entraining noise (after it had terminated) was substantially attenuated when the 50-ms target pulse coincided with when the rhythmic noise would have peaked had it continued. They speculated that “when the entrainment stimulus is irrelevant noise, the brain entrains *against* the noise and thus aims to enhance the processing of salient events happening during the gaps”, i.e., a listening in the dips strategy.^5^ One important difference between the Simon and Wallace (2017) and Forseth et al. (2017) study, other than measurement methodology, is that the former made measurements over the fronto-central regions of the brain (which could reflect a broader range of processing mechanisms including decision or attentional processes) while the latter measured activity directly in the primary auditory cortex.

**Figure 6.**
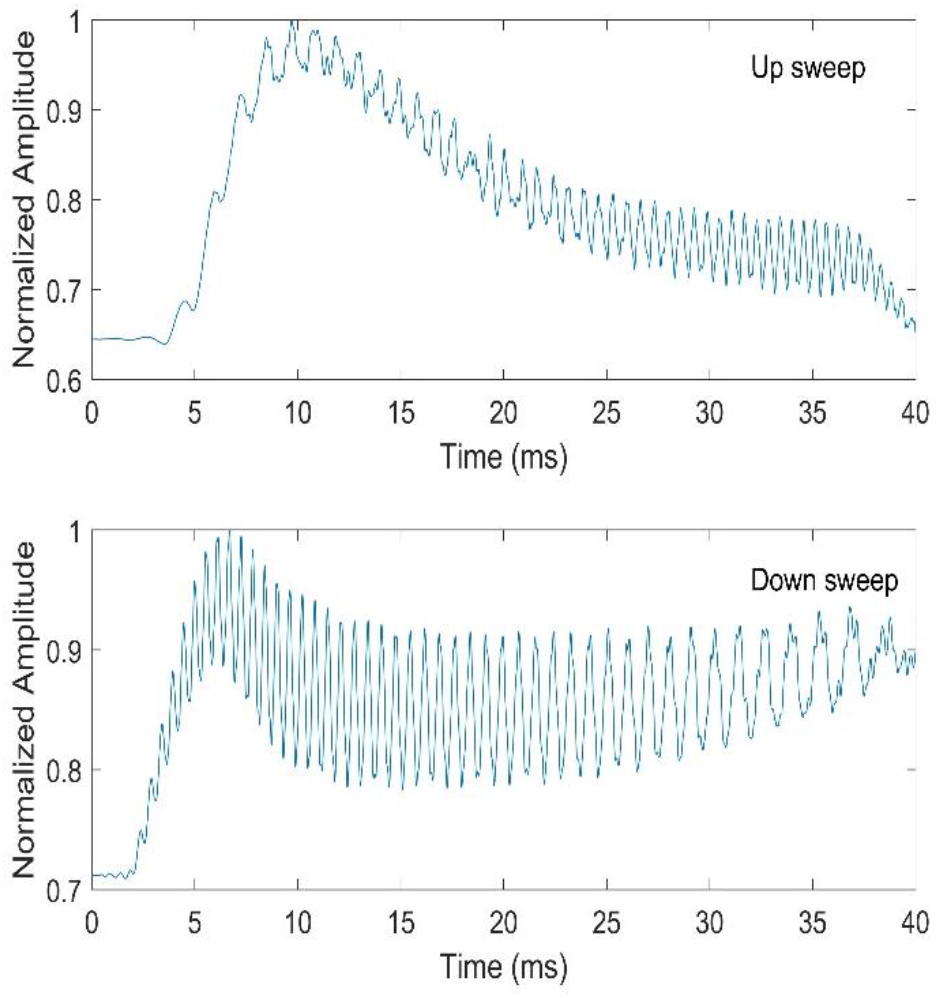
Temporally asymmetric response of an auditory filter model to up and down sweeps. This envelope and fine structure asymmetry contributes to dynamic timbre differences.

Lakatos et al. (2013) measured current source densities (CSD) and multiunit activity (MUA) from the primary auditory cortex (A1) of awake macaque monkeys as they were attending to sequences of pure-tone pulses (25ms) presented at a low rates (from 1.6 to 12.2 Hz). The monkeys were trained to respond to deviant tones (2 to 4 semitone difference) which occurred every 3 to 9 seconds (oddball design). They found that after the termination of the driving sequence, neural activity continued to oscillate rhythmically (and in phase) with the discontinued driving sequence for 5 seconds *after* the end of the sequence. This effect was critically dependent on attention and was absent for non-attended sequences. Figure 3C (reproduced from Fig. 3A of Lakatos et al. 2013) shows 10 seconds of averaged CSD activity in response to the 1.6 Hz tone sequence (5s of stimulation and 5s after the end of the entraining tone sequence). Time zero represents the end of the entraining stimulus (shaded green rectangle). The blue drop lines represent times at which the 25ms tone pulses occurred, and the red drop lines show time points at which these pulses would have occurred if the entraining stimulus had continued (note that negative CSD designates high excitability). The histograms below the CSD trace show the distribution of phases at these time points. Below the histograms, significant *p* values from Rayleigh tests are displayed for each phase distribution. The sustained oscillating neural activity (forward entrainment) was observed for rhythm rates from 0.8 to 6.2 Hz, but not for the higher rate of 12.2 Hz. The range of rates for which neural forward entrainment was observed is approximately the same as those for which our lab has reported behavioral forward entrainment, i.e., 2 to 5 Hz but not at higher rates (Farahbod et al., 2020).^6^

Finally, forward neural entrainment has also been shown in other sensory domains. We highlight one of them here. Spaak et al. (2014) used an entraining stimulus comprised of a rhythmic sequence of visual flashes (10 Hz) while recording MEG signals from sensors overlying the occipitoposterior areas of the cortex. Figure 3D shows an example recording. The red and blue curves correspond to rhythmic stimulation in the right and left visual fields, respectively. The green shaded region (added by us) shows the time period during which the visual flash sequence was presented. Note that after termination of the entrainment sequence the cyclic MEG planar gradient persists for several cycles. One interesting difference between this study in the visual modality and the auditory neurophysiological study by Lakatos is worth noting. Lakatos et al. suggest that the observed ongoing oscillatory neural activity is critically dependent on attention, and that it is not observed when the animal is not attending to the target sequence. This suggestion is consistent with our interpretation of psychophysical forward entrainment (Farahbod et al., 2020). Spaak et al., however, suggest that the ongoing oscillatory pattern of neural (and behavioral) activity they have observed is likely a low-level process that taps into the kinetics of the neural system (similar to the resonance of a system) and not a process that has necessarily evolved to extract temporal information. As evidence for this interpretation, they note that the oscillatory pattern of behavioral performance they observed is antiphasic to the driving entrainment stimulus (see section 1.1 above). They argue that temporal expectancy would predict the opposite pattern and enhanced performance at in-phase delays. This is precisely the argument initially made by Hickok et al. (2015), but additional experiments that investigated the role of stimulus uncertainty (Farahbod et al., 2020) suggested that the antiphasic pattern of behavioral performance is, in fact, consistent with an attentional process that promotes a “listening-in-the-dip” strategy. We should note that Spaak et al. (2014) do not dismiss the role of selective attention given the substantial evidence showing that neural oscillations are strongly affected by top-down attentional control (Haegens et al., 2011; Händel et al., 2011; Bonnefond and Jensen, 2012) and in fact suggest that it would be of significant interest to investigate how attention may interact with entrained oscillations they have observed in the visual cortex.

In addition to the four studies using four different neurophysiological methods highlighted here, there are a number of other similar studies in the auditory and visual domains that have demonstrated forward entrainment of neural activity in the cortex in response to rhythmic stimuli (Lange, 2009; Schmidt-Kassow et al., 2009; Rimmele et al., 2011; Sanabria and Correa, 2013; de Graaf, 2013; Kösem et al., 2018). Most of these studies have also simultaneously shown behavioral correlates that appear phase-locked to the terminated rhythmic stimulus. In summary, forward neural entrainment has been shown using a variety of recording methods (EEG, MEG, MUA, CSD, and ECoG), in humans and animals at multiple levels of the cortex, and across sensory modalities (auditory and visual domains).

## 2. Controversies

In this section we focus on three psychophysical studies that have reported an absence of poststimulus entrainment to a terminated rhythmic stimulus. The first two studies (Bauer et al., 2015 and Lin et al., 2021) have (largely) failed to observe forward entrainment in a *pitch-discrimination* task. These studies were motivated by the original work of Jones et al. (2002) and several follow-up studies from their lab that demonstrated periodic patterns of psychophysical performance after the end of a driving rhythmic sequence (as reviewed above in section 1.1). In particular, we examine the psychophysical and methodological design differences between these studies. The third study we discuss here is a recent signal-detection study by Sun et al. (2021) who investigated forward entrainment using stimuli similar to those used by Hickok et al. (2015). Our goal here is to better understand potential methodological and experimental design differences that may explain different findings across studies.

### 2.1 Bauer et al. (2015) pitch-discrimination study

The Bauer et al. (2015) study is important as it has been widely cited in the literature, including in recent reviews, as an example of the absence of forward entrainment (Haegens and Zion Golumbic, 2018; Shalev et al., 2019; Bauer et al., 2020). Bauer et al. describe 6 experiments in which they (mostly) failed to observe poststimulus entrainment to a rhythmic stimulus in a pitch-discrimination task designed to replicate Jones et al. (2002). Five of these experiments are “conceptual replications” whose experimental and stimulus design features were fundamentally different than those used by Jones et al. (2002). Here we examine only one of their experiments which they called an “exact replication” although we note several differences below. The details of the stimuli and experimental procedures are described above in section 1.1 where we discuss the original Jones et al. study. Bauer et al. found no advantage in pitch-discrimination when the timing of standard and comparison tones were cued by a rhythmic sequence. For the purpose of this review, we will focus on methodological, stimulus design, and subject population differences that may underlie their discrepant findings.

First, subjects in the Bauer et al. (2015) study performed considerably poorer overall compared to subjects used by Jones et al. (2002). In the Bauer study, performance across 30 subjects averaged to 51% correct response level (chance was 33% in the 3AFC design) while subjects in the “methodologically equivalent” Jones et al. study performed at 72% correct level, a mean difference of 21%. This difference is significant with potentially important implications. Subjects in the Jones study were effectively discriminating pitch along a much steeper slope of the psychometric function (where discrimination performance would be optimal) near what is referred to as the sweetpoint of the psychometric function (Green, 1990, 1993, 1995), whereas in the Bauer et al. study subjects were, on average, discriminating pitch along the nonoptimal shallow region of the psychometric function. The 21% difference in correct discrimination level in a 3AFC task translates to an approximately 0.75 *dꞌ* unit difference in sensitivity from 0.59 to 1.32 where threshold is often conventionally set at *dꞌ=1* (Green and Swets, 1966; Macmillan and Creelman, 2005). This increased sensitivity from below to above threshold levels could have led to better encoding of cued-pitch information given that performance is less adversely affected by internal noise.

Second, the subject populations in the two studies were likely quite different. Jones et al. excluded from analysis six subjects that either had extremely high performance levels, or who in a post-experiment questionnaire reported recognizing that the standard tone (pitch) was repeated at the end of the entrainment sequence. This recognition made the rhythmic sequence less useful since subjects only needed to attend to the repeated standard tone at the end of the sequence and ignore the preceding tones in the sequence. Note that the entrainment sequence was actually a distractor to be ignored. The inability to ignore the distractor sequence reduces the overall performance level, though it would be expected to enhance the *difference* in performance at the 5 temporal positions of the target (comparison) tone. Jones et al. also excluded any subject who generated a disproportionately large number of “same” responses (overall, 27 of 122 total subjects were eliminated from the three experiments in Jones et al.). No comparable control or subject-selection policy was employed by Bauer et al. (2015).

Third, there are several differences between the two studies related to stimulus intensities. Bauer et al. normalized loudness in their tone sequences. The level of each tone in their entraining sequence was adjusted to equal loudness according to ANSI S3.4-2007 standards. This departure from Jones et al. (2002) means that the entrainment sequence tones used by Bauer were significantly more intense than the standard or comparison tones since the sequence tones were about one third the duration of the standard/comparison (60 vs. 150 ms). To match for loudness, the intensity of the sequence tones had to be increased by as much as 6 dB (Stevens and Hall, 1966). Tones of different *frequencies* also produce different loudness contours, with higher frequency tones at a fixed intensity associated with greater loudness for the range of pure-tone frequencies used here (Suzuki and Takeshima, 2004). Thus, in addition to duration differences generating intensity confounds when matched for loudness, sound levels were further altered (even between tones within the same sequence) by virtue of their different pitches when matched for loudness. Separate from loudness matching, *overall* levels may also have been a source of methodological difference between the two studies. While neither provides information on precise levels, both state that stimuli were presented at comfortable listening levels (typically 50 to 75 dB SPL). Within this range, more intense sounds, even at comfortable listening levels, may lead to saturation in subpopulations of low-threshold neurons that code for pitch (i.e., rate-intensity functions), resulting in diminished frequency resolution (Kiang et al., 1965; Liberman, 1978; Ohlemiller et al., 1991). Furthermore, the pitch of a tone is a nonlinear function of intensity; for a tone of fixed frequency, a change in intensity could change pitch by as much as the equivalent of a semitone (Stevens, 1935; Cohen, 1961). Thus, aside from differences attributed to equal-loudness adjustments, it is unclear what effect overall intensity may have had on performance differences given that this information is not provided by either study.

Fourth, there are concerns about statistical analyses in Bauer et al. (2015). Bauer reported remarkably high intersubject variability with performance ranging from 33% to 97% across 30 subjects (one subject averaged above 80% correct and one below 40%; see their Fig. 3A).

Subjects in the Jones study (13 subjects) showed significantly less intersubject variability as evidenced from small error variances at each temporal position of the target tone (Fig. 3; Jones et al., 2002). The non-significant F values reported by Bauer et al. were in several cases near zero (e.g., 0.05 or 0.22) instead of the expected value of 1 under null; this is often indicative of a violation of the assumptions that ANOVA depends on, including unequal variances across samples, improper randomization, lack of sample independence, or non-homogenous subject populations (Voelkle et al., 2007). Most importantly, the statistical analysis of the “exact replication” condition of Bauer et al. was based on data that was *pooled* with those from a different experiment (see their Figs. 5A and B). In their experiment 5B, the standard tone was deleted from the end of the entrainment sequence (a design different than that used by Jones). A consequence of this “pooled analysis” was that the effect of the “temporal position” variable was diluted.

Fifth, and of equal concern, is that half of the subjects in the “exact replication” experiment of Bauer et al. first completed the “missing standard” experiment (5B) in a counterbalanced within-subjects design. This gave the latter group significant exposure to a *different* (but related) entrainment sequence prior to participating in the “exact replication” experiment. This is important because having been repeatedly exposed to a different sequence may not only have biased subjects into adopting a different detection strategy (a point that Bauer et al. concede), but may have also affected their level of vigilance in a task where attention is a key factor in whether entrainment is observed. As Bauer et al. note, repeating the standard tone just before the comparison “fundamentally alters the task” since the repeated standard becomes a task-relevant tone.

Finally, across their 6 experiments, Bauer et al. (2015) found that 40 out of 140 subjects actually *did* show the same entrainment pattern reported by Jones et al. (2015). This partial replication was observed in spite of differences in subject-selection policy, in experimental design (even in the “exact” replication case), significantly different average performance levels (a 21% averaged difference), and high intersubject variability (33 to 97%). While some of these differences may be minor, others are important methodological differences that need to be recognized, and to claim “exact replication” is inaccurate.

### 2.2. Lin et al. (2021) pitch-discrimination study

In a recent paper, Lin et al. (2021) reported the results of experiments in which rhythmic sequences were used to cue the discrimination of 40-ms FM pulses that either swept up or down in frequency. They measured two performance variables, accuracy (proportion correct) and reaction time. They found no forward entrainment in “pitch discrimination” contrary to findings by Jones et al. (2002). There are, however, important methodological and stimulus design differences between the two studies that raise questions as to the validity of Lin et al.’s conclusions. We begin with a description of their findings, followed by an analysis of their psychophysical methodology and stimulus design properties. We suggest that Lin et al.’s findings are not based on pitch-discrimination ability within a trial, but rather on timbre discrimination, the statistics of which are learned across trials. Furthermore, there are a number of experimental design confounds in how they set up their entraining sequences which reduce the likelihood of observing forward entrainment.

Lin et al. (2021) describe four experimental conditions comprising two cuing and two target conditions (2×2 design). The cue was either a rhythmic sequence of four pure tones (square wave, i.e., 50% duty cycle) or a continuous pure-tone whose duration matched the total duration of the 4-tone rhythmic sequence (including intertone intervals). The continuous pure-tone cue had a fixed duration within a block of trials. Lin et al. inaccurately refer to this steady state and deterministic cue as the “random-cue” condition. Each cue (rhythmic or steady state) was then followed by the target FM pulse. The target was either presented in-phase with the rhythmic sequence (at the onset of one of the first 4 potential cycles after the end of the rhythmic cue) or at a completely random time after the termination of the cue. The rhythmic sequences had rates of 1.4, 1.7, or 2 Hz in experiment 1 (ISIs of 700, 600, and 500 ms respectively), 1.1, 1.7, or 2.5 Hz in experiment 2, and ranging from 0.8 to 10 Hz in their third experiment. In the first two experiments, the sequence rate was fixed within a run. In the third experiment it was randomly selected on each trial from a closed set. Note that the *target* (single FM pulse) is referred to as “rhythmic” even when it was presented in a block of trials that contained *only* the continuous tonal cue (i.e., no rhythm in cue or target). Cue or target types were not mixed within a run, with one exception. On a small subset of trials (20% of “rhythmic cue”-“rhythmic target” condition) they presented targets that were antiphasic to the rhythmic cue. We believe that this particular comparison between the in-phase and antiphasic target conditions (when the cue was rhythmic) is the most appropriate measure of whether psychophysical entrainment could have been observed. The main (and most relevant) finding from Lin et al. (2021) is the absence of a selective advantage of a rhythmic cue in any of the tested conditions (no forward entrainment).^7^ That is, the rhythmic cue did not improve performance for the rhythmic target relative to the random target, nor did the rhythmic cue provide a selective advantage compared to the steady state cue for either target type. The methodological concerns we have with this study are as follows.

First, for rhythmic targets, they *averaged* performance across 4 cycles after termination of the cue. This clearly dilutes any potential forward entrainment effect which is strongest for the first poststimulus cycle, and absent by the 3^rd^ or 4^th^ cycles as others studies have shown (Hickok et al., 2015; Forseth et al., 2020). No separate analysis was conducted on the most important case, the first or second poststimulus cycles in isolation.^8^ It is unclear why Lin et al. averaged potential forward entrainment effects across 4 cycles when Jones et al. (2002) has shown the effect only for the first two cycles, and others have explicitly shown that there is no effect at cycles 3 and 4. In addition, the *entraining* sequence used by Lin et al. comprised only 4 cycles, a significantly smaller number than the 9 cycles used by Jones et al. (2002), Hickok et al. (2015) and Farahbod et al. (2020), as well as a number of others who employed longer sequences in demonstrating forward entrainment (Lawrance et al., 2014; Barnes and Jones, 2000), with some as many as 12 cycles (Lange, 2009; Rimmele et al., 2011). In fact, Wilsch et al. (2020) in a cross-modal study that used entraining auditory stimuli *identical* to those used by Lin et al., suggest that “perhaps, four cycles is not enough to properly drive the system.”

Second, in the Lin et al. study, “random targets” could have occurred at *any* time after the end of the cuing sequence. This means that, on a subset of trials, the “random targets” occurred at or very near the “in-phase” temporal positions. This would have additionally diluted any selective advantage of the in-phase only condition (rhythmic target) since on an unknown proportion of trials, the “random” target would have overlapped with the expected (cued) position of the in-phase targets. For the 40-ms target pulses used by Lin et al., Monte Carlo simulations show that this overlap occurs on a significant number of “random target” trials (Fig. 4). For a 1.7 Hz rhythmic sequence (600 ms ISI), the condition closest to that used by Jones et al. (2002), simulations show that on 6% of the “random target” trials (on average), a segment of the FM target overlapped with the *onset* of the expected tone pulses (had they continued). On some runs, this number could be as high as 15% of trials within a run (top panel). If, however, we assume that the temporal expectancy window generated by the rhythmic cuing sequence is set by the full “on period” (instead of the onset) of the cuing tones, then this overlap occurs on nearly half the trials of a run (bottom panel). These overlap proportions are even larger for the 2 and 2.5-Hz rhythmic sequences. In other words, on a significant number of trials, the “random” FM target could be heard during some portion of the temporal expectancy window generated by the rhythmic sequence, diminishing performance differences between the rhythmic and “random” target conditions. This problem did not exist in the design used by Jones et al. (2002), Hickok et al. (2015), and other studies of forward entrainment because the temporal positions of targets in those studies were restricted to a closed set and performance was reported separately for each of those temporal positions.

Third, and most important, the stimuli used by Lin et al. (2021) are not suitable for testing pitch discrimination as has been claimed. They used single 40-ms FM sweeps whose direction was to be determined by subjects in a single-interval forced-choice task. This stimulus is markedly different than that used for pitch discrimination by Jones et al. (2002) which comprised a *pair* of 150-ms sequentially presented pure tones (standard and comparison). Subjects in Lin et al. were, by our analysis (see below), likely discriminating timbre differences, the perceptual properties of which were learned across trials. Timbre, which is defined by a sound’s subjective quality, “coloration”, and “texture”, requires use of high-level and computationally more complex processes compared to those involved in simple pitch discrimination (Jenkins, 1961; Berger, 1964; Grey, 1977; Iverson and Krumhansl, 1993; Caclin et al., 2005; Mörchen et al., 2006). Two sounds that have the exact same pitch and amplitude, could have significantly different timbres (e.g., same musical note played at the same amplitude on two different violins). Models of timbre encoding (Plomp, 1970; Grey 1977; Caclin et al., 2005) employ multidimensional scaling in dissimilarity space to quantify the timbre of complex sounds. Such high signal dimensionality may not be affected by a brief 4-cycle entraining stimulus in the same manner (or with the same ease) as lower-level processes like pitch encoding.

Why do we suggest that discrimination performance in Lin et al. is based on timbre cues? There are two ways to think about an FM sound. First, in terms of its instantaneous frequency at any given point in time (this is what Lin et al. assume), and second, in terms of its long-term spectrum (Rabiner and Gold, 1975; Saberi, 1997; Saberi and Hafter, 1995; Hsieh et al., 2012). The latter interpretation is important given that the auditory system requires a minimum window of time (the system’s time constant) to process an acoustic waveform. Although, the long-term amplitude spectrum of the up and down sweeps are identical, their phase spectra, which have been shown to affect timbre (Plomp and Steeneken, 1969), are markedly different (Suppl. Fig. 1). If the sweeps used were logarithmic, then both their amplitude and phase spectra are different. These spectral compositions are further altered differentially between up and down sweeps after passing through auditory filters, adding to timbre differences. To gain better insight into the signal processing dynamics and available “pitch” cues that contributed to performance in the Lin et al. study, we processed their stimuli (as well as those of Jones et al., 2002) through an autocorrelation model of pitch extraction (Licklider, 1951; Rabiner, 1977; Shimamura and Kobayashi, 2001; Hsieh and Saberi, 2007; Balaguer-Ballester, 2008). The model comprised a GammaTone filterbank with 50 filters whose center frequencies (CFs) were logarithmically spaced from 300 to ∼3,000 Hz (Holdsworth et al.,1988; Slaney, 1998) followed by an inner hair-cell model (Meddis et al., 1990; Slaney, 1998) and an autocorrelation function:

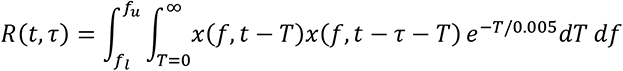

where *x* is the time waveform at the output of each frequency channel, *f_l_* and *f_u_* are the lower and upper filter CFs over which the autocorrelation patterns are integrated, *T* is time into the past relative to current time, the exponential decay has a time constant of 5 ms, and τ is the autocorrelation lag.

The left panels of Fig. 5 show the output of this model in response to sample stimuli used by Jones et al. (2002). This included a standard tone followed, after an ISI, by a comparison tone that is one semitone higher in frequency than the standard tone.^9^ Panel A shows the time-by-frequency output of the filterbank stage. The red and light blue horizontal lines are centered at the frequencies of the standard and comparison tones respectively. Panel B shows the model’s normalized autocorrelation function, integrated across frequency channels (Eq. 1). Peaks at non-zero lags (near 2 ms) are the model’s pitch estimates (i.e., inverse of the lag value). The model clearly predicts significant pitch differences between the two stimuli. Panel C (bottom) shows the model’s time-frequency response to the standard stimulus (magnification of the trace from panel A). The right panels of Fig. 5 (panels D, E, and F) show the model’s output in response to a 40-ms FM pulse. In Lin et al.’s (2021) single-interval task, subjects had to detect whether the FM sweep was up or down (discriminate pitch differences within the single 40-ms sound). Panel E (middle right) shows that the autocorrelation function produces no pitch estimate for these pulses (no peak at a non-zero lag). To more carefully analyze the dynamic nature of pitch cues in their stimuli, we used a short-term (running) autocorrelation function with a 5ms exponential decay time constant to dynamically update pitch estimates throughout the duration of the stimulus, and again found no dynamic pitch estimate (secondary peak) as shown in [Video 1](http://www.socsci.uci.edu/~saberi/periodicity/video1.avi).^10^

In addition, the response of auditory filters to brief pulses is asymmetric, partly because cochlear filters have a sharper slope above the filter’s CF, and partly due to the nonlinear dynamics of cochlear hair cells. This asymmetry may be observed in Fig. 6 which shows the temporal envelope and fine structure response of the filterbank and hair-cell model described earlier to the type of 40-ms FM sweeps used by Lin et al. (2021). Note the asymmetries across the two sweep types in their declining envelope amplitudes as a function of time, as well as the changing AC amplitude (fine structure variance) which is the main determinant of spectral content. For example, in the top panel, the higher frequency segment of the FM will generate greater spectral energy (near 35ms) than the lower-frequency part of the FM (near 10ms) in spite of the larger envelope amplitude at the low-frequency end. Thus, these stimuli can generate perceptually dynamic properties, which include dynamic timbre (Iverson and Krumhansl, 1993) and loudness cues. There is also evidence that a greater number of cortical neurons code for up (compared to down) FM sweeps in some regions of the auditory cortex (mediolateral belt), and in the opposite direction in other regions (anterolateral belt) (Tian and Rauschecker, 2004). This neural population differences may potentially further contribute to perceived timbre difference in unknown ways between the two types of pulses, and may also be the source of the asymmetry observed in psychophysical performance between identification of up and down FM sweeps (Luo et al., 2007).^11^ To be clear, although timbre is by our estimation the predominant cue used to discriminate the up from down FM pulses used by Lin et al., we do not say that there is *no* pitch cue, but rather that any such cue would be very weak and potentially confounded with dynamic timbre cues, and even by changing loudness cues from asymmetric envelopes at the output of cochlear filters. One can, in fact, force a weak pitch cue at the model’s output by introducing a nonlinearity (square or cubic) that enhances the ridges of the filtered FM pulse, though even in this case, such putative cues are significantly weaker than those in the stimuli used by Jones et al. (2002).

Finally, one can also demonstrate that the cues used by subjects in the Lin et al. study are not pitch based either by shifting the entire FM sweep to regions of the acoustic spectrum above 5 kHz ([audio demo 1](http://www.socsci.uci.edu/~saberi/periodicity/audio1.wav): up followed by down sweep, 5.5 to 7 kHz) where pitch cues as defined by ANSI (1973) are nonexistent (or extremely weak) due to the upper bound on neural phase locking (Ward, 1954; Moore, 1973; Palmer and Russell, 1986; Semal and Demany, 1990; Yost, 2009) or by reducing the duration of the FM sweep to 10 ms (5ms rise-decay; [audio demo 2](http://www.socsci.uci.edu/~saberi/periodicity/audio2.wav): up followed by down sweep, 0.5 to 2 kHz), too brief to allow salient pitch cue discrimination *within* the pulse. In both cases (shifting above 5 kHz or reducing duration to 10 ms) the same qualitative perceptual differences (largely dynamic timbre and loudness differences) is heard between up/down sweeps as those heard when listening to the sweeps of the type used by Lin et al., i.e., 40ms FMs at lower frequency regions ([audio demo 3](http://www.socsci.uci.edu/~saberi/periodicity/audio3.wav): 0.5 to 2 kHz, or [audio demo 4](http://www.socsci.uci.edu/~saberi/periodicity/audio4.wav): 0.6 to 0.9 kHz).

To summarize, there are several methodological concerns that raise questions as to the validity of findings reported by Lin et al. (2021). These include averaging performance across 4 poststimulus cycles, use of short entraining sequences (4 cycles), inaccurate definition of a “random” entrainment (using deterministic stimuli), overlap between “random” target times and the temporal expectancy window set by rhythmic sequences on a significant proportion of trials, and confounds associated with timbre as a signal, the detection of which involves computationally complex multidimensional cues that are not likely ideal for detecting forward entrainment.^12^

### 2.3. Sun et al. (2021) signal-detection study

Sun et al. (2021) reported the results of two experiments in which they used stimuli identical to those of Hickok et al. (2015) to investigate forward entrainment in signal detection. Their main finding was that while there was an inverse-U shape pattern in poststimulus performance, no bicyclic (M-shaped) pattern was observed at the overall group level. Approximately 35% of their subjects did, however, show forward entrainment but with no cross-subject consistency in entrainment phase (i.e., power at 3 Hz, but no antiphasic effect). In this section, we describe some of the methodological differences as well as data-analysis dissimilarities between the two studies that may help in understanding the discrepant findings.

As described in section 1.1, Hickok et al. (2015) introduced trial-by-trial level uncertainty by randomly selecting signal level from one of 5 intensities. In their first experiment, Sun et al. used only 2 signal levels. This is important because Farahbod et al. (2020) has shown that level uncertainty appears to be critical for observing forward entrainment, at least in the signal-detection design used by Hickok et al. (2015). Because of this difference in design (among other methodological differences) Sun et al. referred to their first experiment as a “conceptual replication”. Based on Farahbod et al.’s (2020) finding on uncertainty, Sun et al. then ran a second experiment which they referred to as an “exact” replication. However, even in this case, there were nearly a half dozen methodological, procedural, and experimental design differences between their exact replication and the original Hickok et al. (2015) study. Most of these procedural differences are outlined in footnote 13.

The most critical difference between the two studies, however, is in their data-analysis approach. In the main part of their manuscript, Sun et al. used “hit rates” as a dependent measure, restricting their analysis only to “signal trials” (trials that contained a tone to be detected) and eliminating from analysis all no-signal trials. By excluding no-signal trials, Sun et al. effectively used half as many trials as Hickok et al. who evaluated performance using “proportion correct” as the dependent measure. This latter measure includes both signal and no-signal trials. Sun et al.’s reasoning for excluding no-signal trials was to reduce “random response noise”. What they refer to as “random noise”, however, is in reality a measure of false-alarm rates which is just as informative as hit rates in evaluating performance. Exclusive reliance on hit rates in a *single-interval yes-no task* is a critical flaw that produces inaccurate results unless false alarms are also taken into account. In an extreme case, a subject could simply respond “yes” on every trial and get 100% hit rate without attending to the stimulus. Use of “hit rate” as a dependent measure by itself is therefore both misleading and biased. This is not a significant problem for “proportion correct” as a dependent measure (used by Hickok et al., 2015) because indiscriminately responding “yes” on every trial will lead to chance performance (0.5 correct response probability).

Figure 7 shows hit-rate data from Sun et al. (2021) plotted individually for two subjects from their experiment 2. Also included are their false-alarm rates (not shown in Sun et al.’s original paper).^14^ A cursory examination illustrates our concern. Subject 205 has an average hit rate of 0.87, and subject 221 has an average hit rate of 0.66. If we use an unbiased measure of performance, such as the detection index *dꞌ*, it becomes clear that subject 221 is actually significantly outperforming subject 205 in detecting the tonal signal (*dꞌ* = 2.33 vs 1.92). The problem of course is that subject 205 has a liberal criterion that results in a disproportionately large number of “yes” responses (hence the high false-alarm rates for this subject) whereas subject 221 is much more strict (or careful) in reporting “yes”, as evident from this subject’s very low false-alarm rates. This problem is also observed in the data of several other subjects from Sun et al. shown in Supplemental Fig. 3. For example, subjects 202, 217, and 219 all have significantly lower hit rates, but higher performance levels than subject 205 when evaluating performance using an unbiased measure (*dꞌ*).

**Figure 7.**
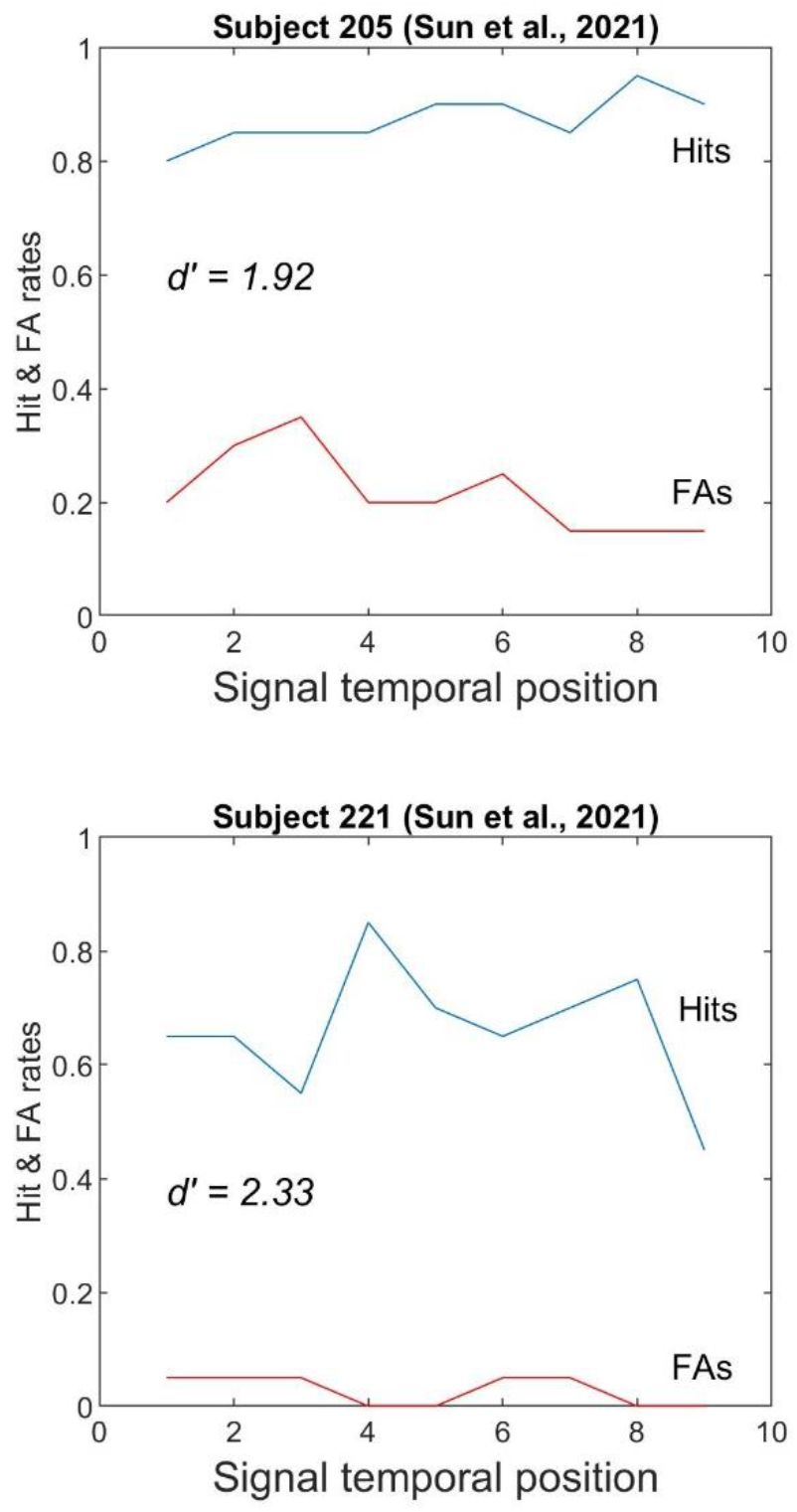
Hit and False-Alarm rates for two subjects from Sun et al. (2021). Subject 205 (top) has a higher averaged hit rate but lower *dꞌ* than subject 221 (bottom) when false-alarms are taken into account in calculating an unbiased measure of performance.

To assess the effects of using a biased versus unbiased measure on their results, we reanalyzed the raw data of Sun et al. (2021) using a detection-theoretic approach. For each of 23 subjects we measured *dꞌ* and percent correct at each temporal position.^15^ We used the same signal level as that used for analysis by Sun et al. (2021) and Hickok et al. (2015), i.e., SNR 2 (3 dB). The averaged curve for the 23 subjects is shown in panel A of Fig. 8 as detection index *dꞌ* and as proportion correct. A bicyclic (M-shaped) pattern of performance is observed when an unbiased measure *dꞌ* is used instead of “hit rates”, with a dip at temporal position 5 which is antiphasic to the expected modulation peak. A similar, though less prominent, M-shaped pattern is observed for percent correct.

A repeated-measures ANOVA on the individual-subject *dꞌ*s of Panel A shows a highly significant effect temporal position (F(8,176)= 2.85, p=0.005; η^2^ = 0.12). Post-hoc paired-samples t-tests showed significant differences between temporal positions 5 (dip) and 8 (right peak) t(22)=2.30, p=0.031, and while there was no significant difference between temporal positions 5 (dip) and 4 (left peak) t(22)=1.68, p=0.107, a one-tailed test supporting the hypothesis that the dip at position 5 would be antiphasic to the expected modulation envelope (i.e., generate poorer performance) is near significant, p=0.0503 (see also our discussion of panel B below). Post-hoc t-tests on *dꞌ* data of panel A also showed a significant difference between temporal positions 8 and 9, t(22)=3.04, p=0.006 and no significant difference between the two peaks at temporal positions 4 and 8, t(22)=0.21, p=0.83. A repeated-measures ANOVA on the proportion correct data (Panel A) showed a highly significant effect of temporal position (F(8,176)=5.36, p<0.001; η^2^ = 0.20). Post-hoc paired-samples t-tests showed significant differences between temporal positions 5 (dip) and 8 (right peak) t(22)=2.45, p=0.023, no significant difference between positions 4 (left peak) and 5 (dip), t(22)=0.838, p=0.41, no difference between the two peaks at temporal positions 4 and 8 (t(22)=1.58, p=0.13), and a significant difference between temporal positions 8 and 9 (t(22) = 5.62, p<0.001).

Panel B shows proportion correct and *dꞌ* data from 12 of the 23 subjects who showed strong entrainment effects in Sun et al.’s exact replication experiment (more than half of their subjects in that experiment).^16^ The data of this subpopulation is particularly interesting not just because of the clear bicyclic (M-shaped) pattern in both measures (*dꞌ* and proportion-correct), but also, more importantly, because statistical analysis shows that the dip at temporal position 5 is significantly different than the peaks at positions 4 and 8, and that there is no statistical difference between the two peaks. A repeated measures ANOVA on the proportion correct data of these subjects showed a significant effect of temporal position (F(8,88)=4.54, p<0.001; η^2^ = 0.29). As noted above, post hoc t-tests showed a significant difference between temporal position 5 (dip) and position 4 (left peak) t(11)= 2.82, p=0.017, and between positions 5 and 8 (right peak) 5v8 t(11)=4.07, p=0.002, but no difference between the two peaks at 4 and 8 (t(11) = 0.81, p=0.437). This suggests that the antiphasic dip is likely a real effect. There were also no statistically significant differences between temporal positions 1, 5, and 9, i.e., the 3 dips at the start, middle, and end points (1 vs 5: t(11)= 0.293. p=.775; 5 vs 9: t(11) = 0.328, p=0.749; 1 vs 9: t(11) = 0.558 p=0.588). Similar results were obtained when analyzing *dꞌ* data (panel B). A repeated measures ANOVA on the *dꞌ* data showed a significant effect of temporal position (F(8,88)=3.54, p=0.001; η^2^ = 0.24). Post hoc t-tests showed a significant difference between temporal position 5 (dip) and position 4 (left peak) t(11)= 4.38, p=0.001, and between positions 5 and 7 (right peak) t(11)=3.43, p=0.006 (also between positions 5 and 8, t(11)=4.09, p=0.002). However, there was no difference between the two peaks at positions 4 and 7 (t(11) = 0.66, p=0.52) nor between positions 4 and 8 (t(11)=0.99, p=0.34). This suggests that the antiphasic dip is likely a real effect. There were also no significant differences between temporal positions 1, 5, and 9 (1 vs 5: t(11)= 0.76, p=.46; 5 vs 9: t(11) = 0.77, p=0.46; 1 vs 9: t(11) = 0.02 p=0.98). For comparison, the data of Hickok et al. (2015) are plotted in Panel C both as *dꞌ* and proportion correct. Note that both in the Sun et al. and Hickok et al. data, the second peak is larger than the first (higher performance level). This may potentially be related to stronger masking effects near the earlier parts of steady state noise (see section 4.3 below; Samoilova, 1959; Zwicker, 1965; Elliott, 1971).^17^

We then asked the following question: If there are slight subject-specific phase shifts (or jitter) in entrainment as previously suggested (Henry and Obleser, 2012) or as seen in rate-specific phase drifts suggested by Farahbod et al. (2020), could minor phase adjustments generate an even stronger bicyclic (M-shaped) pattern of psychophysical performance across all 23 subjects of Sun et al.’s study? To test this, we shifted the individual subject curves that produced the average *proportion-correct* curves of Fig. 8A by *at most* a single time point (i.e., at most a quarter of a cycle) in the direction that would maximize alignment of the individual curves at their antiphasic minima. The result, shown in Suppl. Fig. 5 confirms that even a slight phase adjustment across subjects results in a strong bicyclic pattern of proportion-correct performance averaged across 23 subjects.^18^

**Figure 8.**
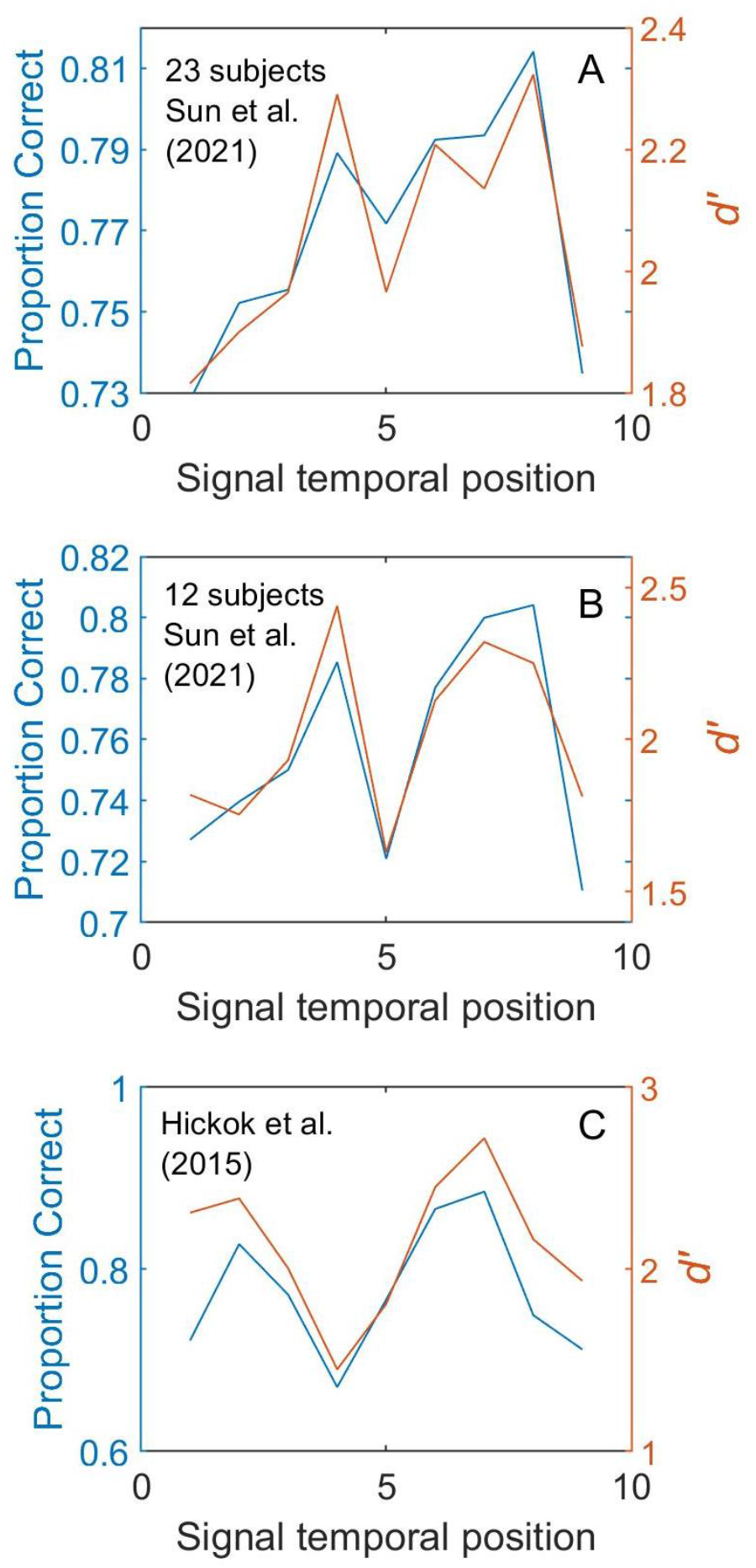
Panel A: Data from Sun et al. (2021) plotted as proportion correct and as an unbiased index of detectability (*dꞌ*) instead of hit rates averaged across 23 subjects. Note the antiphasic dip at temporal position 5. B: *dꞌ* and proportion correct for 12 of 23 subjects from Sun et al. (2021) who showed strong forward entrainment. C: Data from Hickok et al. (2015).

We conducted additional analyses on the raw data of Sun et al. that made use of a larger portion of their dataset. Specifically, we measured performance at peak-phase compared to trough (envelope minima) target positions based not just on a single SNR, but on 3-point psychometric functions that made *simultaneous* use of 3 SNRs, tripling the number of trials from which thresholds are estimated. First, for each subject we measured *dꞌ* psychometric functions for three peak-phase and 2 trough-phase temporal positions (see sinusoidal legend in Fig. 9). We then averaged these functions across the 23 subjects as well as within a “phase category” (peak or trough). This resulted in two 3-point psychometric functions, one for each phase category. We defined SNR threshold to be the point at which regression fits to each psychometric function crossed a specific performance

**Figure 9.**
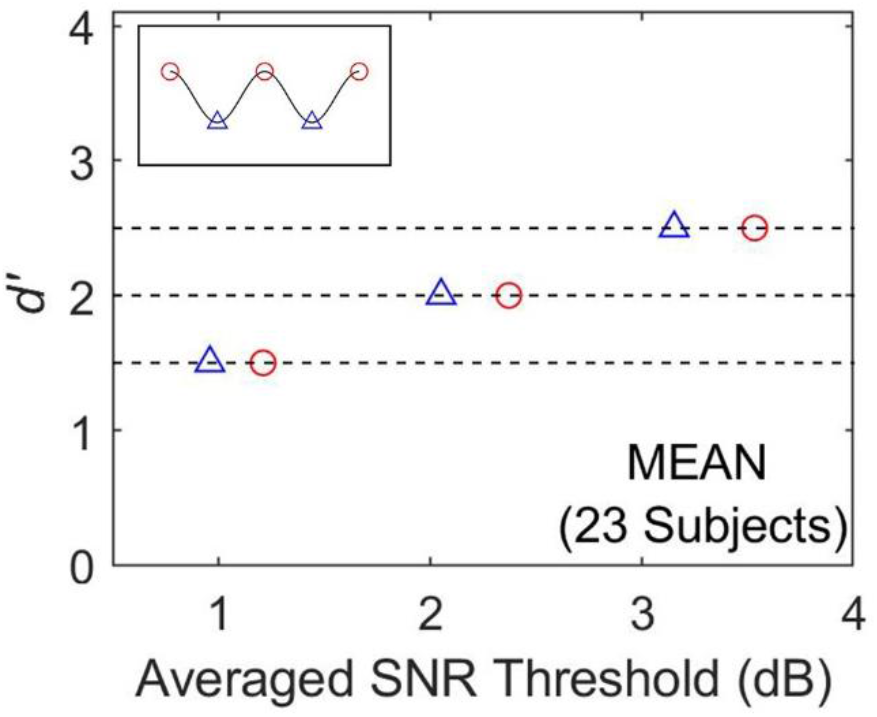
SNR thresholds at 3 *dꞌ* levels calculated from psychometric functions averaged across 23 subjects for each of two “phase categories” (peak-phase = red; trough = blue). Note that the peak-phase category requires a higher SNR threshold to reach a performance level (*dꞌ*) equivalent to that for the trough category (envelope minima). Legend is shown in upper left.

level (e.g., *dꞌ* = 1, 2 or 3). Results shown in Fig. 9 demonstrate that when performance is measured in *dꞌ* units using a larger dataset comprising 3 SNRs, temporal positions at the expected dips of the modulation envelope consistently generate better performance than those at the peaks for any of the 3 *dꞌ* levels at which performance was estimated (i.e., a higher SNR value is needed to reach an equivalent *dꞌ* performance level).

Sun et al. also performed a supplemental analysis of hit-rate curves in which they excluded subjects with high false-alarm rates. The goal was to eliminate “biased subjects”. This, however, misinterprets the meaning of bias in signal-detection terms. Bias here does not mean high false alarms. A subject could be biased *against* responding “yes” and generate very low false-alarm rates. Evaluating performance based on hit rates is biased because it only examines one aspect of performance (hits) and excludes the complementary and inherently linked false-alarm rates. One without the other is a meaningless measure. In a single-interval yes-no task, the only truly unbiased case occurs when the subject places their decision criterion at precisely the intersection of the *internal* “noise alone” and “signal plus noise” probability distributions (Green and Swets, 1966; Macmillan and Creelman, 2005). This is of course unknown to the experimenter and inconsistent across subjects, so the only way to ensure unbiased measurements is to combine hit and false alarm rates in estimating the detection index *dꞌ*. Sun et al. further justify their use of hit rates as their main approach by noting “*One might argue that when participants’ overall hit rate is too high (clearly above threshold) or too low (clearly below threshold), it reduces the likelihood of observing fluctuations across different temporal positions … If there was any truth in this conjecture, one would expect to observe a clearer presence of the entrainment effect across participants whose overall hit rate is confined within a narrower range, …we computed a range-of-interest, … then selected… participants whose overall hit rate… was enclosed in this range… Results from this more restricted analysis showed that the average modulation strength of selected data is not significantly above zero*”. The flaw in this logic can easily be demonstrated by comparing the unbiased performance of subjects who have similar hit rates.

Subjects 222 and 223 have identical hit rates of 0.55556 but very different *dꞌ*s (2.68 vs. 1.97) with subject 222 significantly outperforming subject 223 in detecting the signal. Therefore, it is not methodologically valid to attempt to equate subject performance based on hit rates, or to define a range-of-interest hit rates, in order to eliminate “biased” subjects or outliers with markedly different performance levels.

A second major flaw in Sun et al.’s analysis, other than reliance primarily on hit-rate measures, has to do with the way in which they define the magnitude of entrainment, what they refer to as “modulation strength”, on which they base nearly all of their analysis and from which they draw nearly all of their conclusions. This measure, which is the normalized Fourier transform of the performance curves (mostly hit-rate curves) is not only erroneously calculated but is also unreliable in estimating the strength of entrainment. In defining “modulation strength”, Sun et al. note that they “*performed Fourier transforms of the curves of target detectability (hit rates) for each participant. Given that each curve only contained 9 temporal locations covering a duration of 667 ms, a Fourier transform of these data only provided accurate power estimation at 4 frequencies, corresponding to 1.33 Hz, 2.67 Hz, 4 Hz, and 5.33 Hz. Among these four frequencies, we selected 2.67 Hz – which is closest to the modulation rate (3 Hz) in the experiment – to be the frequency of interest and measured its power for each participant*.” In over 30 places in their paper they note the use of 2.67 Hz as their base analysis. Their Fourier transform calculations, however, are in error. In Fourier analysis, the frequency component spacing is derived from the inverse of the duration of the function whose spectrum is to be calculated. There are 9 points at which performance is measured, temporally spaced at a quarter of a cycle of the 3 Hz sinusoid with successive points separated by 83.33 ms. The “duration” of the performance curve is equal to exactly two cycles at 3 Hz, or 666.666… ms (the devil is in the details). The frequency component spacing is therefore 1/0.666… or exactly 1.5 Hz. Thus the FFT contains energy precisely at 1.5, 3, and 4.5 Hz (not 1.33, 2.67, 4, and 5.33 Hz). In fact, the discrete Fourier transform of this function has no energy at 2.67 Hz to be measured. Because the sampling *rate* of the performance curves is 12 Hz (4 points per cycle at 3 full cycles per second), the frequencies at which spectral energy can be measured have to be below the Nyquist limit to avoid aliasing (i.e., half the sampling rate) and therefore measurements are restricted to component frequencies below 6 Hz (1.5, 3, and 4.5 Hz). We trace the error made by Sun et al. to miscalculating the performance curve “duration”. They appear to have assumed that 9 points results in 9 quarter-cycle segments (instead of n-1 segments). Each segment is 83.33 ms in duration, hence 9×83.33=750 ms, the inverse of which is 1/0.75 or 1.33 Hz component spacing which results in the erroneous estimates of 1.33, 2.67, 4, and 5.33 Hz. The fact that the idealized performance curve was originally *defined* by a 3-Hz sinusoid with points precisely at quarter-cycle intervals for exactly two cycles makes it intuitively obvious that the transform must have contained energy at that frequency (3 Hz).

Separate from this calculation error, there is a more fundamental concern with using the Fourier transform to estimate “modulation strength”, one that Farahbod et al. (2020) had previously cautioned about. When using very brief signals, such as the 2-cycle waveform associated with behavioral performance curves, the duration of the waveform itself (i.e., the analysis window), regardless of the shape of the waveform within that window, will enhance energy at some regions of the spectrum and diminished energy at others. This can confound “modulation strength” measurements. Theoretically, a 2-cycle sinusoid is generated by multiplying an infinitely long sin-wave with a rectangular (boxcar) window in the time domain. The spectrum of such a waveform is the convolution of the spectra of the infinitely long sinusoid (a vertical line with zero bandwidth) with that of the rectangular window (the well-known sinc function). The sinc function has zero-crossings in the magnitude spectrum at the inverse of the window’s duration. The sinusoid’s frequency and the width of the temporal window constraining that sinusoid jointly determine the position of these zero-crossings in the spectrum. The effects of the sinc function on the spectrum can constructively or destructively alter the amplitude of frequency components used to measure “modulation strength” (see Suppl. Fig. 6). Furthermore, for such brief waveforms, the starting phase of the sinusoid itself affects the spectral profile of that waveform. These interactions can generate false positives or misses in identifying whether energy at a given frequency component results from entrainment or from signal-processing artifacts. We therefore caution against use of the discrete Fourier transform in determining the strength of entrainment when analyzing very brief waveforms since their spectra are too severely altered by the spectral profile of the rectangular window that defines their duration.

There are a number of other observations about the findings of Sun et al. that are worth considering here. Sun et al. note that the methodological differences between their two experiments (“conceptual” and “exact” replication) are relatively minor and inconsequential. If this is the case, then one has to explain several significant differences in the results of these experiments. First, as they acknowledge, the two experiments produced peak performance at different temporal positions (time positions 5 and 6 in experiment 1, and positions 6 to 8 in experiment 2). Second, the average starting phase of forward entrainment, for the ∼35% of subjects who showed entrainment, were significantly different across the two experiments. In their experiment 1, subjects had averaged starting phases of 102 degrees (0.6π radians) whereas those in their experiment 2 had an average starting phase of 220 degrees (1.22π radians). Interestingly, the latter is less than a quarter of a cycle (0.14 cycles) away from the antiphasic position (270 degrees, 1.5π) and approximately a tenth of a cycle away from that reported by Hickok et al. (262 degrees, 1.46π).^19^

As mentioned above, a notable difference between the findings of Hickok et al. (2015) and Sun et al. (2021) is the significantly larger intersubject variability in the latter study. Sun et al. contend that “larger cross-participant variability is generally expected with increased sample size”. Increasing subject sample size improves the precision with which the population variance in performance is estimated but has no systematic effect on the comparative size of the variances of two samples if they are unbiased estimators of the population variance. These sample variances, however, will be different if the nature of the underlying populations from which they are sampled are different. One potential population difference is subject experience. Subjects in Hickok et al. (2015) were highly experienced having previously participated in a number of auditory psychophysical tasks (graduate students and postdocs). Sun et al. do not report whether their participants had extensive experience or were experimentally naïve (they used 47 subjects who completed one or two experimental sessions). While we do not know the level of prior psychophysical experience of their subject population, this may be a factor worth considering, especially given new data from our lab that demonstrates more variable patterns of performance for inexperienced and perhaps less motivated subjects (see section 3.4 below).

An interesting finding by Sun et el. (2021) was that a proportion of their subjects (35%) did in fact show forward entrainment even using their biased measure, but that there was no consistent phase alignment and no antiphasic pattern to performance. Bauer et al. (2015) also reported that 40 of their 140 subjects (28%) showed forward entrainment in a pitch-discrimination task (see section 2.1 above). As noted earlier, if Sun et al.’s data are reanalyzed using unbiased estimators of performance such as *dꞌ*, a bicyclic pattern with an antiphasic dip is in fact observed at the full population level (Fig. 8A). We would, nonetheless, like to make that point that even by their own standard of using hit rates, the finding that a segment of the tested population does show the effect and a segment does not, is actually confirmation of the existence of the effect in a subcategory of subjects, and to conclude that an absence of statistical significance at the full sample population level is evidence against existence of such an effect is misleading. There are several auditory phenomena that are consistently observed in a segment of normal-hearing populations and not in others for reasons that may range from experience to genetics (Chubb et al. 2013; Assaneo et al., 2019; Mednicoff et al., 2018; Ho and Chubb, 2020).

Finally, we would also like to briefly comment on Sun et al.’s speculation as to why there were contrasting findings between their study and Hickok at al. (2015). They attribute this to a number of potential factors, including the details of the experimental environment, an accidental underrepresentation of listeners who do not exhibit forward entrainment (note that 35% of their subjects did show entrainment), or subject-dependent entrainment phases which would flatten out bicyclic effects at the group level. They dismiss this latter explanation, arguing that the previously reported subject-specific phase dependency (Henry and Obleser, 2012) was observed only for FM sounds, not for AMs. However, there is significant evidence for FM-to-AM conversion in the auditory periphery as the instantaneous frequency of an FM signal sweeps through the passband of auditory filters (Henning, 1980; Saberi and Hafter, 1995; Saberi, 1998). This conversion pattern is complex and dependent on both the integration time constant of the system and the relative position of an auditory filter to that of the FM carrier. The induced-AM signal resulting from a sinusoidal FM will have a rate that is twice that of the FM at the output of a filter centered on the FM carrier, and match that of the FM rate if the sweep only passes through the lower (or upper) skirt of an off-center filter (but not both). Off-frequency listening away from the FM’s carrier frequency will therefore result in use of AM cues equivalent to the entraining FM rate. Furthermore, several subjects in the study by Henry and Obleser (2012) show a clear antiphasic pattern of behavioral performance relative to the entraining stimulus phase (their Fig. 3 and Fig. S2) while others show an in-phase pattern (or at a different phase).

This may be partially related to implicit subject-specific strategies in off-frequency listening at the output of a filter that is either above or below the FM’s carrier frequency. Listening at the output of a filter above the FM carrier when the instantaneous frequency of the FM is at its lowest value (antiphasic) will maximize SNR since the induced-AM at the output of that filter will be at its minimum amplitude. Off-frequency listening at the output of a filter below the FM carrier will generate the opposite pattern. Thus, there are significant shared mechanisms in how the auditory system processes FM and AM sounds and the subject-specific phase dependency in entrainment reported for FM sounds could have relevance to AM signal processing.

Sun et al. also further speculate on other potential factors that may result in an absence of forward entrainment. They suggest, citing Bauer et al. (2015), that when there are conflicting spectrotemporal cues in the entrainment process (as when the entraining stimulus and the signal are not of the same stimulus class) one may fail to observe forward entrainment. Sun et al. then note that “there is no direct correspondence between the spectral content of the entraining signal (broadband noise) and target stimulus (1 kHz tone)” in their design or that of Hickok et al. (2015). They conclude that it is therefore unclear why an AM *noise* would trigger entrainment that would facilitate the detection of a *tone* pulse (i.e., different classes of sounds). Setting aside the fact that the noise spectrum contains energy at the frequency of the tone that it masks, what Sun et al. fail to recognize is that the entraining AM noise, as suggested by Simon and Wallace (2017), entrains *against* the noise that limits signal detection. From this standpoint, the similarity in spectral characteristics of the entrained and entraining noise can facilitate better isolation of the target to be detected during expected dips (see also the discussion of CMR in section 4.3 below in how one narrow band of noise can entrain a spectrally remote band of noise to improve the detection of a tonal signal). Importantly, however, our reanalysis of their data using unbiased dependent measures argues that appeal to such explanations is generally unnecessary as their study clearly demonstrates forward entrainment when appropriate dependent measures are used.

## 3. Constraints on Forward Entrainment

In our own research, we have observed forward entrainment under some circumstances (as described in section 1.1) and failed to observe the effect under other experimental conditions. These observations are relevant in understanding the variability reported in the broader literature, as reviewed above. In this section, we describe those stimulus and experimental design conditions that generally did not result in poststimulus entrainment to an acoustic modulator.

### 3.1. Effects of signal uncertainty

Selective attention has been shown to improve psychophysical performance under conditions of uncertainly in a number of standard auditory tasks, for example in tasks involving the detection of tones of uncertain frequency (Dai et al., 1991; Hafter and Saberi, 2001; Hafter et al., 2008; Schlauch and Hafter, 1991; Wright and Fitzgerald, 2017) uncertain duration (Dai and Wright, 1995), or uncertain time of occurrence (Bourbon et al., 1966). As uncertainty increases and predictability decreases, the system’s limited attentional resources are allocated to monitoring specific time points set by the rhythmic entraining sequence, resulting in a brief attentional cadence after termination of the entraining stimulus. Removal of signal uncertainty mitigates the need for selective attention and diminishes potential poststimulus modulatory effects in signal detection. This is what we observed when we removed level uncertainty in our forward entrainment paradigm (Farahbod et al., 2020). When signal levels and temporal positions of the signal were mixed within a block of trial, we observed forward entrainment in an antiphasic pattern, yet when we removed level uncertainty in the same listeners, no modulatory effect (no forward entrainment) was observed. The fact that the signals used by Hickok et al. (2015), Farahbod et al. (2020), Forseth et al. (2020), and others were near detection threshold (barely audible) may have additionally contributed to signal uncertainty. Other studies of forward entrainment have also typically used a design in which some aspect of the stimulus was uncertain, usually by mixing signals of various delays or frequencies across trials within the same run (Jones et al., 2002; Barnes and Jones, 2000; Lange, 2009; Ellis and Jones, 2010; Lawrance et al., 2014; Forseth et al., 2020). At least in one case of failure to observe forward entrainment, a block design with little uncertainty in rhythmic target conditions was used (Lin et al., 2021). This, however, is not always the case (Bauer et al., 2015) and additional studies on the role of uncertainty on forward entrainment would be warranted.

### 3.2. Effects of temporal envelope complexity

Nearly all studies of forward entrainment have employed simple rhythmic modulation patterns such as sinusoidal, squarewave, rectangular, or triangular sequences with a simple rhythm. To determine if the effects of masker modulation on signal detection can persist for more complex modulation patterns, we repeated the Hickok et al. (2015) study using noise maskers that were simultaneously modulated at more than one rate. Two such complex patterns were examined: 1) combined modulation rates of 2 and 3 Hz (i.e., the two rates at which the strongest forward entrainment was reported by Farahbod et al., 2020), and 2) combined modulation rates of 3 and 5 Hz. Five subjects participated in the 2+3 condition and four subjects in the 3+5 condition. Each subject completed a minimum of 30 runs per condition, with 100 trials per run. All other procedures, methods, and stimuli were the same as those described by Hickok et al. (2015) and Farahbod et al. (2020). Figure 10 shows results of this experiment. These are new data not previously reported elsewhere. The blue solid lines show average performance across subjects and the red dotted curves show the expected pattern of modulation had it continued through the stationary noise. The mean and variance of the red curves were selected to match those of the data in each panel. Although there appears to be a curvature to the shape of the data in these panels, they do not appear to consistently follow the more complex shape of the modulating envelopes. This suggests that either the modulation pattern is too complex to affect signal detection or the combined frequencies result in an average modulation rate that is too high to yield entrainment (Farahbod et al., 2020). Interestingly, Henry et al. (2014) have demonstrated simultaneous (not forward) entrainment using mixed-modulated waveforms (simultaneous FM and AM) presented at different rates (∼3 for FM and ∼5 Hz for AM) in a gap-detection experiment. The interaction patterns between the FM and AM modulators, which were presented within the same low-frequency region of the spectrum (800 to 1200 Hz), are complex because of FM-to-AM transduction in the auditory periphery as described in section 2.3 of the current paper.^20^

**Figure 10.**
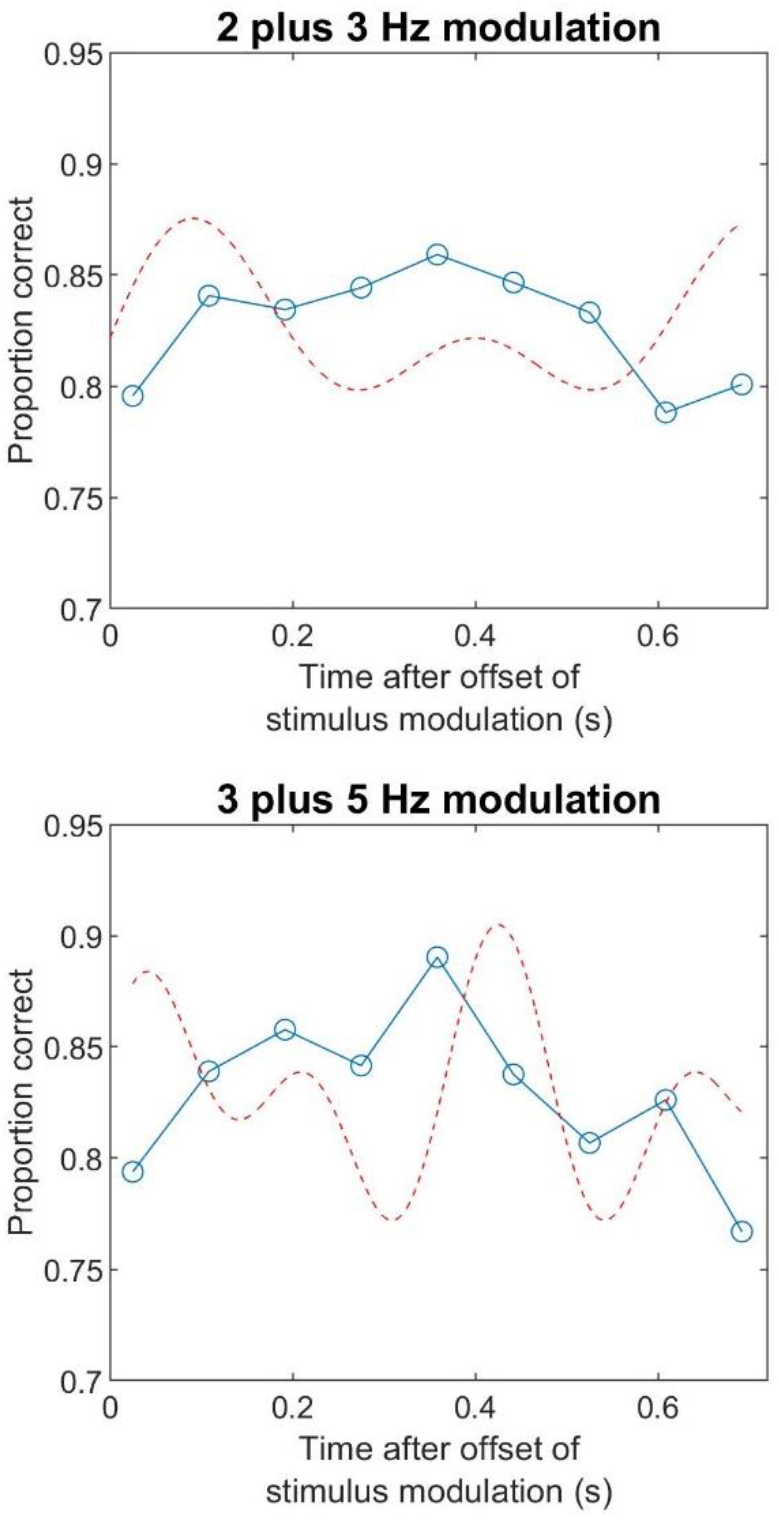
Absence of forward entrainment for complex modulation patterns. Red dotted curves show the expected modulation envelopes.

### 3.3. Effects of SNR

Hickok et al. (2015) reported forward entrainment at one SNR that generated the largest modulatory effect. A reasonable concern is that the reported effect is based on a low number of trials since performance was measured using a subset of trials (i.e., one of 5 intermixed SNRs). Here we show results for all other SNRs collected simultaneously with those reported by Hickok et al. (2015). The 5 SNRs spanned a 12 dB range, all intermixed within the same block of trials. The top panel of Fig. 11 shows performance as a function of signal temporal position averaged across subjects. Time zero represents the end of the entraining stimulus (end of noise modulation). The parameter is SNR, with the 3.5 dB condition representing the data reported by Hickok et al. (2015), and the other 4 SNRs showing previously unpublished data from the same experiment. The 0-dB SNR represents baseline (lowest signal level tested). A few points are worth emphasizing here. First, on majority of trials, the signal was clearly audible as is evident from the three curves at performance levels above 90%. This, we believe, adds to the uncertainty of detecting the less audible signals at 0 and 3.5 dB, and perhaps, as discussed above, enhance the role of selective attention. Second, no modulatory effect is observed for two of the three highest SNRs, and a mild bicyclic effect for SNR of 9.5 dB. This may just be chance variation or possibly related to a weak forward entrainment effect. While the absence or weak modulation effects at high SNRs is partially associated with ceiling effects, it may also be related to the fact that the higher SNRs reduce uncertainty and need for selective attention. At the lowest SNR tested (0 dB), while we do not observe the same bicyclic pattern as that reported by Hickok et al. (2015), there does seem to be a dip in performance in line with that seen for the 3.5 dB SNR data. The difference between the dip at 0 dB SNR and the peak on the same curve approaches statistical significance (t(4)=2.67, *p* = 0.056). A one-way ANOVA showed no significant effect of signal delay (no forward entrainment) at 0 dB SNR (F(8, 36)=1.88, *p* = 0.094) even though nearly all points at this SNR are above chance and not constrained by floor performance (0.5 probability).

**Figure 11.**
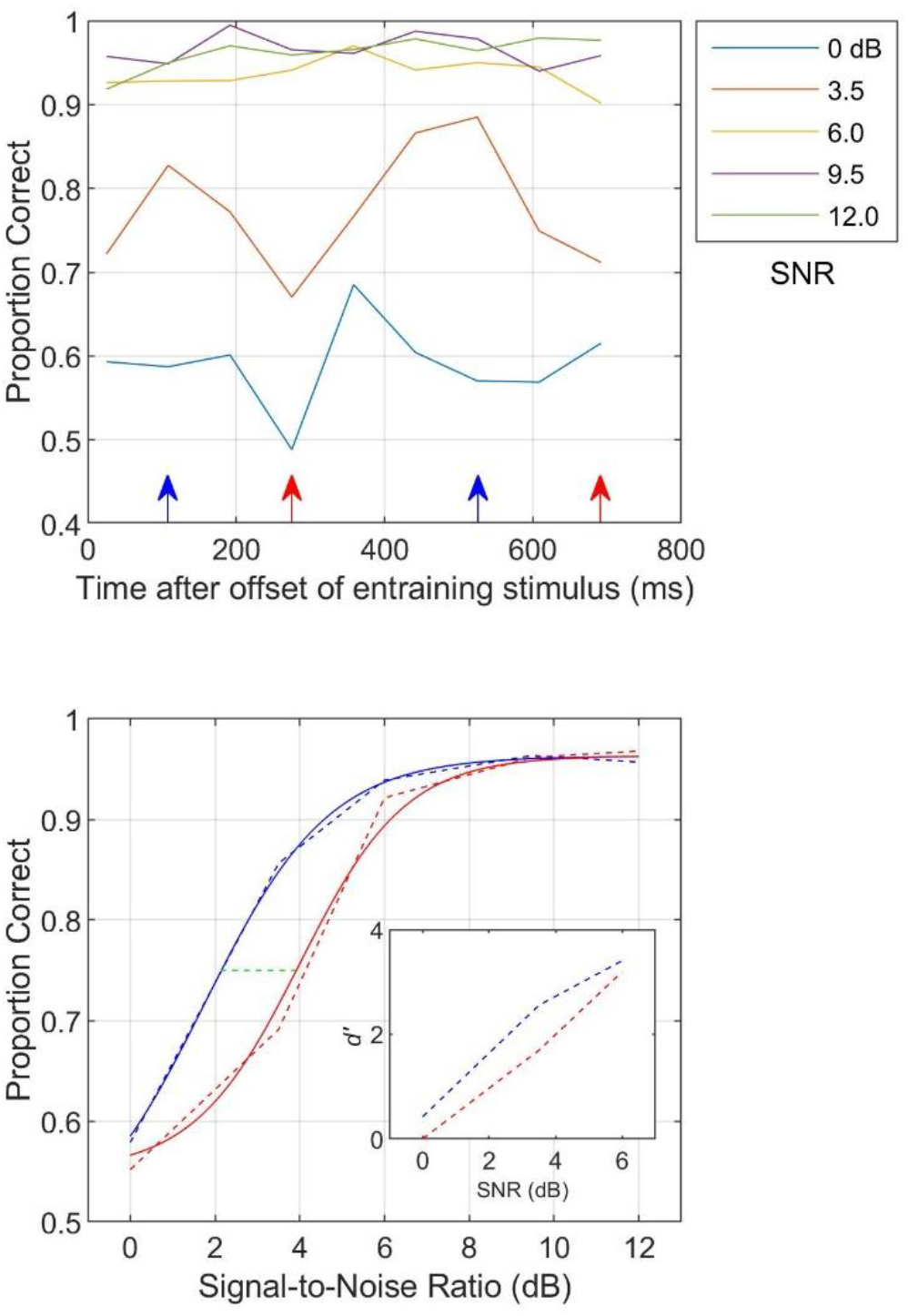
Top: Performance as a function of signal temporal position averaged across subjects. Time zero represents the end of the entraining stimulus. The parameter is SNR, with the 3.5-dB condition showing data reported by Hickok et al. (2015). Bottom: Psychometric functions estimated from data in the top panel (arrows). Red and blue curves shows functions generate from data marked by arrows of the same colors in the top panel. These arrows designate the peaks and dips of the curve at 3.5 dB in the top panel (orange). See text for details.

The bottom panel of Fig. 11 shows a different approach to analyzing forward entrainment using the entire psychometric function. It shows performance as a function of SNR under two conditions. The two conditions are associated with the peaks and troughs of the 3.5-dB SNR function shown in the top panel (orange curve) where the largest modulation in performance was observed (i.e., as reported in Hickok et al., 2015). In order to maximize the number of trials that generated each of two psychometric functions, the data were pooled across the two peaks (marked by blue arrows in the top panel) producing the blue psychometric function in the bottom panel, and the two dips (red arrows) producing the red psychometric function. The dashed lines in the bottom panel represent the actual data and the solid curves are modified logistic psychometric functions fitted to the data. These two psychometric functions, represented by the blue and red curves, show differences in performance between peaks and troughs in performance using a much larger dataset at 5 SNRs. Several interesting trends emerge. For all SNRs, except for the near-ceiling SNR of 12 dB, performance is better for the blue function (associated with peaks) than the red function (dips).

Note that these functions were selected only by virtue of the observed performance at the 3.5dB SNR. The difference in performance at the other SNRs is notably small, in the order of 2 to 3% (compared to an approximately 17% difference at the 3.5-dB SNR). Nonetheless, their direction is consistent with that observed at the 3.5dB SNR. The inset shows the same data plotted as 3-point psychometric functions in *dꞌ* units derived from hit- and false-alarm rates. The two highest SNRs were excluded because of the difficulty in estimating differences in *dꞌ*s near ceiling values without large sample sizes (due to the expanding nature of *p*-to-*z* transforms near unity).

Threshold improvements (difference between blue and red curves) measured at the 75% performance level (green dashed line) is approximately 1.8 dB. This gain in performance, which is estimated from the entire psychometric function, is consistent with the 1.5 dB improvement reported by Lawrance et al. (2014) who also used a signal-in-noise paradigm to compare thresholds for a signal that was either antiphasic or in-phase with a terminated rhythmic noise sequence (see section 1.1).

### 3.4. Effects of experience and intersubject variability

In our recent work we have found what seems to be differences between experienced and inexperienced subjects that manifest largely in intersubject variability and the SNR at which strongest entrainment effects are observed. Figure 12 shows data from 11 subjects who completed the task described by Hickok et al. (2015). Each subject had to detect a tone in steady state noise that followed a modulated stimulus (3 Hz). The data of S1 to S6 were collected recently whereas data from S7 to S11 are those reported previously by Hickok et al. (2015). The first six subjects (S1 to S6) were experimentally naïve (selected from the UCI psychology subject pool) whereas the next 5 (S7 to S11) were highly experienced subjects (graduate students and postdocs) who had participated in a number of other psychoacoustics studies. Of the 5 SNRs tested, the SNR that produced the best bicyclic pattern was selected individually for each subject. This may generally be good practice in order to identify the strongest entrainment effect. We conducted a similar analysis on Sun et al.’s raw data and found that a number of their subjects show stronger entrainment effects at SNRs 1 or 3. Similar to Sun et al. (2021) we observed significant variability in the data of inexperienced subjects and the best SNR at which forward entrainment is detected. In contrast, the performance of experienced subjects is more stable with robust entrainment at an SNR of 2 (3 dB level in Hickok et al., 2015). Similar intersubject variability has also been reported by Jones et al. (2002), Lawrence et al. (2014) and Bauer et al. (2015) where a proportion of their subjects show forward entrainment and a proportion do not under the same experimental conditions. These proportions vary widely across studies. Further evidence of intersubject variability is that in cases where bicyclic patterns are observed, the phases at which a dip (or peak) in behavioral performance is seen do not always precisely line up across subjects. This is not that surprising given the statistical nature of performance and limited sample size, but is important because minor phase misalignments can diminish or flatten modulation patterns in behavioral performance when data are averaged across subjects. Subject-specific phase dependency, as noted earlier, has also been reported for simultaneous entrainment (Henry and Obleser 2012), and a phase drift in the dips and peaks of performance has been reported by Farahbod et al. (2020) in a forward entrainment task as a function of the entraining modulation rate. Variable starting-phase effects has also been reported by Sun et al. (2021) for that proportion of their subjects who they reported as having shown forward entrainment. Supplemental Fig. 8 shows the averaged bicyclic pattern obtained if the individual subject curves of Fig. 12 are shifted by *at most* a single time point in the direction that maximally aligns the dips of these functions. A much stronger M-shaped pattern is observed.

**Figure 12.**
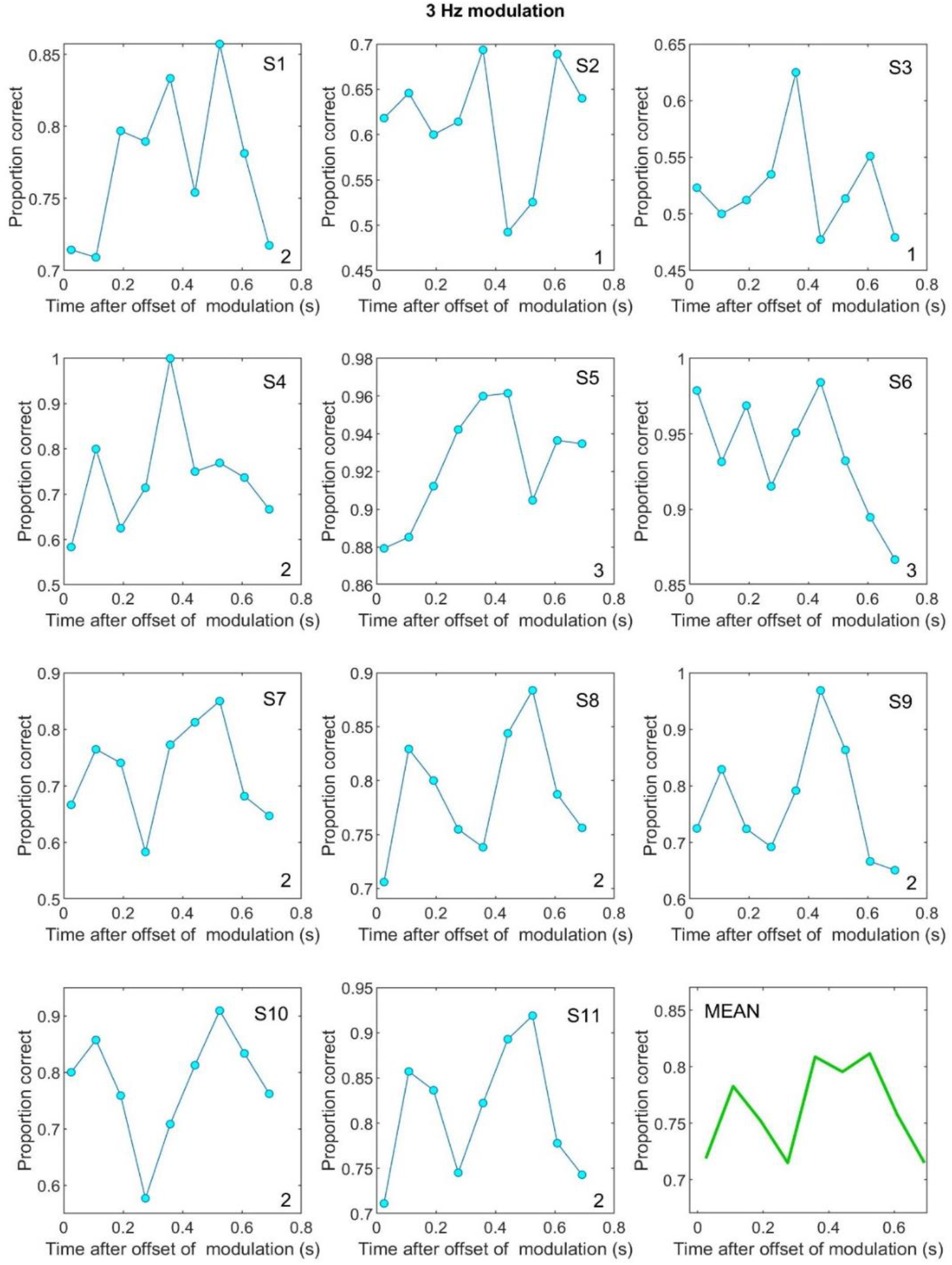
Forward entrainment shown for 11 subjects. S1 to S6 were experimentally naïve (new data), and S7 to S11 were highly experienced (data from Hickok et al., 2015). The number in the lower-right corner of each panel is the SNR (1 to 5) associated with the curve selected individually for each subject as having produced the strongest entrainment effect. Not all subjects show an entrainment effect, e.g., S6.

One reasonable concern that may partially explain large intersubject variability, other than subject experience, might be the number of trials per subject per temporal position used to estimate poststimulus entrainment curves. Both Hickok et al. (2015) and Sun et al. (2021) restricted their analysis to only 1 of 5 SNRs at which performance was simultaneously measured. To address this issue we conducted additional analyses that estimated 3-point psychometric functions using 3 SNRs simultaneously for all 11 subjects from Fig. 12. We measured these functions at each of 9 temporal positions that spanned 2 full cycles of modulation (had the modulation continued). We excluded the two highest SNRs as they were at ceiling performance and therefore uninformative (see Fig. 11). For each psychometric function, we first transformed the individual subject data into detection index *dꞌ* and then averaged these functions across the 11 subjects (similar to the analysis in section 2.3). The resultant 9 psychometric functions (for 9 temporal positions) are shown in the top panel of Fig. 13. The sinusoidal curve in the upper-left side of the panel is a graphical legend that shows the 9 temporal positions at which each psychometric function was measured. Individual subject psychometric functions are shown in Supplemental Fig. 9.

**Figure 13.**
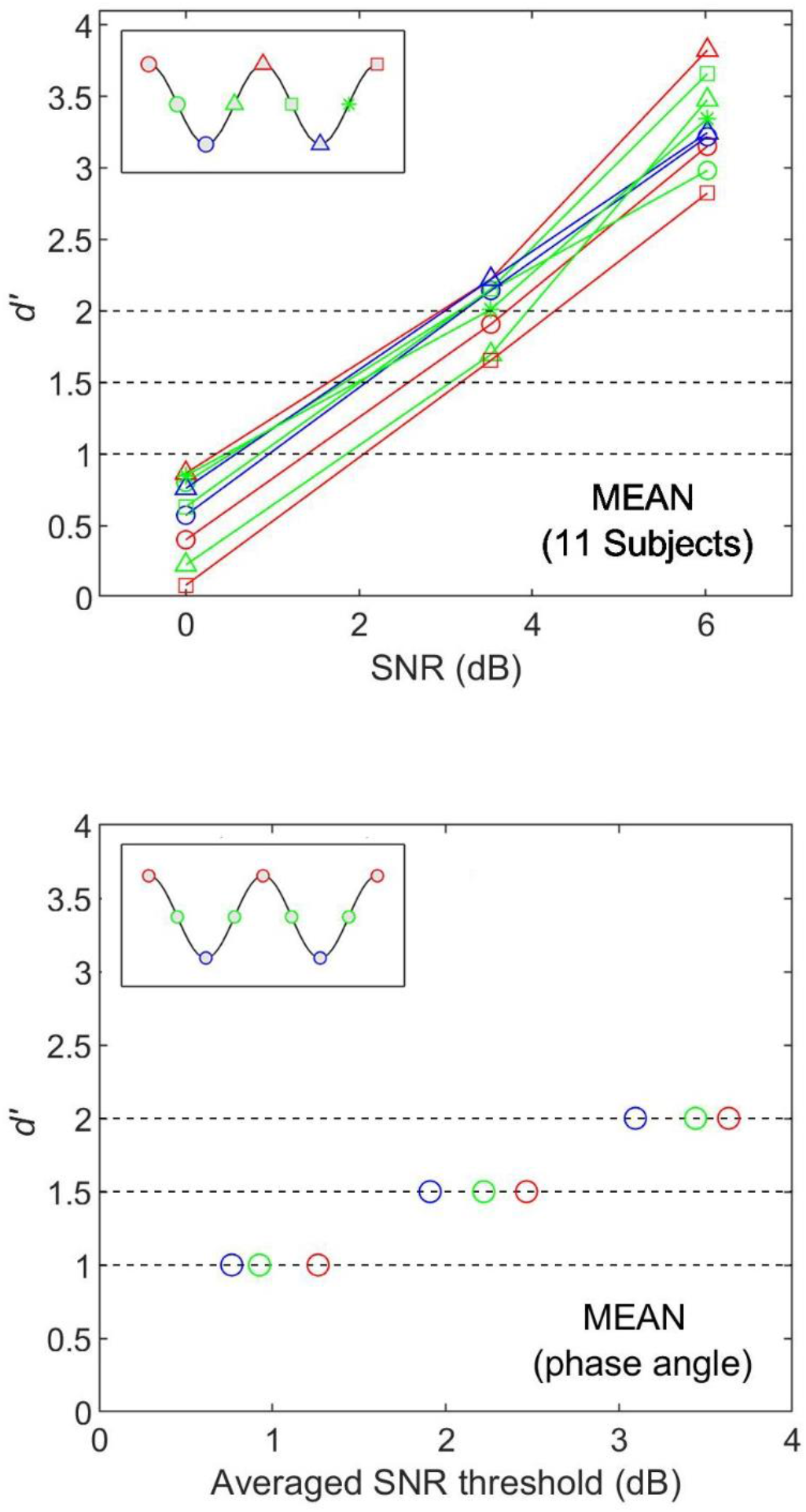
Top: Nine psychometric functions associated with 9 temporal positions shown in the sinusoidal (graphical) legend in the top-left section of the panel (red = peak phase; blue = troughs; green = mid-points). Bottom: SNR thresholds estimated at three different *dꞌ* levels (horizontal dashed lines). Each symbol on each *dꞌ* line is the average of 3 values. For example, the red circle at *dꞌ* = 2 in the bottom panel was obtained from the top panel where the *dꞌ* = 2 line crossed the three red psychometric functions.

Our expectation was that psychometric functions generated from temporal positions at the peaks of the expected modulation curves (red symbols; see legend) would be associated with the poorest performance and those from trough positions (blue) would be associated with best performance, consistent with an antiphasic pattern. We estimated SNR thresholds at three separate *dꞌ* values of 1, 1.5, and 2 (Green and Swets, 1966; Macmillan and Creelman, 2005) represented by the horizontal dashed lines. The predicted SNR threshold for each function was defined as the point at which the *dꞌ* line crossed that function. Hence for each of the 9 psychometric functions (representing the 9 temporal positions) we obtained 3 SNR thresholds defined by the 3 *dꞌ* values. These threshold were then averaged for each phase category (peak=red, trough=blue, and middle=green) yielding 3 SNRs per *dꞌ* as shown in the bottom panel of Fig. 13. Thus, for example, at a *dꞌ* of 2, the three SNR thresholds associated with the red psychometric functions (where the *dꞌ* =2 line crosses the three red psychometric functions in the top panel) were averaged to obtain a single averaged SNR threshold for the peak-phase category (red) at that *dꞌ*. A similar procedure resulted in an averaged SNR threshold for the trough (blue) and midpoint (green) phase categories. The results of this analysis are consistent with an antiphasic pattern of performance in that the peak-phase category produced the poorest thresholds, requiring the highest SNR at a fixed *dꞌ* (farthest to the right on each *dꞌ* line), and the trough temporal positions (blue) produced the best performance, requiring the lowest SNR at any fixed *dꞌ* value, with mid-point positions (green) generating intermediate values as expected. One can argue that *dꞌ* estimates for the peak-phase category are dominated by the extreme end points, i.e., red symbols in the sinusoidal legend at the start and end of the curve, but not middle. This is true, as is also evident from the red-triangle psychometric function in the top panel. However, note that, as shown in Figs. 12 and 8D, while the pattern of performance is antiphasic, i.e, M-shaped, the exact position of the middle dip is on average slightly advanced, often at position 4 instead of 5. The antiphasic *pattern* of performance is further confirmed by the fact that the SNR thresholds for the mid-phase positions (green) fall precisely between the peak-phase (red) and trough positions (blue) at all three *dꞌ* values. These findings, which are based on a significantly larger number of trials compared to those shown in Hickok et al. (2015) are in line with the original results that a tone occurring at approximately a trough temporal position (had the modulation continued) is easiest to detect, followed by mid-phase positions, and lastly the peak-phase positions.

### 3.5 Effects of rhythmic rate

Finally, we have found that forward entrainment is rate limited and lowpass in nature. We tested entrainment for rhythmic rates of 2 to 32 Hz and found that it is strongest when the rate is 2 or 3 Hz, weaker at 5 Hz, and nonexistent for rates from 8 to 32 Hz. Performance for the 2, 3, and 5 Hz were antiphasic to the expected modulation envelope. The absence of poststimulus entrainment at higher rates has also been observed in neural recordings. Lakatos et al. (2013) reported that neural activity continued to oscillate rhythmically for 5 seconds after termination of the entraining sequence (see section 1.3 above) but only for rhythmic rates from 0.8 to 6.2 Hz, and not at the higher rate of 12.2 Hz. This effective range of forward entrainment is approximately the same as that which we have psychophysically observed (Farahbod et al., 2020). These range of rates are also in general agreement with, but somewhat lower than, those reported both for temporal modulation transfer functions (TMTFs) which measure modulation detection thresholds as a function of modulation rate (Viemeister, 1979; Eddins, 1993, 1999; Hsieh and Saberi, 2010; Scott & Humes, 1990; Morimoto et al., 2019), as well as with the firing-rate limits of auditory cortical neuron in response to AM sounds (Joris et al., 2004; Barton et al., 2012).

## 4. Mechanisms

### 4.1. What is entrainment?

In its broadest sense, entrainment in physical systems, and by extension neural systems, occurs when the temporal dynamics of one system is captured by another, resulting in correlated activity beyond chance correlation. More restrictive definitions have been advanced by Haegens and Zion Golumbic (2018) and Obleser and Kayser (2019) that the entrained system be endogenously oscillating at a characteristic frequency (i.e., show natural self-sustained periodicity) and that the entraining system itself be an autonomous oscillator (see also Wilsch et al., 2020). This “coupled oscillators” definition, however describes only a subclass of entrainment phenomena. In applied physics, an entrained system need not be endogenously periodic but can be in a default aperiodic or rest state; similarly, in the case of neural systems, a network can display scale-free or other nonoscillatory activity *prior* to entrainment (He, 2014; Voytek et al., 2015; Schaworonkow and Voytek, 2018; Maniscalco et al., 2018). Some have suggested using the term “neural tracking” or “envelope locking” for this set of phenomena. However, this distinction, while useful (and valid) for explaining certain neural phenomena using certain measurement methods, unnecessarily constrains the definition and fails to conform to broader classifications in physics. The more universal classification scheme, which we favor, includes the entrainment of intrinsically nonoscillatory networks whose computations are often at scales too fine-grained for (and opaque to) extracranial recordings.^21^ Other nonoscillatory entrainment phenomena include stochastic (or noise-induced) entrainment that enhances a system’s nonlinear response to weak or subthreshold signals (Mori and Kai, 2002; Collins et al., 1995, 1996; Wang and Peskin, 2015; Read and Siegel, 1996), aperiodic entrainment that allows irregular neural activity to reliably transmit critical information about external nonperiodic sensory events (Butzin et al., 2015; Mainen and Sejnowski, 1995; Phogat and Parmananda, 2018), chaotic synchronization where systems with close initial conditions in phase space desynchronize and then, counterintuitively, converge via entrainment, to the same trajectory in evolution of their dynamical states (Pecora and Carroll, 2015; Parlitz et al., 1997; Akhmet and Fen, 2015), and fractal entrainment occurring on multiple time scales (Lowen and Teich, 2005; Rhea et al., 2014; Marmelat, 2014). There is significant evidence that auditory nerve firing patterns exhibit such fractal coding whose dimensionality can be modulated aperiodically by environmental input, including potentially by speech and music (Teich, 1989; Teich and Lowen, 1994; Lowen and Teich, 2005).

There are additional aspects of the strict definition that are worth reconsideration. One is the requirement that the entrainment process outlast, in an oscillatory manner, the end of the entraining stimulus (Obleser and Keyser, 2019). We argue that this may or may not be part of an entrainment mechanism (but is not required). The poststimulus decay may be near instantaneous particularly in strongly coupled systems with step-function decay (e.g., electric or laser systems) but also for critically damped (or overdamped) neural systems that do not overshoot (no forward entrainment). Second, the entrainment process need not necessarily be directionally causal (an entraining and entrained system) but rather the coupled systems may be mutually interactive with synchronous activity resulting from bidirectional energy transfer and mode-locking at equilibrium (e.g., interacting networks in the brain that give rise to an intermediate state). This interactive aspect of coupled nonlinear oscillators is, in fact, how Huygens originally defined antiphasic entrainment in physical systems.^22^ Note also that the definition of entrainment as an iterative phase-resetting process in endogenous oscillators is a relatively recent development in neuroscience and less frequent in usage than the broader definition of entrainment in physics. In our view, the proposed narrow (strict) definition aims to promote a particular and valuable perspective about the functional significance of periodic cortical oscillations, but is neither sufficiently comprehensive nor universally established in neuroscience or physics. Even Obleser and Keyser (2019) acknowledge that “the more common term ‘synchronization’ could be used instead” of the term entrainment to describe their narrow definition, and Wilsch et al. (2020), noting the limiting nature of the definition, analyze their data in light of a broader and more nuanced perspective, concluding that they have observed lowpass synchronization but have not found “conclusive evidence” for frequency-specific (narrowband) entrainment as per the strict definition. We therefore favor a broader definition of entrainment, in line with its usage in physics that may or may not include periodic and endogenous oscillators, aperiodic or stochastic systems, neural tracking, resonant coupling, and envelope locking (Eggermont, 1991). From this standpoint, phase resetting is a subtype of physical entrainment that is phenomenologically different than aperiodic, stochastic, or fractal entrainment which also capture the ongoing nonlinear dynamics of an entrained system.

More importantly, and relevant to the current study, the narrow neural definition of entrainment does not naturally extend to psychophysics where the term is used descriptively to represent a wide-ranging set of phenomena in which performance is temporally correlated with a modulating rhythm. Psychophysical studies of entrained performance do not typically take a position on what the specific underlying neural mechanisms might be (e.g., endogenous oscillators) but rather attempt to model behavioral data on a different scale of analysis, i.e., in the context of potential cognitive or perceptual mechanisms that give rise to the observed patterns of performance (e.g., voluntary selective attention, involuntary attentional capture, ringing of modulation filters, listening in the dip strategy at favorable SNRs, symbolic or cognitive cuing, priming, etc.). The questions addressed by psychophysical studies are therefore often quite different than those probed by neural studies of entrainment and direct causal inferences should not be drawn without compelling evidence beyond correlative measures.

### 4.2. Forward entrainment

Forward entrainment describes that part of the entrainment process that outlasts the entraining stimulus. Forward entrainment has been shown using a variety of psychophysical methods (detection, discrimination, and RT designs), with a variety of target signals (tones, noise pulses, temporal gaps or silent intervals), a variety of entraining stimuli (sinusoidal or square-wave modulated noise, triangular or rectangular tone pulse sequences), in different modalities (auditory, visual, tactile; Jones, 2019), across modalities (audiovisual, auditory-motor; Bouvet et al., 2018, 2019, 2020ab), using different neurophysiological techniques (MEG, EEG, ECoG, CSD, and multiunit recordings), and in different species.

How robust is forward entrainment? There are a number of conditions under which forward entrainment fails to be observed. These could potentially be associated with methodological or stimulus design differences. Prior experience, inattention, and intersubject variability may also play a role. Some psychophysical studies that have shown forward entrainment, have also reported the failure of a proportion of their subjects to show the effect under the same experimental conditions (Jones et al., 2002; Lawrence et al., 2014; Bauer et al., 2015). Signal to noise ratio has also been shown to affect the strength of forward entrainment, with weaker or non-existent effects at low or high SNRs. Some studies have shown forward entrainment that lasts for more than one cycle of expected modulation, both behaviorally (Jones et al., 2002; Hickok et al., 2015; Farahbod et al., 2020; Spaak et al., 2014; de Graaf et al., 2013) and neurophysiologically (Lakatos et al. 2013; Spaak et al., 2014; de Graaf et al., 2013), and others have shown an effect that lasts only a single cycle (Forseth et al., 2020; Barnes and Jones, 2000) though the latter have often restricted their measurements to one poststimulus cycle. Both neural and psychophysical studies have shown that forward entrainment dissipates rapidly and is usually nonexistent by the third or fourth cycle after the end of the entraining stimulus. In our view, while the effect has been demonstrated in a large number of studies, it is sensitive to several factors that are not yet fully understood or explored. We have enumerated some of these, but additional studies are warranted to understand which factors (positively or negatively) influence the salience of forward entrainment either behaviorally or neurophysiologically.

### 4.3 Simultaneous vs forward entrainment

Most prior studies have focused on simultaneous entrainment in which the entraining and entrained processes are concurrently active (Henry and Obleser, 2012; Henry et al., 2014; ten Oever et al., 2014; Bauer et al., 2018; for reviews see VanRullen et al., 2011 and Haegens and Zion Golumbic, 2018). The current study is the first review paper to exclusively focus on forward entrainment. Simultaneous and forward entrainment are clearly related but distinct phenomena.^23^ In simultaneous entrainment, phase effects on detection of target signals are more reliable (smaller variance) and do not typically decay with time since the process is reset at every repetition cycle of the entraining stimulus. In fact, in some cases, there is a build-up (instead of decay) of the entrainment effect (Bauer et al., 2018). There are also differences between simultaneous and forward entrainment in measurement of signal predictability. In the latter case, there is no question that entrainment affects processing of future signals locked into a pattern of information change set by the entraining stimulus. This cannot be stated unambiguously in the case of simultaneous entrainment where events to be detected coincide in time with some feature of the ongoing signal. As a consequence, predictive effects cannot be disentangled from ongoing neural processes that potentially include forward and backward masking, comodulation masking release across frequency channels (see below), evoked neural responses, neural inhibition, and a variety of other phenomena that confound interpretation of the entrainment process when the entrained stimulus co-occurs with the entraining stimulus. Furthermore, in simultaneous entrainment paradigms, the entraining pattern need not be fixed but may dynamically vary (as is typical under natural and real-world conditions; see Butzin et al., 2015). The implications for how this affects signal predictability has not been carefully studied. How quickly does the entrained response (neural or behavioral) adapt to new and dynamically changing patterns? Even in the case of fixed modulation rates (and envelope shapes), predictive measurements in simultaneous entrainment are restricted to a single cycle (as contrasted to the sustained activity lasting multiple cycles in forward entrainment). This is because unless the entraining stimulus is dynamically changing in rate or some other physical aspect, one cannot isolate the nonlinear effects of one entrainment cycle from the next in simultaneous entrainment. Forward entrainment, in addition, avoids beats and combination tones produced when two periodic signals (entrained and entraining) are presented simultaneously, particularly for narrowband waveforms with complex harmonic structures (e.g., sinusoidal FM and AM sounds).

Are simultaneous and forward entrainment in signal detection related to simultaneous and forward *masking*, and how do they potentially relate to energetic versus informational masking? Signal detection in stationary noise is primarily, but not exclusively, limited by energetic masking in the passband of auditory filters centered on the signal frequency (i.e., the critical band; Fletcher, 1940; Green and Swets, 1966; Lyon et al., 2010). This is the type of masking that largely limited detection of signals in the steady state part of noise maskers used by Hickok et al. (2015). In addition to energetic masking, however, information derived from the entraining stimulus as to the expected temporal position of masker dips may also have implicitly directed attention to times at which SNR may have been expected to be most favorable. What is facilitated, however, isn’t an *informational* contrast (figure-ground) as is typical in studies of informational unmasking, but rather a process that directs attention. As such, we do not think that the psychophysical patterns of performance in forward entrainment are directly related to informational (un)masking beyond directed attention. Is forward entrainment related to forward (or backward) masking? The detection of a signal in quiet after termination of a masking noise is affected by several factors, including masker level, temporal separation of masker and signal, masker and signal frequency content, and masker and signal duration. Forward masking could be as large as 40 to 50 dB, decays rapidly and linearly as a function of log delay (between end of masker and onset of signal), but could still be as large as 8 to 10 dB at a delay of 100ms (Elliott, 1971; Jesteadt et al., 1982). Interestingly, in tone-on-tone forward masking, the phase of the signal relative to the phase of the masker (had the masker continued) affects the amount of forward masking. When the signal is in phase with the masker, forward masking is approximately 3.5 dB larger than when the signal and masker tones are antiphasic (Jesteadt et al., 1982). This parallels the antiphasic effects that we and others have observed behaviorally in forward entrainment. Therefore, there may be some contribution of temporal masking in entrainment (both forward and simultaneous).

A related finding from auditory psychophysics is that simultaneous masking is more pronounced when a signal occurs soon after masker onset than at later points in a steady state masker (Samoilova, 1959; Zwicker, 1965; Elliott, 1971). Thus, at the very beginning of the steady state noise used by Hickok et al. (2015) the detection of a signal may be more adversely affected because of the terminating cosine phase of the modulating masker that precedes it. This may partially explain the low performance levels observed for the first temporal position at which signal detection was measured (after termination of the modulating noise). However, it cannot explain the continued oscillation in performance. Thus, while temporal masking (forward or backward) may be a contributive factor at early temporal positions, it is not the primary factor in determining the observed psychophysical patterns of performance.

In light of these observations, what do the results of Hickok et al. (2015) mean as they relate to forward entrainment? We think that one of two processes may be involved. Either the modulating noise implicitly draws attention to expected envelope dips or the neural trace of the modulating noise persists after the end of modulation for a fraction of a second (a few cycles) independent of an attentional effect, resulting in improved SNR at antiphasic points in time.

There is increasing evidence supporting an explanation based on attentional mechanisms. Several studies have shown that the entrainment phase dissociates from the entraining stimulus phase. Some studies have shown an antiphasic effect (Hickok et al., 2015; Farahbod et al., 2020; Spaak et al., 2014; Simon and Wallace, 2017) while others have shown an in-phase effect (Jones et al., 2002; Ellias and Jones, 2010; Barnes and Johnston, 2010) or even a subject-dependent entrainment phase (Henry and Obleser, 2012) which may be explained by listening strategy (see section 2.3 in the current paper on how off-frequency listening and different attentional strategies may explain Henry and Obleser’s behavioral data). Further evidence in support of an attentional mechanism is that the effect seems critically dependent on signal-level uncertainty, at least in tone-in-noise detection tasks where switching to a block design eliminates forward entrainment and reintroducing uncertainty recovers the effect in the *same* subjects. The idea that signal uncertainly engages selective attention is consistent with several psychophysical studies in other auditory domains that have traditionally been modeled with attentional filters, for example in tasks involving the detection of tones of uncertain frequency or uncertain duration. Further evidence comes from the large intersubject variability observed in studies of forward entrainment, which may at least partially result from differences in attentional strategies instead of a more rigid bottom-up process. Neurophysiological studies have also shown that neural forward entrainment requires sustained attention to the entraining stimulus (Lakatos et al., 2013).

Finally, we’d like to conclude by noting that there are several well-established psychophysical phenomena that are possibly related to simultaneous and forward entrainment. This link has not previously been made in the literature. These include comodulation masking release (CMR; Hall et al., 1984), comodulation detection differences (CDD; McFadden, 1987; Wright, 1990), and modulation detection interference (MDI; Yost et al., 1989; Hall and Grose, 1991). These processes have been extensively studied in the field of auditory psychophysics and relate to how the modulation pattern in one frequency band affects psychophysical performance in a remote frequency band (several critical bands away) when the modulation envelopes of the band centered on the signal and the spectrally remote band are correlated. It is important to note that conventional theories of signal detection suggest that the detection of a signal (tone) is not affected by noise that is spectrally outside the signal’s critical band. Several across-frequency-channel effects, such as those noted above, violate critical-band predictions for correlated but spectrally distant narrow bands of noise. For example, in CMR, the detection of a signal (tone pulse) in bandlimited noise improves when *additional* noise is presented at a remote frequency band if the two noisebands have correlated envelopes. This process may be interpreted as the capture of the signal-centered noiseband by the remote band, resulting in better isolation of the signal to be detected. CMR has also been observed in a forward-masking paradigm in which the addition of a spectrally remote but correlated noiseband improves the detection of a target tone presented *after* the termination of the making bands (Wright and McFadden, 1987). In CDD, the detection of a near-threshold narrowband *noise signal* is degraded when a remote noiseband whose envelope is correlated with that of the signal band is presented simultaneously. Detection improves when the bands are uncorrelated. In MDI, the detection of the *modulation* of a suprathreshold noiseband is interfered with by the presence of a remote noiseband with the same modulation rate as that of the signal (but, interestingly, not necessarily the same phase; Moore et al., 1991). What these psychophysical phenomena have in common is that the processing of a signal is improved when the noise that limits its detection is “captured” (or entrained) by spectrally distant noisebands with correlated temporal envelopes. From an evolutionary standpoint, such cross-channel entrainment has adaptive value. If a predator’s movement generates correlated modulation in spectrally remote bands, it would be advantageous to encode this activity as a single auditory object instead of as multiple sources with separate spectral identities. During ongoing modulation, top-down signals corresponding to that modulation pattern could generate corollary spectrotemporal predictions within each frequency band. Deviations from those predictions could then be augmented or suppressed as adaptive needs dictate to reduce the entropy of the system’s sensory states (Friston, 2009; Friston et al., 2012; Rao and Ballard, 1999; Clark, 2013). *Forward entrainment may, in this context, instantiate a dynamic auditory afterimage that lasts a fraction of a second to minimize prediction error in signal processing*.

## Acknowledgements

Work supported by NIH R01DC009659 and R01DC03681.

## Data Availability

Raw data are available on request. Video and audio demos referenced in the paper and the Matlab program used to generate Fig. 8 are available in the Supplemental Material section.

**Video demos 1 and 2.** [Video 1](http://www.socsci.uci.edu/~saberi/periodicity/video1.avi) shows the output of a running autocorrelation model of pitch extraction with a 5ms exponential decay in response to a 40-ms FM sound that linearly swept up from 500 to 2000 Hz. Top panel shows normalized autocorrelation amplitude as a function of autocorrelation lag. No pitch estimate is generated by the model (i.e., no secondary peak in the autocorrelation function). The bottom panel shows the response of a GammaTone filterbank to the same sound. The filterbank comprised 50 logarithmically spaced filters whose CFs ranged from 300 to ∼3000 Hz. The vertical red line shows current time. [Video 2](http://www.socsci.uci.edu/~saberi/periodicity/video2.avi) shows the same as video 1 but for a narrower sweep bandwidth from 600 to 900 Hz.

**[Audio demo 1](http://www.socsci.uci.edu/~saberi/periodicity/audio1.wav).** Up sweep followed by down sweep at high frequencies: 40-ms linear FM sweep from 5.5 to 7 kHz (and vice versa). Note that on each trial of Lin et al’s experiment, only a single sweep was presented. Subjects had to judge the direction of the single sweep. The audio sample sequentially plays both types of up and down sweeps for comparison.

**[Audio demo 2](http://www.socsci.uci.edu/~saberi/periodicity/audio2.wav).** Same as audio demo 1 except for a 10-ms FM that linearly swept from 0.5 to 2 kHz. Up sweep followed by down sweep.

**[Audio demo 3](http://www.socsci.uci.edu/~saberi/periodicity/audio3.wav).** Same as audio demo 1 except for a 40-ms FM that linearly swept from 0.5 to 2 kHz. Up sweep followed by down sweep.

**[Audio demo 4](http://www.socsci.uci.edu/~saberi/periodicity/audio4.wav).** Same as audio demo 1 except for a 40-ms FM that linearly swept from 0.6 to 0.9 kHz. Up sweep followed by down sweep.

Video and audio demos are available in the Supplemental Materials Section of this article, as well as for a limited period at: http://www.socsci.uci.edu/~saberi/periodicity/

---

**Supplemental Figure 1.**
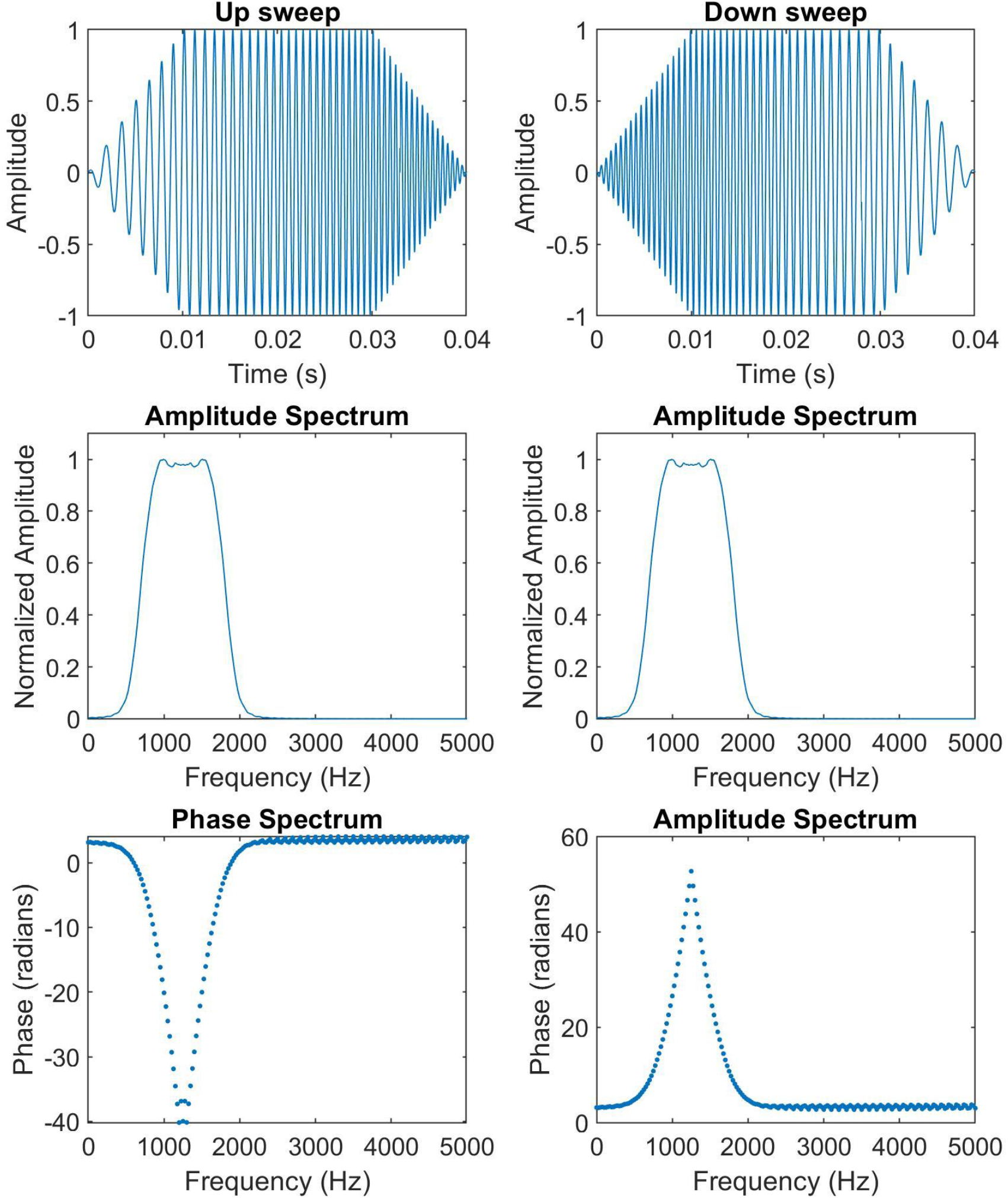
Top panels show the time waveform of 40-ms linear FM up and down sweeps. Middle panels show that their amplitude spectra are the same, and the bottom panels show the differences in their phase spectra.

**Supplemental Figure 2.**
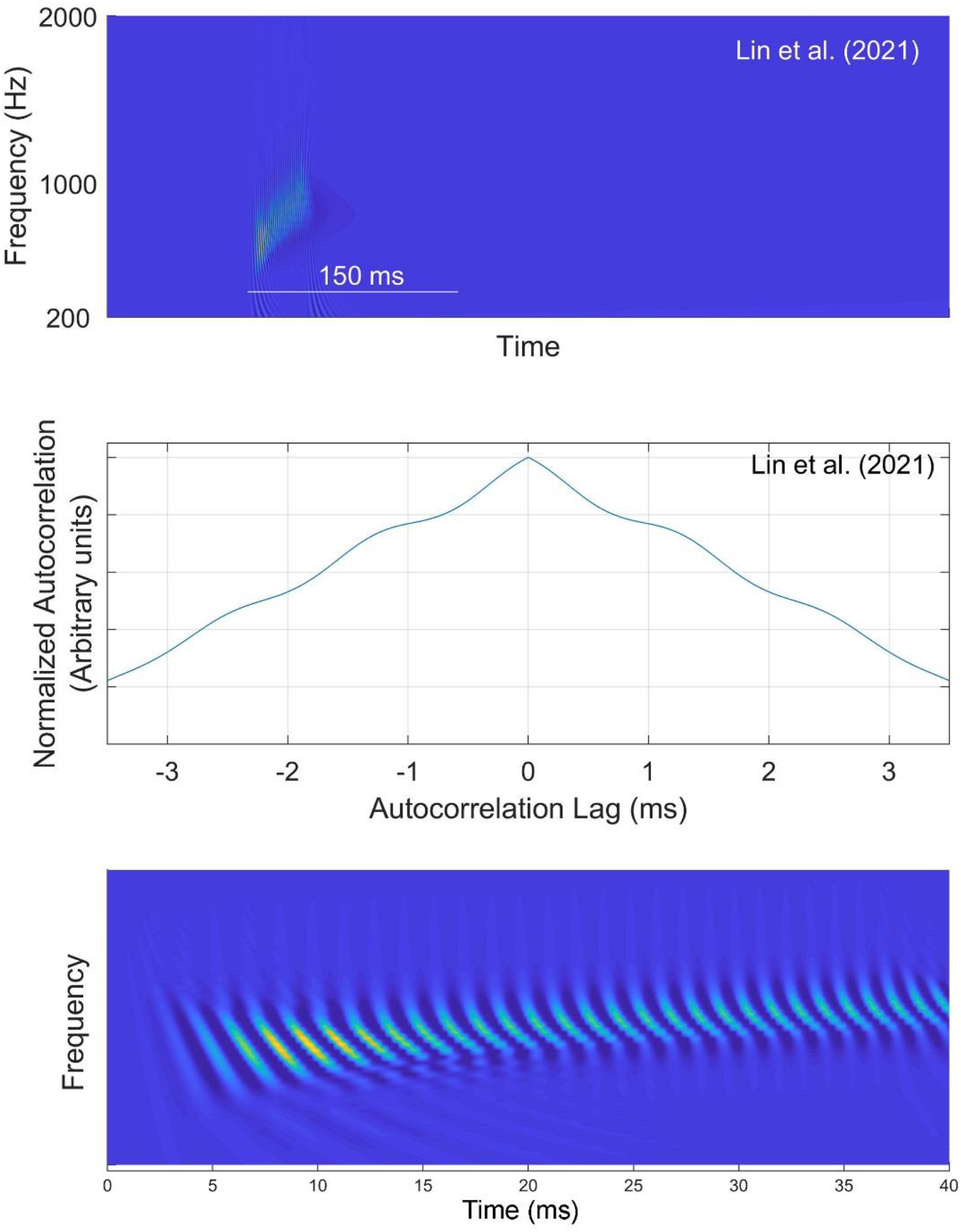
Same as Fig. 5 for a narrower sweep bandwidth (from 600 to 900 Hz). Note that while the curvature (second derivative) of the autocorrelation function shown in the middle panel changes signs, no peak is observed other than at zero lag, resulting in no pitch estimation at the model’s output. The changing curvatures, however, may contribute to a dynamic timbre cue (see also [video 2](http://www.socsci.uci.edu/~saberi/periodicity/video2.avi)).

**Supplemental Figure 3.**
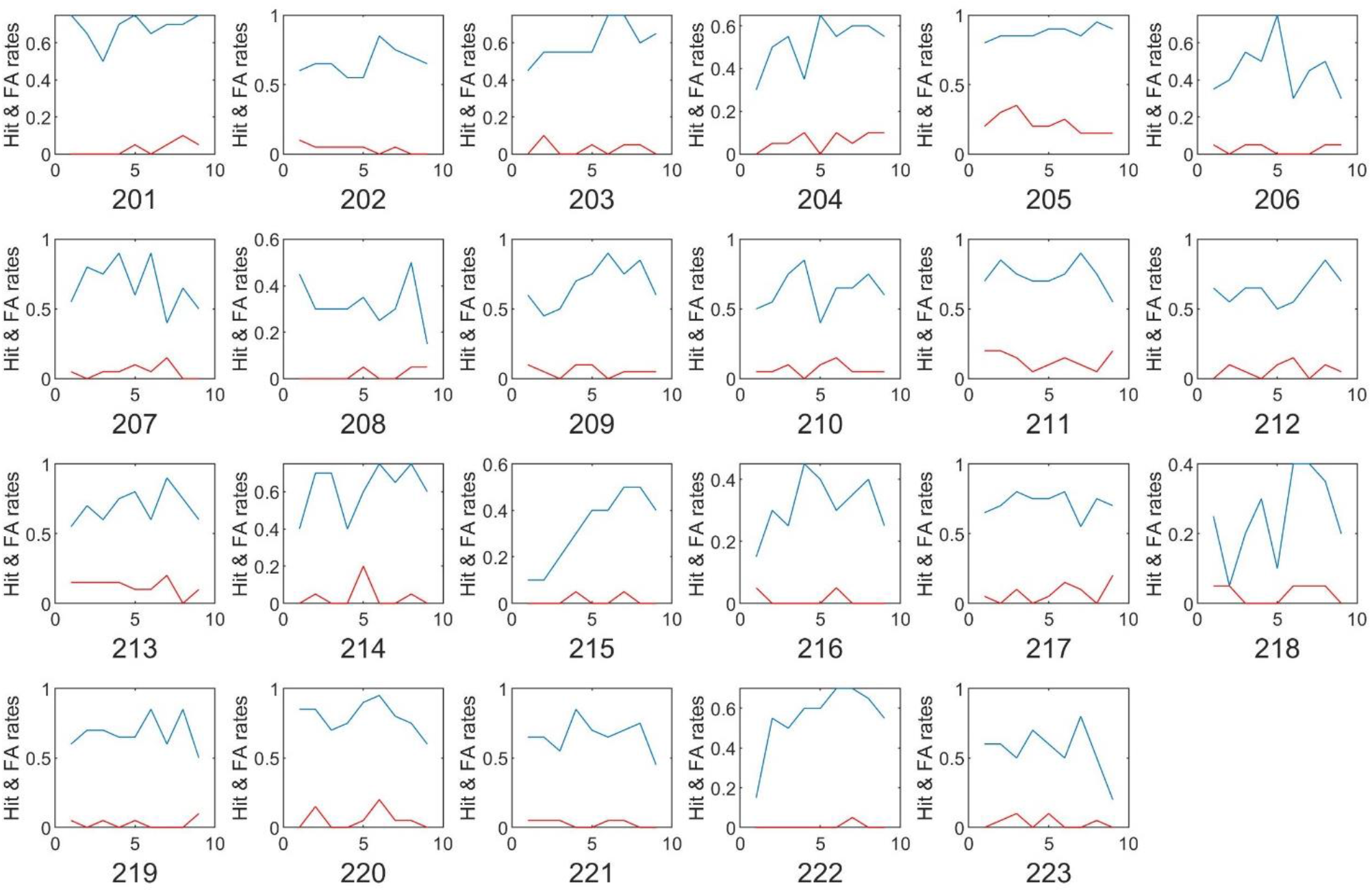
Hit rates (blue) and false-alarm rates (red) for 23 subjects calculated from the raw data of Sun et al. (2021).

**Supplemental Figure 4.**
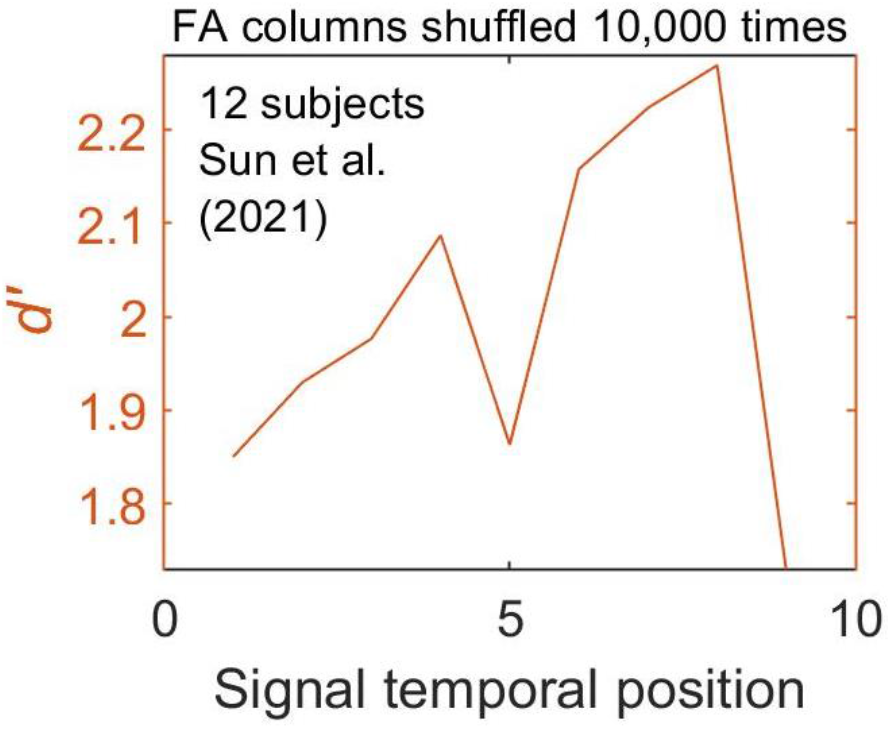
d*ꞌ* curve for 12 of Sun et al.’s subjects who showed strong forward entrainment (see Fig. 8B). For each subject, the *dꞌ* for a given temporal position was calculated from the hit rate for that position and the false-alarm rate randomly selected from any of the 9 temporal positions based on Sun et al.’s labeled data (shuffled false-alarm rates). The curve shown is the average of 10,000 such shuffled curves. See footnote 15 for additional details.

**Supplemental Figure 5.**
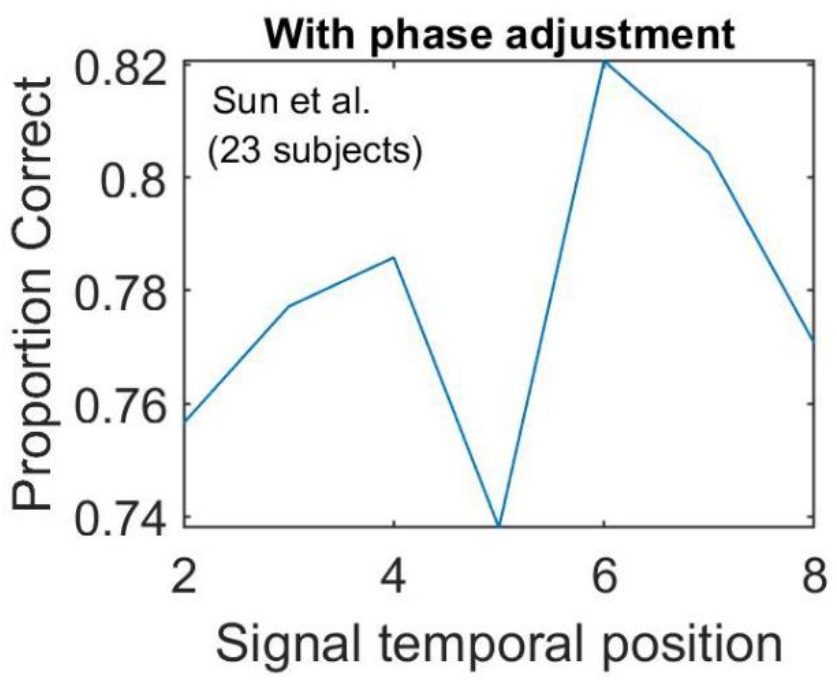
Effects of minor phase alignment: Proportion-correct response curves for each of the 23 subjects from Sun et al. (2021) were shifted by *at most* a single time point (quarter cycle) in the direction that maximally aligned the dips (antiphasic to the expected modulation peak). This figure shows the phase-aligned versions of the same curves that produced the average proportion correct curve in Fig. 8A. See text and footnote 18 for additional information.

**Supplemental Figure 6.**
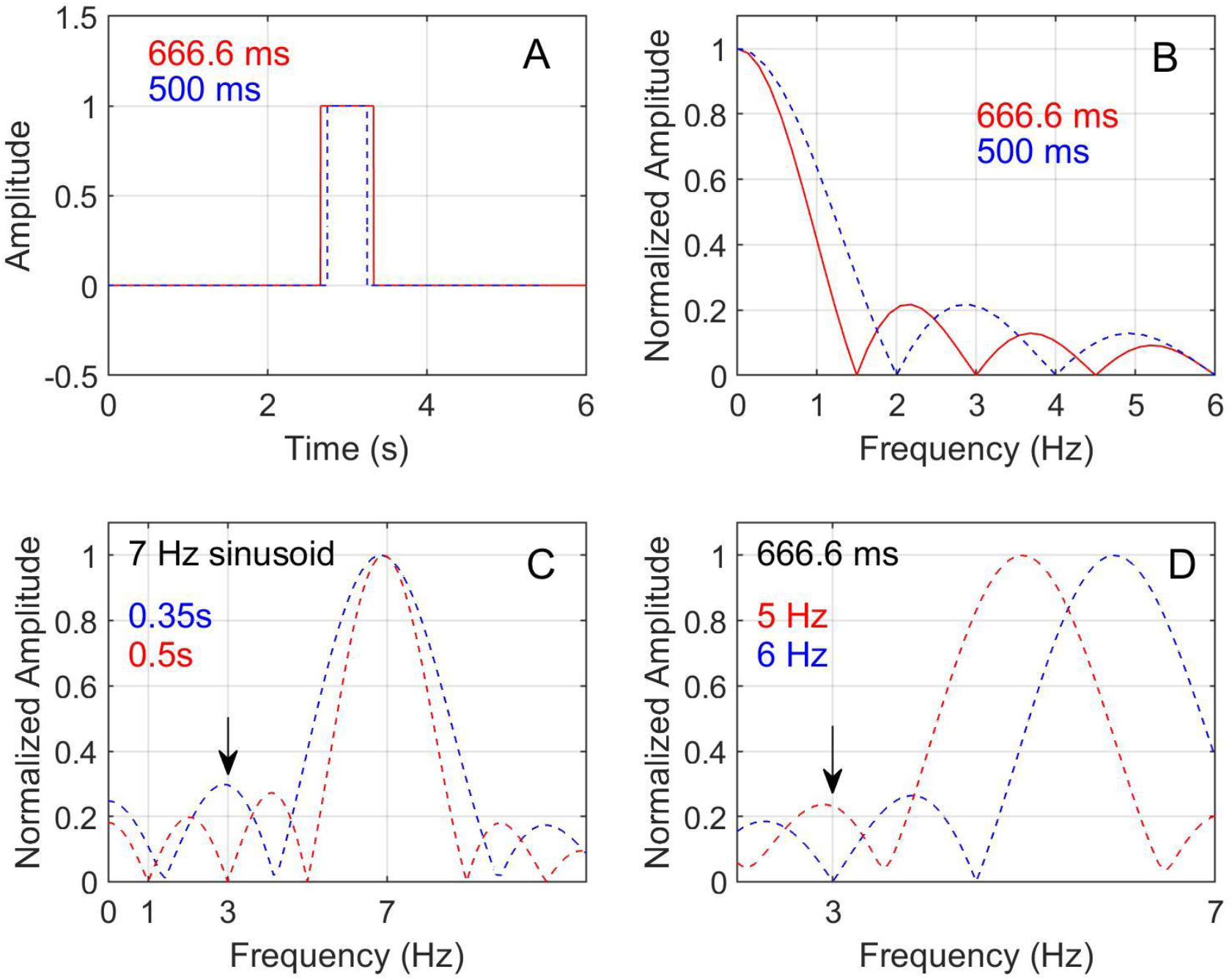
A) Time domain representations of two rectangular windows of differing durations (500 and 666.6 ms). B) The spectrum of each is the sinc function with zero-crossings at the inverse of each window’s duration. C) The spectrum of a sinusoidal pulse with an arbitrary frequency of 7 Hz shown only to demonstrate how false positives can be generated in overestimating entrainment at 3 Hz, or how entrainment strength can be underestimated if the enhanced energy in antiphasic to the frequency of interest, or have no effect if the zero-crossing falls on 3 Hz (note the effect is additive). This is especially a problem when the waveform is very brief and has energy at frequencies that are not of interest. D) For a fixed duration of 666.6 ms, which is equal to 2 cycles of a 3 Hz sinusoid, energy at a number of frequencies (both below and above 3 Hz) can enhance or diminish activity at 3 Hz, resulting in false positives or misses when using “modulation strength” as defined by Sun et al. (2021) to quantify entrainment.

**Supplemental Figure 7.**
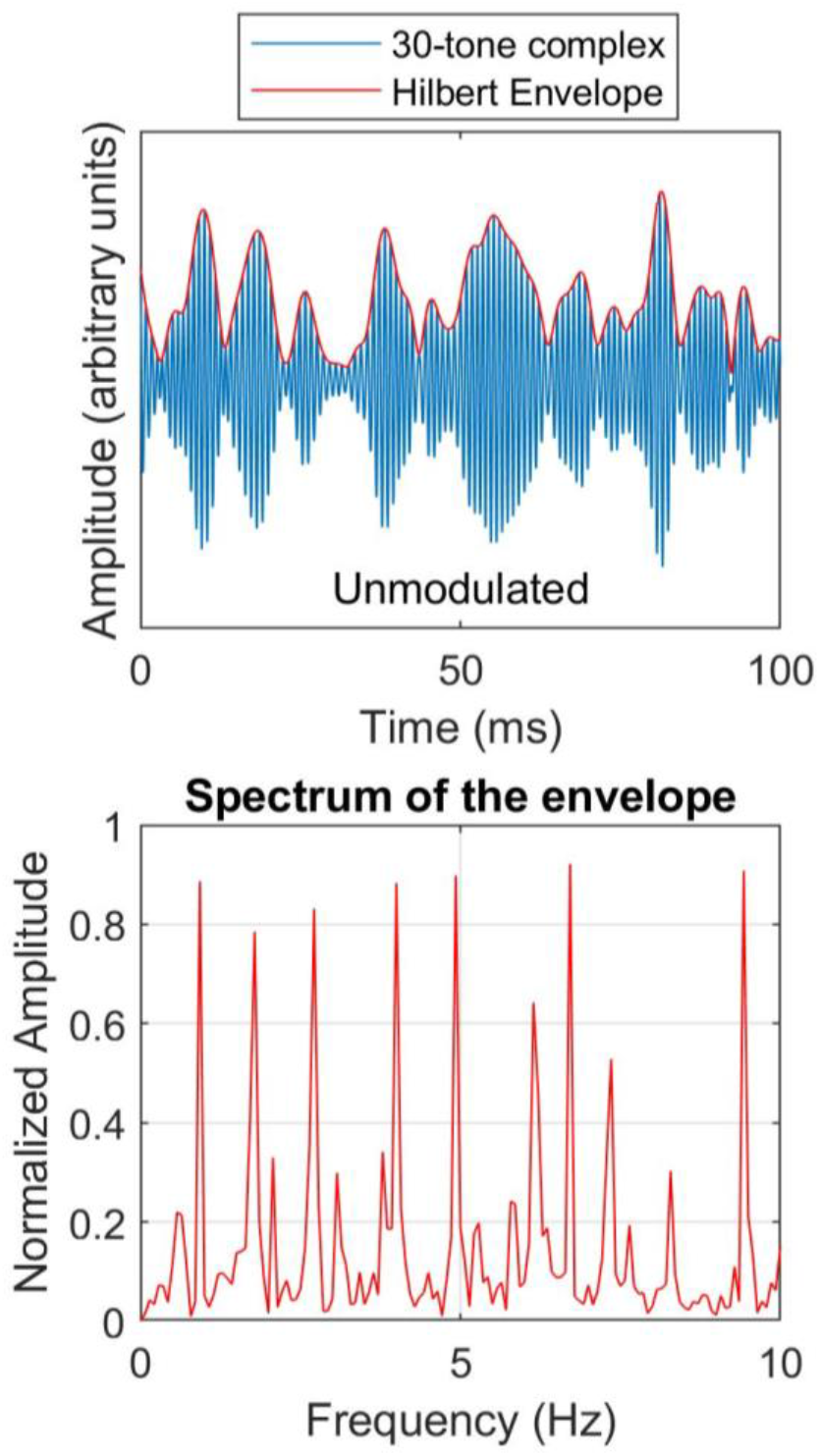
Top panel shows the time waveform of a 30-tone complex (blue) and its envelope (red) calculated from the Hilbert transform of the complex. Bottom panel shows the envelope spectrum, the Fourier transform of the Hilbert envelope (red waveform in the top panel) restricted to below 10 Hz to show the most relevant components. The shape of the envelope spectrum is lowpass with a 3dB (half power) cutoff at approximately 150 Hz. Components below 30 Hz, and especially below 10 Hz, have the largest amplitudes. The total duration of the waveform was 14s (See footnote 20 for details).

**Supplemental Figure 8.**
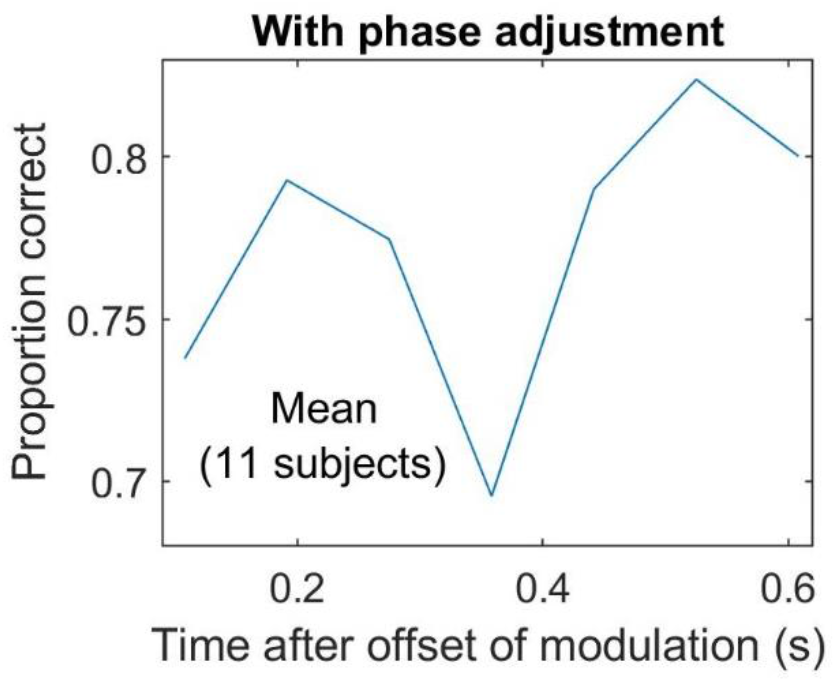
Effects of aligning the curves shown in Fig. 12 by at most 1 time step in the direction that maximally aligned their dips. The curve shows mean performance for 11 subjects after alignment. See also footnote 18.

**Supplemental Figure 9.**
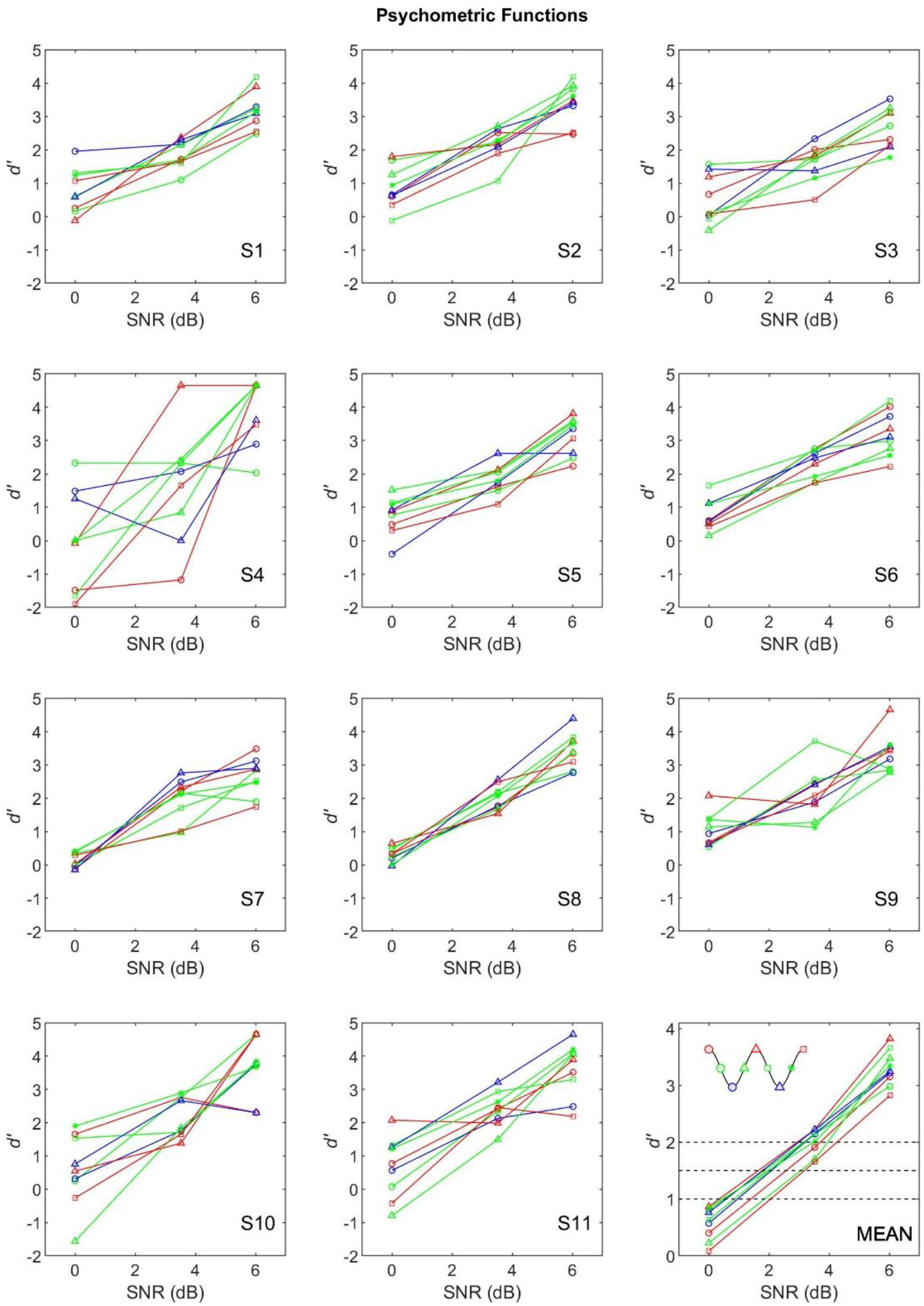
Each panel shows data from one of 11 subjects. Within each panel, 9 psychometric functions are shown for the 9 temporal positions of the signal after the end of the entraining stimulus. The average psychometric functions for these 11 subjects are shown in the lower-right panel and in the top panel of Fig. 13. S4’s functions were based on only 500 trials for reasons unrelated to the experiment, hence the larger than average variance for this subject’s psychometric functions.

## Footnotes

1 Data are reproduced from the original figures using Matlab’s ginput function on figures superimposed on matched axes.

2 The pitch of the first (150ms) tone and the final (60 ms) tone in the driving sequence had the same frequency (unknown to the subject) in order to reduce pitch bias effects. Subjects who reported (in a post-experiment questionnaire) that they realized the standard was repeated were excluded from further analysis.

3 Spaak et al. refer to their measure as a “hit rate” but performance was measured in a 2IFC task where subjects had to indicate whether a target occurred in the left or right hemifield. Thus, their dependent measure was accuracy (proportion correct with chance performance at 0.5) not hit rate in detection-theoretic terms.

4 There are a number of other studies that have shown improvements in RT to signals that follow an auditory sequence, but these often use symbolic cuing, memory-based temporal predictions, or cognitive expectancies (e.g., Stefanics et al., 2010; Breska and Deouell, 2017) that draw voluntary attention to certain aspects of the signal’s timing, and as such fall into a categorically different class of phenomena.

5 Simon and Wallace (2017) also interestingly found that when the pure-tone target was coupled with a brief visual flash (both 50 ms), the cyclical poststimulus power modulation was abolished. Subtracting the auditory only from the audiovisual condition showed that the visual stimulus added its own counteracting (in phase) sinusoidal modulation.

6 Lakatos et al. also found that *ongoing* neural activity (forward entrainment) recorded at two sites in A1 (2 octaves apart) were antiphasic relative to each other. At the best-frequency (BF) site, where the tone frequency matched the BF, the CSD’s negative dip corresponded to the expected time of occurrence of the 25ms tone pulses had these pulses continued, and at the non-BF site, measured simultaneously, the positive peak of CSD activity coincided with the expected time of pulses. This antiphasic effect sharpens responses at the attended BF site. While we considered whether off-frequency listening (Leshowitz and Wightman, 1971; Patterson and Nimmo-Smith, 1980; O’Loughlin and Moore, 1981; Cheatham and Dallos, 1991) may make use of this neural antiphasic activity to improve signal detection, not only are the frequency differences too large (2 octaves) but the antiphasic effect at the off-frequency site has its dip at the *in-phase* time point, not at the trough of the expected modulation envelope, and thus are not likely to be related to the antiphasic behavioral patterns we have observed.

7 Though in one experiment they did observe a near-significant (0.065) trend in favor of the antiphasic target condition in reaction-time measures.

8 In one analysis, they compared *overall* performance differences between the first and second pairs of poststimulus cycles (1-2 vs 3-4), averaged across entrained and “random” conditions. Surprisingly, they did not examine *entraining effects* between these pairs of cycles. It is also worth noting that even in their antiphasic condition, Lin et al. (2021) evaluated performance starting at 1.5 cycles after the termination of the rhythmic cue instead of at 0.5 cycles after cue termination. An optimum selective advantage of an antiphasic condition would have likely been observed at 0.5, not 1.5, cycles after the end of the entraining stimulus. Furthermore, they averaged the antiphasic data for 3 cycles (1.5, 2.5, 3.5) similar to the in-phase averaging confound described above in which performance measures were combined for cycles 1 to 4 after end of the entraining sequence. As noted earlier, this stacks the deck again the possibility of finding forward entrainment which is strongest during the first two cycles and nonexistent by the third and fourth cycles after the end of the entraining stimulus.

9 Only the standard and comparison tones are shown, with the interleaving tones described above in section 1.1 left out for clarity as they do not affect the model’s pitch estimation for the standard and comparison.

10 Neither Lin et al. (2021) nor Wilsch et al. (2020) provide sufficient information about the bandwidth of the FM sweep used other than to say that it was set to generate performance in the range of 65 to 85% correct. Here we have used a reasonable sweep extent of 1.5 kHz (0.5 to 2 kHz), though a lesser or wider bandwidth has no noticeable effect on the current analysis. This can be seen in Supplemental Fig. 2 and [Video 2](http://www.socsci.uci.edu/~saberi/periodicity/video2.avi) which show the model’s output for an FM sound that sweeps from 600 to 900 Hz. This latter range matches that used by Luo et al. (2007) in measuring psychophysical thresholds for brief FM pulses.

11 Similar arguments on use of timbre cues extend to prior work on discrimination of FM pulses (Luo et al., 2007).

12 There are other methodological concerns with Lin et al. (2021) which we do not have sufficient space to discuss. Briefly, two of these are: 1) Use of different stimulus classes for entraining (tones) and entrained (FM) stimuli. Others have typically used within-stimulus-class features for both the entraining and entrained stimuli (e.g., Jones et al. used tones to entrain tones, Hickok et al. used modulated noise to entrain *against* noise to isolate a target tone, Lawrence et al. used entraining noise to entrain noise signals, etc.). A more equivalent design would have been for Lin et al. to use FM pulses or narrowband FM noise carriers as the entraining stimulus. This would have matched the stimulus characteristics of the signal to be entrained. 2) Use of a block design in which rhythmic and random targets were never mixed within a run. This significantly diminishes uncertainty effects that have been shown to be critical to forward entrainment. Jones et al. (2002), Hickok et al., (2015), Farahbod et al. (2020), and many others introduce such uncertainty. A block design also allowed use of offset cues in the steady state continuous tone (what they refer to as “random cue”) as a marker to predict rhythmic target timing. This offset time consistently and deterministically marked the beginning of the target observation interval, providing implicit temporal information over the course of a block of trials as to when a rhythmic target may occur. The one notable exception, as described above, was that on a small proportion of rhythmic cue trials (20%), an antiphasic target was presented instead of a rhythmic target (80% of trials). This comparison is likely the most reasonable experimental contrast in attempting to find an entrainment effect. In spite of the low number of trials, Lin et al. do report a near significant difference (p=0.065) in reaction times between the in-phase and antiphasic conditions when the cue was rhythmic (at least in Experiment 3). Nonetheless, in the majority of cases, Lin et al.’s use of a block design provided cues as to rhythmic target conditions even in the “random” (continuous tone) cue condition, diminishing potential differences between cue types.

13 Some of the methodological differences between Hickok et al. (2015) and Sun et al. (2021) include the following. Sun et al. used a staircase procedure to set signal levels individually for each subject (experiment 1), used practice runs prior to the actual experiments that exposed subjects to high SNR levels, participants were restricted to a response time window of 1.5s, intertrial interval was randomized (between 1 and 2s), and a pseudorandomized procedure was used in assignment of trial types instead of a fully randomized design used by Hickok et al. (2015). In their pseudorandomized design, they forced an equal number of trials for signal and no-signal conditions (as well as an equal number of trials per temporal position and signal level). Psychophysicists typically avoid use of pseudorandom designs largely because it introduces serial dependencies near the end of runs that can affect subject response strategy. An examination of the raw data of Sun et al. (2021), which they made publicly available, shows such non-independence resulted in aggregation of specific trial types, for example “signal present” trials, at the end of many run, e.g., subject 205, run #18; subject 207, run #13; and subject 213, run #7. This serial dependency can, for example, increase the likelihood of false alarms on runs in which the prior expectancy has changed in favor of responding “yes” near the end of a run (i.e., a decision criterion shift). Subjects can pick up on these trends and develop effective response strategies (same argument applies to signal level and temporal positions). The authors state that balancing of trial types was “particularly crucial to prevent participants from developing response biases”. We argue that in fact such pseudorandomization almost ensures development of response biases within a run, and especially near the end of a run.

14 In the following analysis we focus exclusively on their experiment 2 (“exact” replication). Sun et al. (2021) have made their raw data publicly available at: https://dx.doi.org/10.17617/3.5c

15 *dꞌ* = Z_hits_ - Z_false alarms_ . By convention we assumed a small inattention rate in setting a ceiling hit rate of 0.99 and a floor false-alarm rate of 0.01 (Green, 1995; Saberi and Green, 1997; Macmillan and Creelman, 2005). This avoids the practical problem of *dꞌ* = ∞ when estimating performance in small samples. For each combination of temporal position and signal level, false-alarm was determined from the raw data of Sun et al. which for *each* trial included a designation of signal temporal position and level, as well as whether or not a signal was presented on that trial. While all no-signal trials are by definition the same, false-alarms may be independently calculated for each time-level combination from the designated labels for temporal position and level on each trial *prior* to the final decision on whether a signal should be presented. Using a single *averaged* false-alarm rate from all “no-signal” trials of a subject to calculate *dꞌ* for different temporal positions is not ideal as it will introduce partial correlation across temporal positions and diminish differences in performance across these temporal positions. We also considered what may happen if in calculating false-alarm rates we replaced Sun et al.’s labeled trials with shuffled false-alarm rates across temporal positions while maintaining independent estimates at each position (i.e., calculating *dꞌ* for one temporal position using the hit rate for that position and a false-alarm rate randomly selected from another temporal position using Sun et al.s labels). While the full-subject set of curves were variable as expected given the reported large intersubject variability (with many, but not all, permutations showing M-shaped patterns), the *dꞌ* curve from the 12 subjects who showed strong forward entrainment (Fig. 8B) were extremely stable (consistently M-shaped patterns). Supplemental Fig. 4 shows the averaged *dꞌ* curve for these subjects obtained from 10,000 permutations of false-alarm rates across temporal positions. The bicyclic pattern is clearly maintained in spite of shuffled false-alarm rates. This shuffling approach, of course, assumes that false-alarm rates are derived from stationary internal noise across temporal positions. If the internal noise that limits performance is nonstationary, with its variance modulating in-phase with the terminated entraining stimulus (as some neurophysiological findings suggest, Lakatos et al., 2013), and if the listener attends to a specific temporal position in making a response decision, then the false-alarm rate may also modulate accordingly as a function of temporal position. However, any such nonstationarity, if it exists, would only serve to enhance bicyclic patterns of performance, i.e., random assignment of “no-signal” trials to a specific temporal position in calculating *dꞌ* would only wash out any *real* modulatory effect in false-alarms, resulting in a reduced likelihood of observing a bicyclic pattern. In other words, random assignment of “no-signal” trials (or use of Sun et al.’s labels for such trials) works against our hypothesis (in favor of the null) and the fact that we still observe bicyclic patterns serves to confirm forward entrainment in signal-detection.

16 Subject numbers 202, 203, 208, 210, 211, 212, 214, 216, 218, 219, 221, and 223 from Sun et al. (2021).

17 In their original manuscript which Sun et al. graciously shared with us, only hit-rate measurements were made. The peer review history of their paper, made publically available by the European Journal of Neuroscience (https://publons.com/publon/10.1111/ejn.15367) shows that the reviewers cautioned against this approach. In spite of these cautions the authors maintained hit-rates as the basis of their primary analysis, largely dismissing use of false alarms in calculating *dꞌ* as just adding noise to the data. In the final version of their paper, the authors, to their credit, do show secondary measurements using *dꞌ*s, but this analysis is incomplete, failing to take the extra step of reporting the averaged *dꞌ* curve that shows a bicyclic pattern (our Fig. 8) and failing to conduct the necessary statistical analyses that would have confirmed an antiphasic dip in performance (see our critique of their “modulation strength” measurements below).

18 The phase alignment effects in Supplemental Fig. 5 is just for illustration purposes as we found it to be instructive. Obviously aligning dips of curves across subjects, even if shifted by only one time point, will by definition generate bicyclic peaks adjacent to the dip. Not all individual subject curve-shifts resulted in precise alignment of dips, especially if a minimum was two or more time points away. Because of shifting curves to the left or right, time points 1 and 9 were excluded from this figure (Suppl. Fig. 5) since the data at these points were based on fewer than 23 subjects. We should also note that estimating the exact phase of a signal relative to the modulation envelope is complicated because of the duration of the signal (50ms) relative to that of the quarter cycle distance between time points at which performance at was measured (83 ms). Hickok et al. plotted their data displaced by 25ms to designate the center of the tone signal as a nominal point for comparison. Sun et al. uses the onset of the tone as an anchor. Because the tone signal is not an impulse with an infinitely narrow duration, some uncertainty is introduced as to the precision with which one may estimate the exact phase of the signal relative to that of the modulating noise at which best performance is observed. This uncertainty is amplified if there is subject-specific variability in entrainment phase as previously reported by Henry and Obleser (2012). The small quarter-cycle (1 time point) alignment that enhances bicyclic patterns of performance may also have relevance to the results of Forseth et al. (2020) who, using the same stimuli as Hickok et al. (2015), found best performance at a modulation phase that was a quarter-cycle away from that reported by the latter study (i.e., at the rising slope instead of dip of the expected modulation). Sun et al. inaccurately state that Forseth et al. found best performance that “seemed to occur in-phase with the preceding modulation.” The optimum time point reported by Forseth et al. was in fact midway between the peak and dip temporal positions.

19 Note that the duration of the tonal signal (50ms) is equivalent to 0.15 cycles of modulation at 3 Hz, while the average difference in starting phases of performance curves between Hickok et al. (2015) and Sun et al. (2021) is 42 degrees, or 0.12 cycles at 3 Hz, less than the range of phases covered by the duration span of the 50ms tone.

20 Even though the study by Henry et al. (2014) is related to simultaneous (not forward) entrainment, because it is both interesting and widely cited, we would like to briefly comment on the structure of their stimuli which consisted of a 30-tone complex simultaneously modulated both in frequency (FM = 3.1 Hz) and amplitude (AM = ∼5 Hz). Each component of the complex was randomly selected on each trial from a range of 500 Hz, centered on a carrier that itself was randomly selected on each trial from the range 800 to 1200 Hz (resulting in an overall range of 550 to 1450 Hz across trials). The level of each component linearly declined with increasing distance from the center frequency. It was this 30-tone complex that was simultaneously frequency and amplitude modulated *en masse*. There are three points we would like to make about this stimulus structure. First, the 30-tone complex itself, *prior* to the addition of any FM or AM structure, contained significant envelope amplitude modulation as shown in Suppl. Fig. 7, including envelope energy near frequencies of 3 and 5 Hz. No information is provided about the slope of the linear function that attenuated the component levels; Suppl. Fig. 7 shows the most severe case where the levels attenuate to zero at the edges of the frequency range. However, we also examined the case where the edge components attenuated to the half-amplitude point and found similar though not as pronounced lowpass modulation patterns in the *unmodulated* complex (i.e., prior to adding AM or FM to the carrier). The shallower the slope of the attenuation function, the less of an issue is the presence of such cues resulting primarily from acoustic “beats” of components adjacent to the center frequency. Second, as noted in section 2.3, there is an FM-to-AM conversion from cochlear filtering in the auditory periphery, with opposite phases of the induced-AM at the output of filters above and below the FM carrier. Third, the spectrum of a sinusoidal AM comprises 3 frequency components, the carrier and two sidebands that are in-phase with the carrier and spaced equidistant above and below the carrier (the spacing is equal to the modulation rate). The spectrum of a sinusoidal FM is made up of the carrier and an infinite number of sidebands symmetric above and below the carrier, with the lower odd harmonics inverted in phase relative to the carrier. The amplitudes of these sidebands decline as Bessel functions of order n (sideband number) (Inglis, 1998; Carlson, 1986). These sidebands interact with nearby tones of the 30-tone complex and potentially generate beats or cancel components if they are of the same frequency but opposite in phase.

21 We are making two points here, first that many (likely most) local cortical networks are not intrinsically oscillatory and their computations are not characterized by simple sinusoids but rather require much more mathematically complex descriptions, and second because of technical limitations in measurement methodology, their critical role is often missed (or undervalued) in analyses that depends exclusively on extracranial recordings. As the late computational neuroscientist Walter Freeman (2000) notes, the problem is equivalent to trying to figure out how a car engine works by using a stethoscope.

22 Christiaan Huygens invented the pendulum clock to improve maritime navigation. The clock’s pendulum motion allowed robust measurements that could withstand the rigors of sea travel. He conducted tests on *pairs* of sea clocks for practical redundancy in case one of the clocks stopped or had to be cleaned during travel. Huygens inadvertently physically coupled two clocks by hanging them from the same imperceptibly loose wooden beam and noticed that within half an hour they became synchronized. In a 1665 letter to Sir Robert Moray, read to the Royal Society, Huygens’ discovery is described as “an odd kind of sympathy perceived by him in these watches suspended by the side of each other.” (Bennett et al., 2002). This observation appears to be the first known scientific demonstration of what today we call “entrainment” in a physical system, though the actual term “entrainment” first appears later in the scientific literature, e.g., in the field of relativistic physics (Laue, 1907; von Hirsch, 1908). Interestingly, in a letter to his father Huygens notes that while confined at home with a brief illness, he observed that paired clocks *always* swung 180 degrees out of phase when synchronized. More recent experiments have demonstrated that a small oscillation frequency difference between the two clocks enhances antiphasic synchronization (Czolczynski et al., 2010; Willms, 2017; Yang et al., 2018). The resultant oscillation frequency of the coupled system is between the oscillation frequencies of the two clocks and does not match one or the other as is the case with neural or psychophysical studies of entrainment using an external driving modulator that directionally captures the behavior of the entrained system. Lakatos et al. (2019) has proposed restricting the definition of entrainment to a particular type of neural phenomenon in which a rhythmic stimulus directionally and repetitively resets the phase of an endogenous neural oscillator, distinguishing between this phenomenon and transient phase resetting resulting from a single external event, and bidirectional effects in coupled oscillators described above. This definition may have practical and theoretical advantages when analyzing certain types of neural data. However, consistent with its use in the physical sciences, we prefer the more widely used (and broader) definition which facilitates analysis across disciplines.

23 While one may consider forward entrainment to be an extension of simultaneous entrainment (or even the same process) it is instructive to consider their differences in what they reveal about the processes involved in both neural and psychophysical entrainment. Similar contrasts between forward and simultaneous making have provided important insights into the processes that underlie auditory masking.

