## Supplemental AV and Figure files for "Forward Entrainment: Evidence, Controversies, Constraints, and Mechanisms": PlotFig8.m

```
clear;
clc;
close all;

load('sundata'); % From experiment 2
x=sundata; % 41400 rows by 8 columns
% Each subject's data covers 1800 rows

% Columns:
% 1) Subj # (201 to 223 - 23 subjects in Expt 2)
% 2) Trial # (1 to 1800 per subject)
% 3) Trial # in a block (1 to 90)
% 4) SNR (1 to 5)
% 5) Tone presence (0 or 1)
% 6) Tone position (1 to 9)
% 7) Resp (0=no sig, 1=sig)
% 8) Correct/Incorrct (1 or 0)

snr=2; % Use SNR 2

for ss=1:23;

snum=200+ss;
a=find(x(:,1)==snum);s=x(a,:);  %find all data for this ss
a=find(s(:,4)==snr);s2=s(a,:);  %just keep this SNR

    for tau=1:9  %9 temporal positions
    a=find(s2(:,6)==tau);  %indexes for all trials for this temporal position tau (1 to 9)
    s2tau=s2(a,:); % All trials for this temporal position tau (1 to 9)
    a=find(s2tau(:,5)==1); % Just target trials
    hits(tau)=sum(s2tau(a,8))/length(s2tau(a,8));  % Hits
    a=find(s2tau(:,5)==0); % Just No-target trials
    fa(tau)=sum(s2tau(a,7))/length(s2tau(a,7));  % False Alarms
     
     if hits(tau)>0.99;hits(tau)=0.99;end;  % Assume 2 percent inattention rate
     if hits(tau)<0.01;hits(tau)=0.01;end;
     if fa(tau)>0.99;fa(tau)=0.99;end;
     if fa(tau)<0.01;fa(tau)=0.01;end;
    dprime(tau)=norminv(hits(tau))-norminv(fa(tau));
    pc(tau)=sum(s2tau(:,8))/length(s2tau(:,8));
         
    end; %for tau
     
allhits(ss,:)=hits; %keep track of hit rate for every subject
allfa(ss,:)=fa; %keep track of false alarm rate for every subject
allpc(ss,:)=pc; %keep track of proportion correct for every subject
alldp(ss,:)=dprime; %keep track of d prime for every subject

end; %for ss

subplot(3,2,1);
h=plotyy([1:9],mean(allpc),[1:9],mean(alldp)); 
xlabel('Signal temporal position');
ylabel(h(1),'Proportion Correct')
ylabel(h(2),'d''','fontsize',11,'fontangle','italic');
text(0.5,.8,{'23 subjects';'Sun et al.';'(2021)'},'fontsize',8);
text(8.9,.81,'A');
ylim([0.73 0.82]);
set(h(2),'ylim',[1.8 2.4],'ytick',[1.8 2 2.2 2.4]);
set(h(1),'ytick',[0.73 0.75 0.77 0.79 0.81]);

subplot(3,2,3);  %12 subjects from Expt. 2 of Sun et al. 
aa=[202 203 208 210 211 212 214 216 218 219 221 223]'-200;  %Subtract 200 to get row #
somepc=allpc(aa,:);
somedp=alldp(aa,:);
h=plotyy([1:9],mean(somepc),[1:9],mean(somedp)); 
xlabel('Signal temporal position');
ylabel(h(1),'Proportion Correct')
ylabel(h(2),'d''','fontsize',11,'fontangle','italic');
text(0.46,.795,{'12 subjects';'Sun et al.';'(2021)'},'fontsize',8);
text(8.9,0.805,'B');
set(h(2),'ytick',[1.5 2 2.5]);

subplot(3,2,5);
%From Hickok et al. 2015, their Fig. 3 dprime and PC data
%Raw data from Hickok et al. 2015 are available in the 
%data repository referenced in Farahbod et al. (2020)
hickokdp=[2.3100    2.3881    2.0046    1.4495    1.8092    2.4505    2.7189    2.1636    1.9355]; 
hickokpc=[0.7217    0.8275    0.7720    0.6705    0.7666    0.8661    0.8851    0.7494    0.7118];
h=plotyy([1:9],hickokpc,[1:9],hickokdp);
xlabel('Signal temporal position');
ylabel(h(1),'Proportion Correct');
ylabel(h(2),'d''','fontsize',11,'fontangle','italic');
text(0.4,0.94,{'Hickok et al.','(2015)'},'fontsize',8);
text(8.9,0.96,'C');

set(1,'pos',[85 85 490 590]);
```
