## Supplemental AV and Figure files for "Forward Entrainment: Evidence, Controversies, Constraints, and Mechanisms": readme.docx

The Matlab program “PlotFig8” will use data in file sundata.mat to generate averaged *dꞌ* and proportion correct curves shown in Fig. 8.

The data file contains 41400 rows by 8 columns.

Each subject’s data is contained in 1800 rows (1800x23=41400)

Columns represent:

1) Subject # (201 to 223)

2) Trial # (1 to 1800 per subject)

3) Trial # in a block (1 to 90)

4) SNR (1 to 5)

5) Tone presence (0 or 1)

6) Tone position (1 to 9)

7) Response (0=no signal, 1=signal)

8) Correct/Incorrect (1 or 0)
